# Collective biosynthesis of plant spirooxindole alkaloids through enzyme discovery and engineering

**DOI:** 10.1101/2025.01.28.635216

**Authors:** Danni Chu, Hongkui Wang, Zhiwen Nie, Kang-li Li, Jiaqing Cao, Meijing Yang, Qin Yin, Yang Gu, Yindi Jiang

## Abstract

Collective synthesis is a well-established strategy to produce natural product collections from common intermediates. Extending this concept to enzymatic systems, collective biosynthesis offers a sustainable approach to diverse molecular collections but faces distinct challenges, including incomplete pathway knowledge and narrow substrate specificity of biosynthetic enzymes. Here, we overcome these limitations through enzyme discovery and protein engineering to achieve collective biosynthesis of spirooxindole alkaloids, an important class of natural products with diverse biological activities. We identified two key enzymes that catalyze enzymatic epimerization, working sequentially with a previously discovered cytochrome P450 enzyme to fully elucidate the biosynthetic pathway of pentacyclic spirooxindole alkaloids. Structure-guided engineering expanded substrate scope, enabling collective biosynthesis of 12 tetracyclic and pentacyclic spirooxindole natural products. We also generated new-to-nature fluorinated and deuterated derivatives through precursor-directed biosynthesis. This work provides crucial insights into plant spirooxindole alkaloid biosynthesis while establishing a powerful approach for sustainable production of complex natural products.

## Introduction

Spirooxindole alkaloids, a significant class of natural products found in various microorganisms and plants, have attracted considerable interest due to their broad spectrum of biological activities and unique molecular architecture. These compounds are characterized by a spiro fusion of oxindole core with a cycloalkyl moiety, creating a quaternary stereocenter at the spiro junction (Fig. 1a). The structural complexity and stereochemical diversity of spirooxindole alkaloids contribute to their wide range of biological properties, including anticancer, antihypertensive, anti-inflammatory, and anti-neurodegenerative properties^1,2,3,4,5^. Inspired by their therapeutic potential and structure complexity, synthetic chemists have developed various strategies for the stereospecific synthesis of individual spirooxindole scaffolds (Fig. 1b)^6,7,8^. The inefficiency and resource-intensive nature of these single-target approaches have motivated the development of collective synthesis strategies targeting common intermediates that can access various natural product families^9,10^. This concept of common intermediates draws inspiration from nature’s biosynthetic logic, and implementing a collective biosynthetic approach – where multiple related scaffolds are generated through a unified set of enzymes – represents a potentially more sustainable alternative to chemical synthesis. However, unlike the versatile chemical reagents employed in collective synthesis, many biosynthetic pathways remain incompletely elucidated, and biosynthetic enzymes often exhibit narrow substrate scope, limiting their application in collective biosynthesis of natural products.

**Fig. 1:**
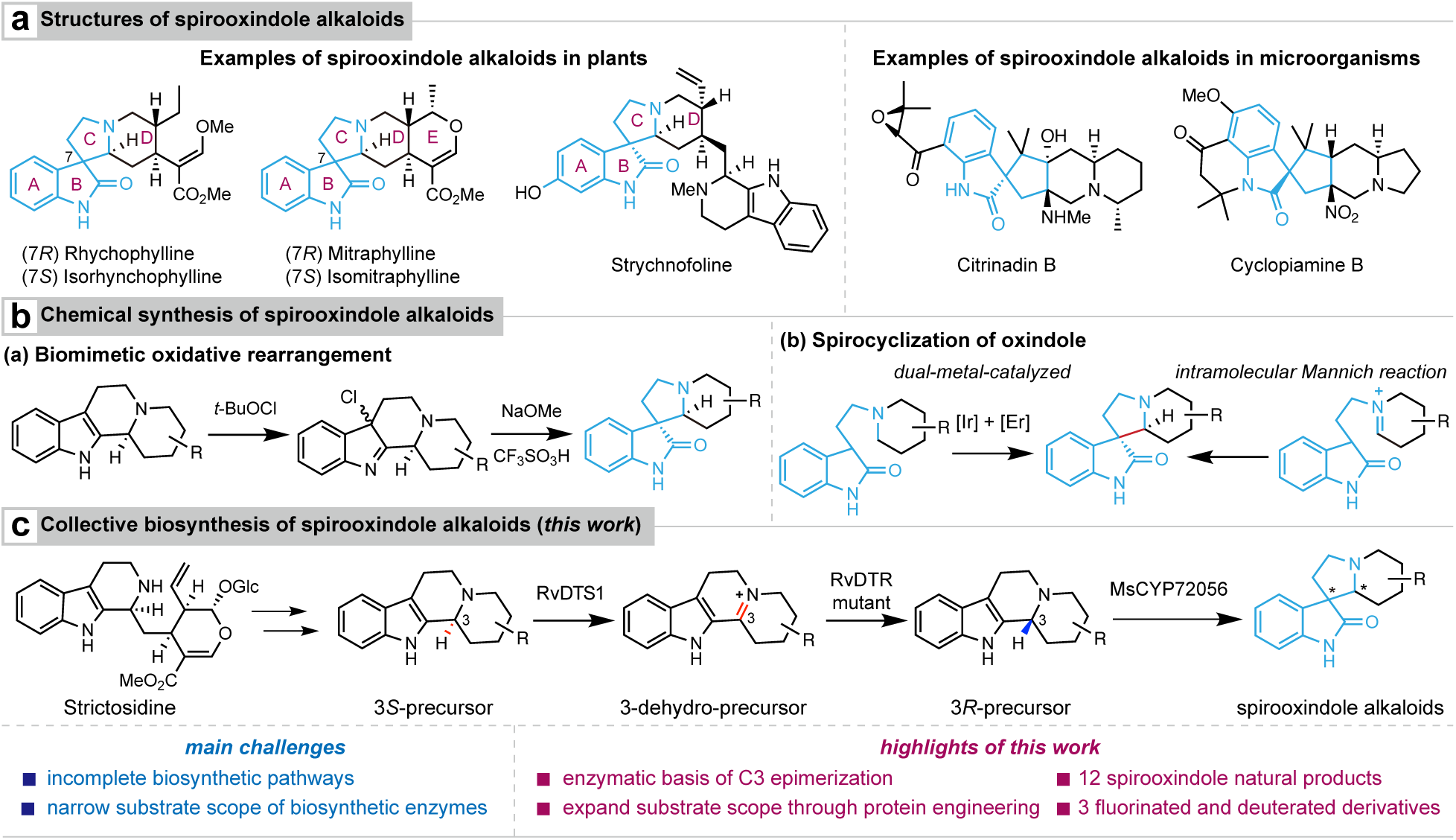
Overview of spirooxindole alkaloids and their synthetic approaches. **a**, Representative structures of spirooxindole alkaloids found in plants and microorganisms. **b**, Chemical approaches for spirooxindole alkaloid synthesis. **c**, Collective biosynthetic route developed in this work, showing the conversion of strictosidine to spirooxindole alkaloids through a sequence of enzymatic transformations involving RvDTS1, engineered RvDTR mutant, and MsCYP72056.

Plant-derived spirooxindole alkaloids, predominantly isolated from the *Apocynaceae*, *Gelsemiaceae*, and *Rubiaceae* families, are a subclass of monoterpene indole alkaloids (MIAs) ^11,12,13,14^. The well-characterized MIA biosynthetic pathway begins with strictosidine synthase (STR) catalyzing the Pictet-Spengler condensation of tryptamine and secologanin to form 3*S*- strictosidine, a common intermediate in MIA biosynthesis^15^. Following deglycosylation by strictosidine glucosidase (SGD), the resulting reactive strictosidine aglycone undergoes NADPH-dependent reduction by various medium-chain dehydrogenases/reductases (MDRs) ^16,17,18,19,20,21,22,23^. This reduction yields tetracyclic corynanthe type alkaloids and pentacyclic heteroyohimbine type alkaloids, maintaining the 3*S* configuration. The prevalence of 3*S* configuration in plant spirooxindole alkaloids led to the hypothesis that their skeleton derives from oxidative rearrangement of 3*S* precursors’ indole rings^24,25^, a mechanism supported by flavin-dependent monooxygenases in microbial paraherquamide and citrinadin biosynthesis^26,27^. However, the recent discovery of cytochrome P450 enzyme MsCYP72056 from *Mitragyna speciosa* has challenged this hypothesis^28,29^. This enzyme oxidizes tetracyclic 3*R*-hirsuteine to form two tetracyclic spirooxindole alkaloids, yet notably cannot recognize C3 epimerized hirsuteine. While the discovery has advanced our understanding of spirooxindole biosynthesis, questions remain about its applicability to pentacyclic precursors and the broader biosynthesis of 3*R* monoterpene indole alkaloids in plants.

While natural biosynthetic pathways offer promising opportunities for accessing large collections of structurally related scaffolds, the narrow substrate scope of biosynthetic enzymes often limits their broader application in collective biosynthesis. Recent advances in protein engineering have begun to address these limitations through various strategies, including active site saturated mutagenesis, rational protein design, and directed evolution. These approaches have successfully expanded the reaction scope, enhanced stereoselectivity, and enabled new-to-nature chemistry ^30,31^. Notable achievements include Lei and coworkers’ engineering of a FAD-dependent enzyme for high-yield late-stage enzymatic oxidation in alchivemycin A synthesis^32^. Similarly, O’Connor and coworkers achieved inverted stereoselectivity through targeted mutation of dihydrocorynantheine synthase’s binding pocket^21^. Despite these advances, the application of such engineering strategies to collective biosynthesis remains largely unexplored.

Here, we present a combined enzyme discovery and protein engineering approach for collective biosynthesis of spirooxindole alkaloids (Fig. 1c). Through multiomics analysis, we identified a flavin-dependent enzyme RvDTS1 and a medium-chain dehydrogenase/reductase (MDR) RvDTR that epimerizes 3*S*-tetrahydroalstonine (**5**) to 3*R*-akuammigine (**6**). Subsequent cytochrome P450- mediated oxidative rearrangement yields uncarines C, D, E, and F (**20-23**). Protein engineering of RvDTR expanded the substrate scope, enabling collective biosynthesis of 12 tetracyclic and pentacyclic spirooxindole natural products. Finally, we employed a precursor-directed biosynthesis approach to create 3 new-to-nature halogenated and deuterated spirooxindole alkaloids in planta. This work demonstrates the potential of combining enzyme discovery and protein engineering to realize collective biosynthesis of diverse collections of structurally related natural products and their derivatives, offering a sustainable alternative to traditional synthetic approaches.

## Results & Discussion

### Biosynthetic proposals of plant spirooxindole alkaloids

Successful implementation of collective biosynthesis relies on enzymes with broad substrate scope. While the biosynthetic pathways of spirooxindole alkaloids remain largely unexplored, recent work by Dang and coworkers showed that two tetracyclic spirooxindole alkaloids are biosynthesized from 3*R*-hirsuteine (**12**) through oxidative rearrangement and subsequent C3 isomerization, catalyzed by a *M. speciosa* cytochrome P450 enzyme MsCYP72056^28^. To investigate whether this biotransformation could be extended to pentacyclic spirooxindole alkaloids and evaluate the enzyme’s potential utility in collective biosynthesis, we transiently expressed *MsCYP72056* in *Nicotiana benthamiana* via agrobacterium-mediated transformation. The transformed plants were then infiltrated with various commercially available pentacyclic alkaloids: 3*S*-tetrahydroalstonine (**5**), 3*S*-ajmalicine (**7**), 3*S*-mayumbine (**9**), and 3*R*-akuammigine (**6**) (Supplementary Fig. 1). LC-MS analysis of the methanol extracts of tobacco leaves revealed that uncarine C (**20**), D (**21**), E (**22**), and F (**23**) were produced exclusively from akuammigine (**6**). This strict stereochemical selectivity aligns with the previously established 3*R*-precursor specificity of MsCYP72056 (Supplementary Fig. 1), suggesting a conserved oxidative pathway for both tetracyclic and pentacyclic spirooxindole alkaloid biosynthesis from 3*R* precursors.

The biosynthetic origin of 3*R* tetracyclic and pentacyclic indole alkaloids in plants, however, remains unclear. The co-occurrence of 3*S*-tetrahydroalstonine (**5**) and 3*R*-akuammigine (**6**), as well as 3*S*-corynoxeine (**11**) and 3*R*-hirsuteine (**12**), within the same plants suggested that 3*R* alkaloids might be derived from their 3*S* counterparts through epimerization. Epimerization could be catalyzed by epimerases, enzymes known to be critical in both primary metabolism and secondary metabolite biosynthesis^33,34^. A precedent for enzymatic epimerization exists in morphine biosynthesis, where reticuline epimerase (REPI) catalyzes *S*-reticuline to *R*-reticuline conversion through sequential oxidation and reduction by its two domains (Supplementary Fig. 2)^35^. The enzyme’s N-terminal cytochrome P450 domain oxidizes *S*-reticuline to an iminium intermediate, which is then reduced stereospecifically to *R*-reticuline by the C-terminal aldo-keto reductase (AKR) domain. Based on these observations, we propose that plant spirooxindole alkaloid biosynthesis follows a similar pathway, where 3*S* precursors undergo enzymatic epimerization to 3*R* precursors through sequential oxidation and reduction, followed by cytochrome P450 mediated oxidation to form spirooxindole alkaloids (Fig. 1c).

### Identification and characterization of 3-dehydroalstonine synthase (RvDTS1)

To identify genes involved in tetracyclic and pentacyclic spirooxindole formation, we focused on two Chinese medicinal plants from the *Apocynaceae* and *Rubiaceae* families, *Rauvolfia verticillata* and *Uncaria rhynchophylla*. Previous phytochemical analysis reported pentacyclic spirooxindole alkaloids from *Rauvolfia* species and tetracyclic spirooxindole alkaloids as major bioactive components in *U. rhynchophylla*^11,13^. Both plants also produce various 3*R* indole alkaloids, making them suitable for studying 3*S* epimerization. We thus collected leaf, stem, and root tissues in triplicate from each species for transcriptome sequencing.

Our biosynthetic hypothesis for spirooxindole alkaloid formation begins with the oxidation of 3*S* precursors to an iminium intermediate, which has been isolated in plants^36^. In *Catharanthus roseus*, a flavin-dependent enzyme, precondylocarpine acetate synthase (PAS), oxidizes the tertiary amine in stemmadenine acetate to an iminium intermediate during vinblastine biosynthesis^37^. We hypothesized that a PAS homolog might similarly oxidize tetrahydroalstonine (**5**). We identified four *CrPAS* homolog genes in the *R. verticillata* transcriptome and tested their oxidative ability toward **5** in *N. benthamiana* (Fig. 2a and Supplementary Fig. 3). LC-MS analysis of leaf extracts revealed that one candidate gene catalyzed the formation of a product with the exact mass ([M+H]^+^ = 351.1709) corresponding to an oxidized **5** derivative (Fig. 2b). We confirmed the structure of this product by comparing it to a chemically synthesized standard of 3-dehydro- tetrahydroalstonine (**15**) (Supplementary Fig. 3), which showed identical exact mass, retention time, and fragmentation patterns. Based on the product of this oxidase, we named this enzyme as 3-dehydro-tetrahydroalstonine synthase (RvDTS1).

**Fig. 2:**
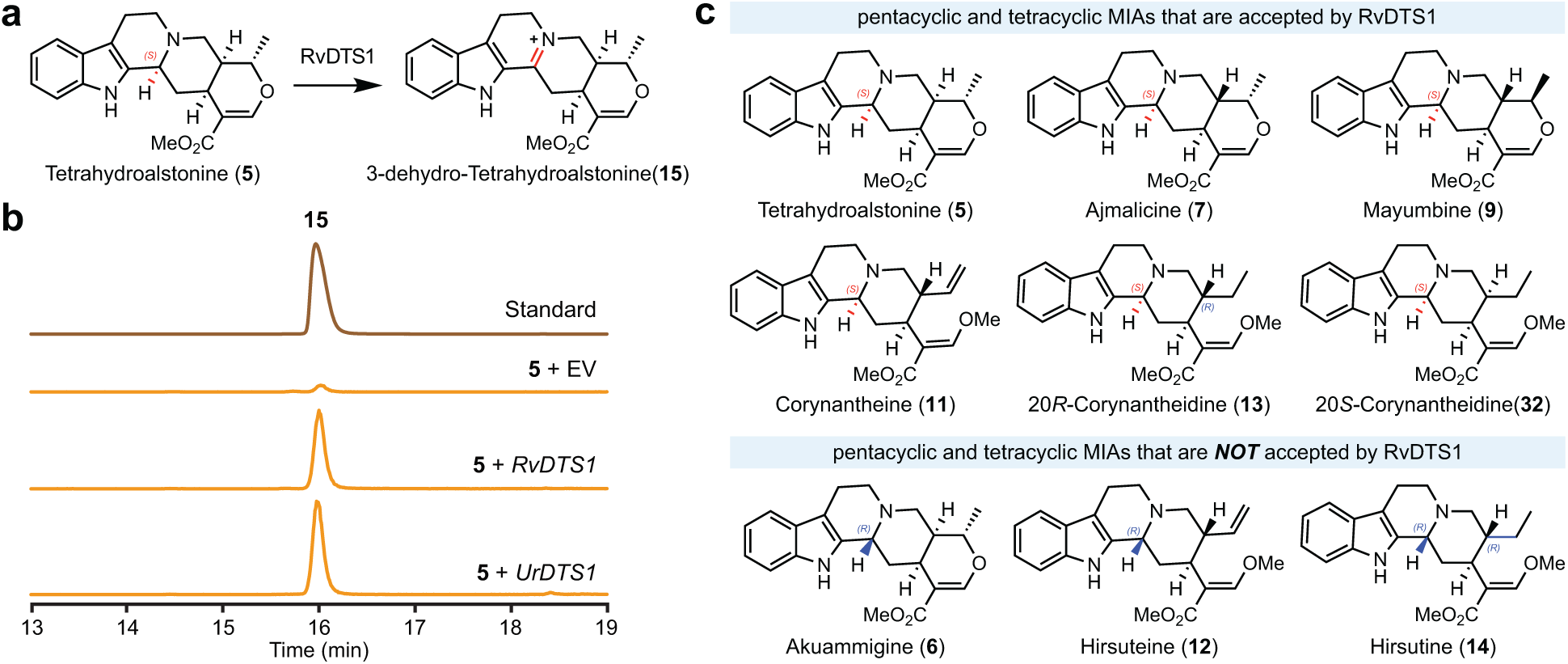
Discovery and characterization of RvDTS1 **a**. Biosynthetic pathway showing the conversion of tetrahydroalstonine (**5**) to 3-dehydro- tetrahydroalstonine (**15**) catalyzed by RvDTS1. **b**, LCMS analysis of the formation of **15**. Traces include synthetic standard, empty vector control (EV), *RvDTS1*, and *UrDTS1* reactions. **c**, Substrate scope of RvDTS1, highlighting accepted substrates (top panel) and non-accepted 3*R*- configured substrates (bottom panel).

To explore the conservation of this FAD-dependent oxidation mechanism across the *Rubiaceae* family, we searched for *RvDTS1* homologs in the *U. rhynchophylla* transcriptome. We identified *UrDTS1*, sharing 59% sequence identity with *RvDTS1*, and functionally characterized through transient expression in *N. benthamiana* leaves infiltrated with tetrahydroalstonine (**5**). The successful formation of 3-dehydro-tetrahydroalstonine (**15**) upon *UrDTS1* expression suggests that this oxidative mechanism is conserved within the *Rubiaceae* family (Fig. 2b).

To further evaluate RvDTS1’s substrate scope, we tested various putative oxindole biosynthetic precursors in *RvDTS1*-overexpressing *N. benthamiana* leaves. LC-MS analysis showed that RvDTS1 oxidized precursors with 3*S* configuration, including pentacyclic ajmalicine (**7**) and mayumbine (**9**), and tetracyclic corynantheine (**11**), 20*R*-corynantheidine (**13**), or 20*S*- corynantheidine (**32**) (Fig. 2c and Supplementary Fig. 4 - 8). Precursors with 3*R* configuration, such as akuammigine (**6**), hirsuteine (**12**), and hirsutine (**14**) remain unoxidized, indicating RvDTS1’s stereospecificity for 3*S* alkaloids. These results also suggest that the 3*S* to 3*R* stereochemical inversion is irreversible through enzymatic mechanism, as the resulting 3*R* products resist RvDTS1 oxidation.

### Identification and characterization of 3-dehydrotetrahydroalstonine reductase (RvDTR)

We next sought to identify the reductase responsible for stereospecific reduction of the iminium intermediate from the convex face to generate 3*R* precursors (Fig. 3a). In MIA biosynthesis, iminium reduction is typically catalyzed by medium-chain dehydrogenase/reductase (MDR) enzymes, as exemplified by tetrahydroalstonine synthase (THAS) in *Catharanthus roseus* that performs 1,2-reduction of iminium strictosidine aglycone to yield tetrahydroalstonine^16^. To identify potential MDRs capable of reducing the iminium intermediate, we constructed a phylogenetic tree of MDRs in *R. verticillata* and *U. rhynchophylla* with previously characterized MDRs in MIA biosynthetic pathway (Fig. 3b and Supplementary Fig. 9). We screened MDR candidate genes within the clades of *CrGS*, *CrHYS*, and *CrTHAS* through transient overexpression with *RvDTS1* in *N. benthamiana* leaves, using tetrahydroalstonine (**5**) as substrate. LC-MS analysis revealed that two MDR candidates, *RvDTR8472* (*RvDTR*) and *RvDTR13501* (97% sequence identity), produced a new peak matching the exact mass, retention time, and fragmentation pattern of akuammigine (**6**), the 3*R* stereoisomer of **5** previously isolated from *U. rhynchophylla* (Fig. 3c and Supplementary Fig. 10). In planta experiments lacking both *RvDTS1* and *RvDTR* showed no product formation, while those lacking only *RvDTS1* yielded a minor peak, likely resulting from the reduction of an auto-oxidized product of **5**. Given their functional equivalence, we used RvDTR for the following experiments.

**Fig. 3:**
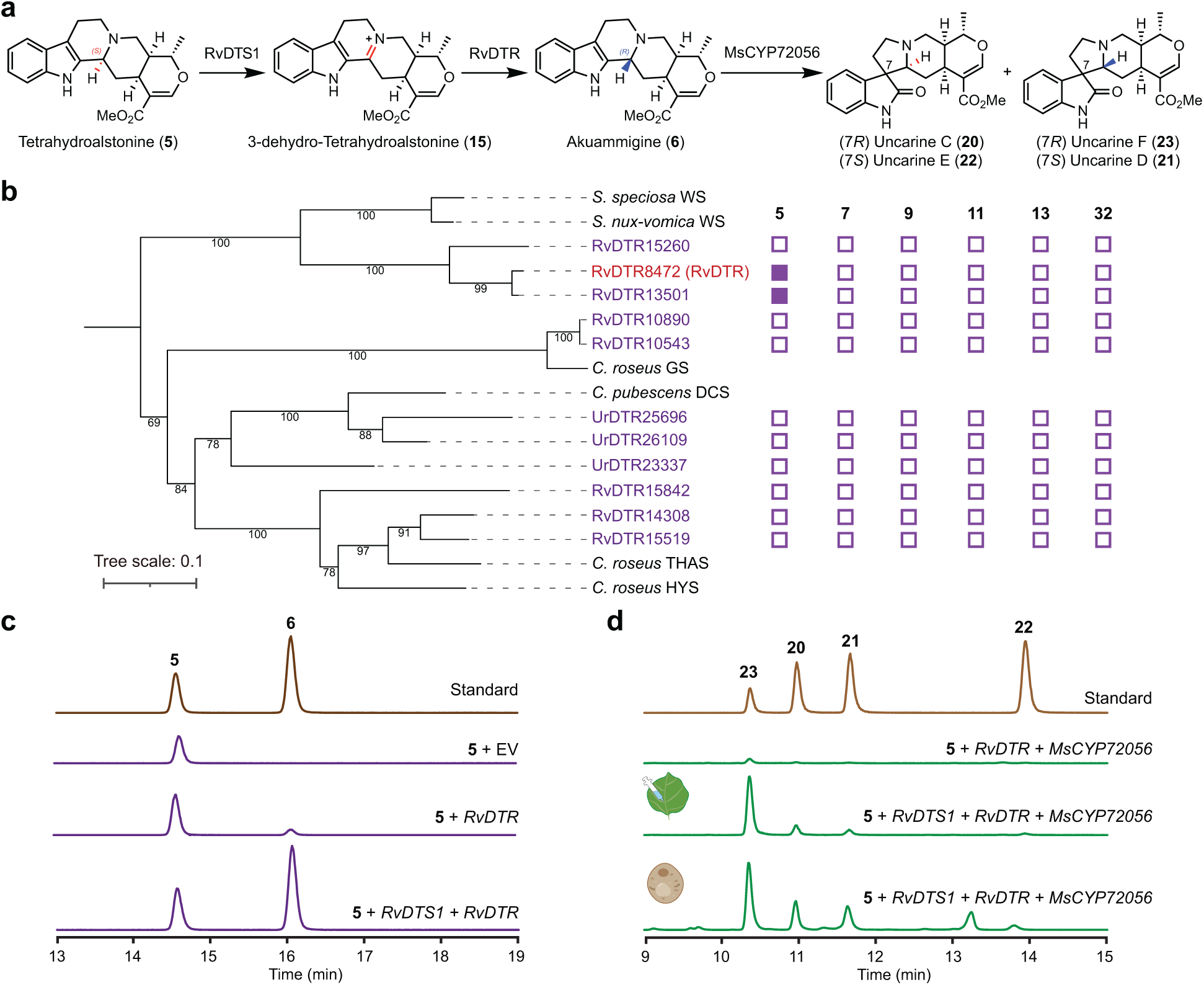
Identification and characterization of RvDTR in pentacyclic spirooxindole alkaloid biosynthesis**. a**, Biosynthetic pathway showing the conversion of tetrahydroalstonine (**5**) to uncarine isomers (**20**-**23**). **b**, Phylogenetic analysis of medium-chain dehydrogenases/reductases (MDRs) from *R. verticillata*, *U. rhynchophylla*, and known MDR enzymes. Substrate screening results are indicated by filled (active) and empty (inactive) purple squares for various compounds (**5**, **7**, **9**, **11**, **13**, and **32**). **c**, LC-MS analysis of the sequential conversion of tetrahydroalstonine (**5**) to akuammigine (**6**) by *RvDTS1* and *RvDTR*. **d**, LC-MS analysis of uncarine isomer (**20**-**23**) production in tobacco leaves (indicated by leaf icon) and in yeast (indicated by yeast cell icon).

To confirm the in planta production of akuammigine (**6**), we co-expressed *MsCYP72056* with *RvDTS1* and *RvDTR* in *N. benthamiana* leaves and infiltrated tetrahydroalstonine (**5**) as substrate. LC-MS analysis revealed four spirooxindole products with exact masses identical to uncarine C (**20**), D (**21**), and E (**22**), and F (**23**), with uncarine F (**23**) predominating (Fig. 3d). This confirmed the successful enzymatic epimerization of **5** to 3*R*-akuammigine (**6**) in planta. Having established the pathway in planta, we next sought to demonstrate its functionality in a microbial host. A *Saccharomyces cerevisiae* (baker’s yeast) strain expressing *RvDTS1*, *RvDTR*, and *MsCYP72056* was cultured in synthetic complete medium supplemented with **5**. After five days of fermentation, **20**-**23** were detected in the culture (Fig. 3d). This heterologous reconstitution in both tobacco and yeast demonstrated successful elucidation of the pentacyclic spirooxindole alkaloid biosynthetic pathway.

We then investigated whether other 3*R* precursors could be formed in a similar mechanism as akuammigine. We tested pentacyclic ajmalicine (**7**) and mayumbine (**9**), and tetracyclic corynantheine (**11**), 20*R*-corynantheidine (**13**), and 20*S*-corynantheidine (**32**) by infiltration into tobacco leaves expressing *RvDTS1* and MDR candidate genes. LC-MS analysis showed no major peaks with masses corresponding to potential products at retention times distinct from the substrates (Fig. 3b). To account for potential overlapping retention times between substrates and products, we included cytochrome P450 *MsCYP72056* in the assays. However, no spirooxindole products were detected. These results suggest that while tetrahydroalstonine (**5**) epimerization occurs through consecutive oxidation and reduction, the enzymatic basis for epimerization of other precursors remained unclear. The epimerization of non-tetrahydroalstonine precursors likely involves non-DTS type reductases, though further investigation is needed to fully elucidate these biosynthetic pathways. While we were preparing this manuscript, two independent studies reported enzymatic mechanisms for C3 epimerization in MIA biosynthesis^38,39^. Interestingly, O’Connor and coworkers reported the enzymatic epimerization in *M. speciosa*, where an isoflavone reductase homolog can reduce 3-dehydrotetrahydroalstonine (**15**) but not 3-dehydroajmalicine (**16**), further highlighting the challenges in identifying reductases for 3-dehydro MIAs from nature.

### Protein engineering of RvDTR for expanded substrate scope

While the successful reconstitution of uncarine C (**20**), D (**21**), E (**22**), and F (**23**) in tobacco and yeast demonstrated the utility of RvDTR, its high specificity towards 3-dehydro- tetrahydroalstonine (**15**) prevented its use as a biocatalytic tool for the collective biosynthesis of spirooxindole alkaloids. Recent advances in protein engineering have significantly expanded substrate scope and reaction diversity of numerous biosynthetic enzymes^30^. Inspired by O’Connor and coworker’s recent success in engineering dihydrocorynantheine synthase (DCS) to invert the ratio between two stereoisomer products^21^, we decided to expand the catalytic scope of RvDTR through protein engineering.

Given that 3-dehydro-tetrahydroalstonine (**15**) and 3-dehydro-ajmalicine (**16**) differ only in their C20 stereochemistry, we hypothesized that subtle modifications to RvDTR’s substrate binding pocket might enable recognition of **16**. To guide our engineering efforts, we constructed a structural model of RvDTR complexed with **15** and NADPH using AlphaFold 3 and Autodock Vina (Fig. 4a)^40,41^. Based on this model, we selected eight residues proximal to the binding pocket for site-directed mutagenesis (Fig. 4a and Supplementary Fig. 11). The resulting mutants were coexpressed with *RvDTS1* in *N. benthamiana* and infiltrated with tetrahydroalstonine (**5**) and ajmalicine (**7**). While these mutations did not affect akuammigine (**6**) formation (Supplementary Fig. 12-14), four mutants (H99A, H99S, T123A, and T123G) generated a novel product with identical mass and fragmentation pattern as ajmalicine (**7**), with T123G showing the highest conversion efficiency (Fig. 4b and Supplementary Fig. 15). This product, barely detected in wild-type (WT) RvDTR, was predicted to be 3-epi-ajmalicine (**8**). To confirm this structure, we incorporated *MsCYP72056* into the assay, and LC-MS analysis revealed co-elution of mitraphylline (**24**) and isomitraphylline (**25**) standards with products from the mutant-expressing tobacco leaves (Supplementary Fig. 16). These findings demonstrate that a single amino acid substitution in RvDTR enables the enzyme to recognize 3-dehydro-ajmalicine (**16**).

**Fig. 4:**
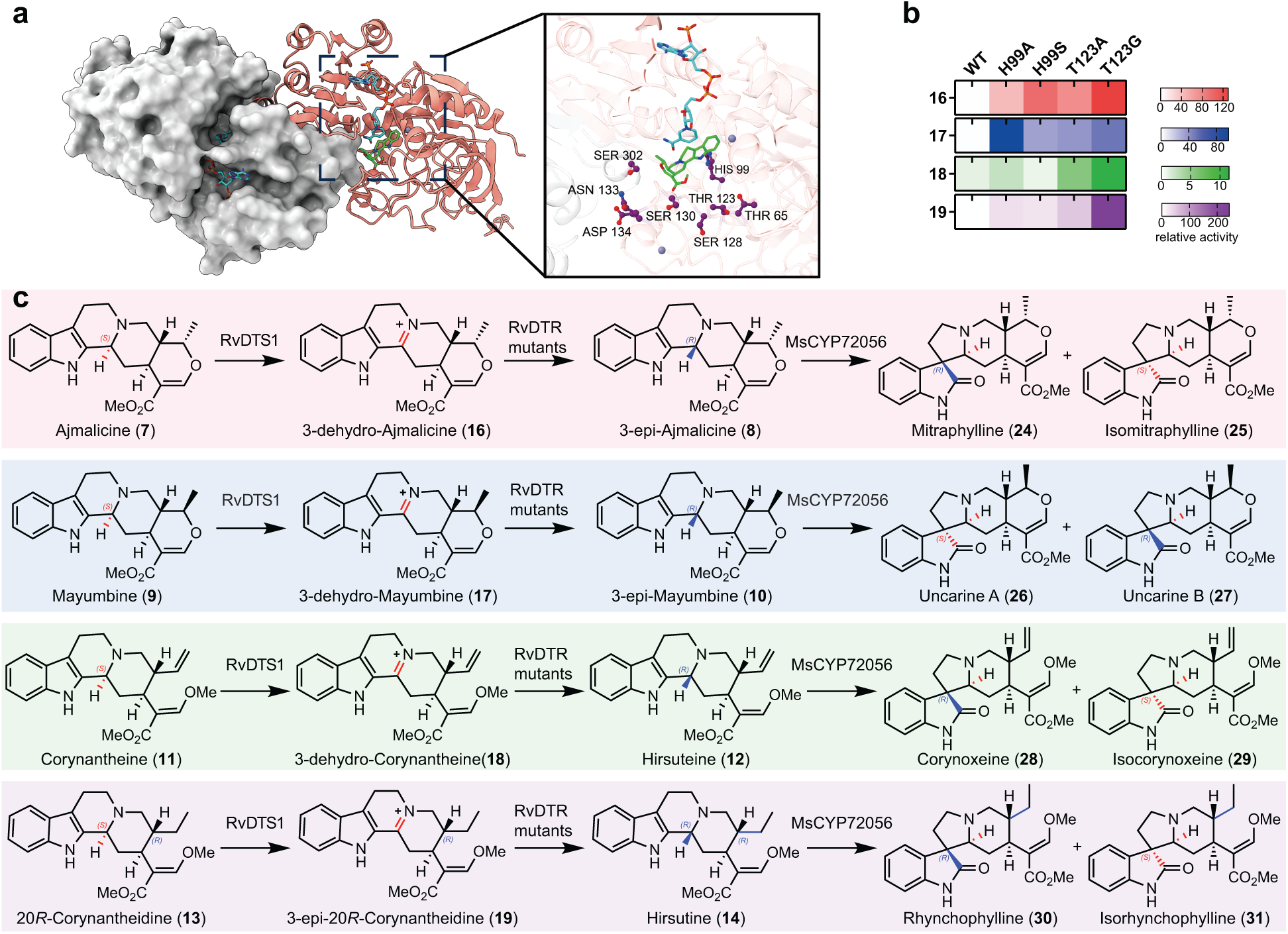
Protein engineering and substrate scope expansion of RvDTR **a**, Overview of the three-dimensional model of RvDTR docked with 3-dehydro- tetrahydroalstonine (**15**). The dimer structure is shown as a surface model in gray and a cartoon model in hot pink. The zoom area is the close-up view of the active pocket of RvDTR. The substrate-binding pocket is shown as a cartoon model, with NADPH, substrate, activity- maintained mutation sites represented as a ball stick model in cyan, lime, and purple, respectively. **b**, Heatmap showing the relative activities of wild-type RvDTR and its mutants toward different 3-dehydro substrates (**16**-**19**). Color intensity indicates product formation normalized to wild-type activity. **c**, Biosynthetic pathways illustrating the production of diverse spirooxindole alkaloids using engineered RvDTR mutants.

Encouraged by these results, we broadened our study to examine additional 3*R* precursors, including pentacyclic and tetracyclic substrates. In *N. benthamiana* leaves expressing *RvDTS1* and *RvDTR* mutants, infiltration with mayumbine (**9**), corynantheine (**11**), or 20*R*-corynantheidine (**13**) yielded their respective 3-epi products (Fig. 4b and Supplementary Fig. 17, 19, 21). Further incorporation of *MsCYP72056* led to the production of uncarine A (**26**) and B (**27**) from mayumbine (**9**), corynoxeine (**28**) and isocorynoxeine (**29**) from corynantheine (**11**), and rhynchophylline (**30**) and isorhynchophylline (**31**) from 20*R*-corynantheidine (**13**) (Supplementary Fig. 18, 20, 22). The engineered enzymes exhibited broad substrate scope, demonstrating the versatility of RvDTR mutants in producing diverse spirooxindole alkaloids (Fig. 4c).

To understand the molecular basis underlying the enhanced reactivity of RvDTR mutants, we performed molecular dynamics (MD) simulations to the best-performing T123G variant using the Amber program package. The simulations revealed that in the T123G mutant, the distance between the carbon atom of iminium in 3-dehydroajmalicine (**16**) and the C4N atom of NADPH was shorter compared to the WT RvDTR (3.9 Å versus 4.1 Å, respectively; Supplementary Fig. 23). Our computational model further identified a hydrogen bond network connecting T123 with D125, W53, and T127, which confers rigidity to the gate loop in the WT enzyme. We propose that the T123G mutation disrupts the hydrogen bond network, increasing the flexibility of the gate loop and transforming RvDTR from a substrate-specific enzyme into one with broader substrate scope (Supplementary Fig. 24).

### Generation of new-to-nature spirooxindole derivatives through precursor-directed biosynthesis

Having established the reconstitution of spirooxindole natural products and expanded the substrate scope of RvDTR, we next explored the generation of new-to-nature spirooxindole analogues through precursor-directed biosynthesis (Fig. 5a, 5b). Given that strictosidine synthase exhibits promiscuous activity towards tryptamine derivatives, previous studies have shown that feeding 4-, 5-, 6-, or 7-substrituted tryptamines can yield diverse MIA derivatives in native plants and in engineered yeast strains^24,42,43^.

**Fig. 5:**
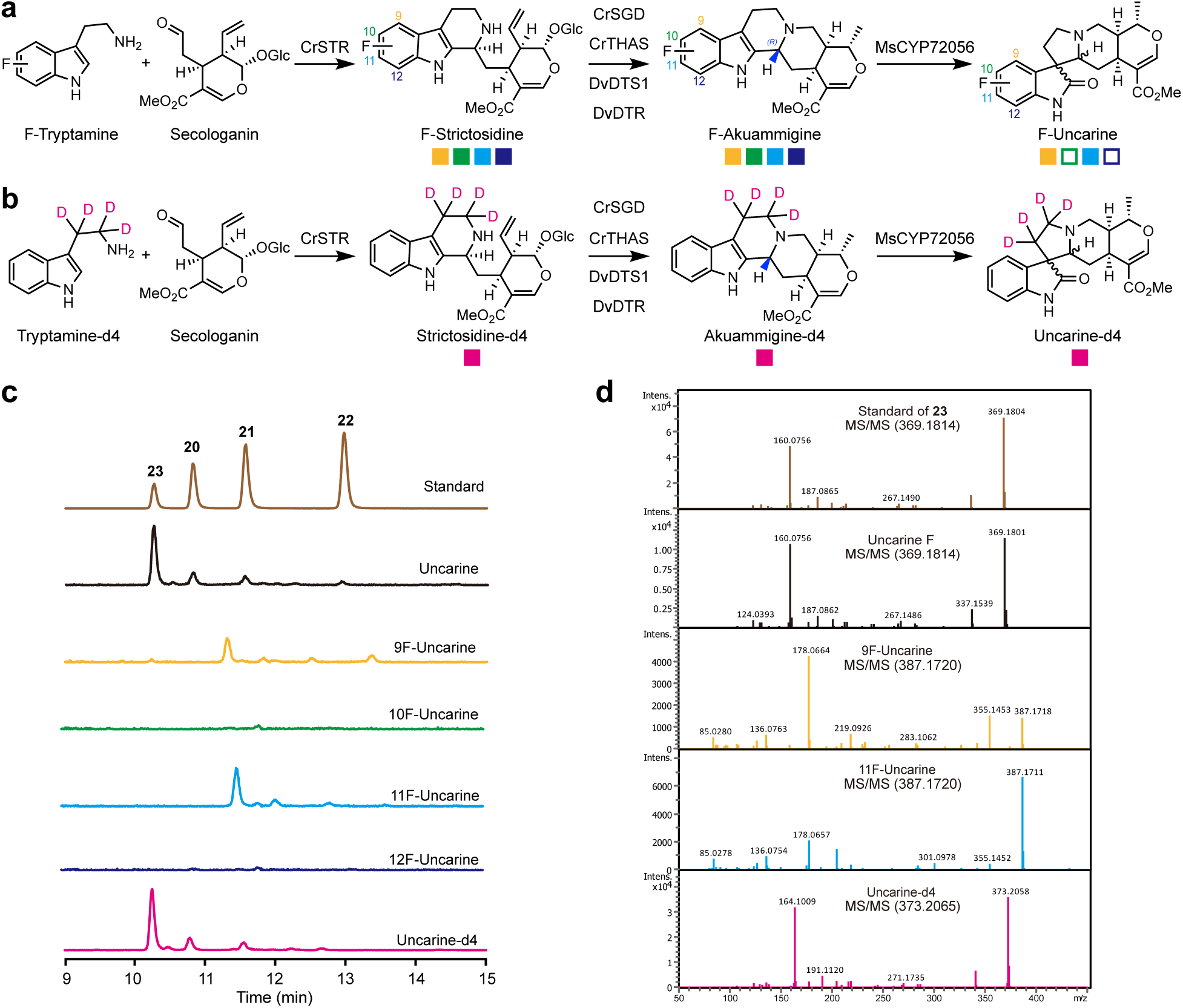
Precursor-directed biosynthesis of novel spirooxindole analogues. **a**, Biosynthetic scheme showing the conversion of fluorinated tryptamine and secologanin to fluorinated uncarine derivatives. The colored squares correspond to different fluorine substitution positions (yellow: 9F-, green: 10F-, light blue: 11F-, dark blue: 12F-) on the indole ring, with filled squares indicating detected compounds and empty squares showed undetected derivatives. **b**, Biosynthetic scheme showing the conversion of deuterated tryptamine and secologanin to deuterated uncarine derivatives. Filled magenta squares indicate detected compounds. **c**, Extracted ion chromatograms showing the production of native uncarines (**20**-**23**) and modified derivatives (9F-, 10F-, 11F-, 12F-, and D4-uncarine) in *N. benthamiana*. **d**, MS/MS analysis comparing uncarine F (**23**) standard with fluorinated and deuterated uncarine derivatives.

To develop a platform for producing spirooxindole analogues from substituted tryptamine, we first validated the production of uncarine C (**20**), D (**21**), E (**22**), and F (**23**) through feeding of tryptamine (**1**) in *N. benthamiana*. We transiently overexpressed *C. roseus* strictosidine synthase (*CrSTR*), *C. roseus* strictosidine-β-D-glucosidase (*CrSGD*), *CrTHAS*, *RvDTS1*, *RvDTR*, and *MsCYP72056* in tobacco leaves and infiltrated with **1** and secologanin (**2**). LC-MS analysis revealed the production of **20**-**23** (Fig. 5c), suggesting that tobacco is a suitable platform for the production of halogenated spirooxindole.

The introduction of halogens to natural products can enhance their biological activities and pharmacological properties. To investigate the potential of RvDTR in producing halogenated spirooxindoles, we transiently expressed the biosynthetic genes in *N. benthamiana* and infiltrated with **2** and various fluorinated tryptamines. New peaks corresponding to fluorinated uncarines were detected upon supplementation with 4- and 6- fluorinated tryptamine (Fig. 5c, 5d, Supplementary Fig. 25-28). MS/MS analysis revealed a mass shift of 18 Da, confirming fluorine incorporation into uncarines. While 5- and 7-fluorinated tryptamine failed to yield uncarines derivatives, they produced fluorinated akuammigine, likely due to insufficient substrate recognition by MsCYP72056. Engineering of this P450 enzyme could potentially enable the production of spirooxindole incorporating 5- and 7-position substitutions.

Beyond halogenation, deuterium incorporation into complex natural products has been shown to enhance metabolic stability, potentially improving therapeutic efficacy and reducing dosing frequency^44^. Deuterated natural products could also serve as research tools for precise tracking in complex biological systems. Substituting tryptamine (**1**) with D_4_-tryptamine in our uncarine reconstitution system yielded deuterated uncarines, characterized by a distinctive mass shift of +4 Da (Fig. 5c, 5d). These results demonstrate the versatility of our platform for generating novel spirooxindole derivatives with enhanced properties and potential therapeutic applications.

## Conclusion

To achieve collective biosynthesis of spirooxindole alkaloids, we first elucidated the biosynthetic pathway for pentacyclic spirooxindole alkaloids in plants, revealing the sequential actions of RvDTS1 and RvDTR in the stereospecific epimerization of 3*S*-tetrahydroalstonine (**5**) to 3*R*- akuamigine (**6**). The discovery of these enzymes, alongside the cytochrome P450 enzyme identified in *M. speciosa*, establishes the enzymatic framework for the bioproduction of spirooxindole alkaloids in both *N. benthamiana* and *S. cerevisiae*. Through protein engineering of RvDTR, we significantly expanded its substrate scope, achieving the collective biosynthesis of 12 structurally diverse tetracyclic and pentacyclic spirooxindole natural products from a single biosynthetic platform. This collective biosynthetic approach represents a significant advancement over traditional approaches that require separate elucidation and optimization of individual pathways. The platform’s versatility was further demonstrated through successful incorporation of halogenated and deuterated precursors, enabling rapid access to novel derivatives. Our strategy of combining enzyme discovery and engineering to achieve collective biosynthesis could be broadly applicable to other families of complex natural products, offering a powerful approach for sustainable production of diverse molecular collections with therapeutic potential.

## Methods

### RNA-sequencing

The tissues of *Rauvolfia verticillata* (roots, stem, leaves) and *Uncaria rhynchophylla* (roots, stem, leaves, hook) were collected and immediately flash-frozen in liquid nitrogen. Approximately one gram of frozen tissue samples were submitted to Novogene Corporation (Beijing, China) for total RNA sequencing. RNA-seq libraries were prepared following the companýs standard protocols. Sequencing was performed on an Illumina NovaSeq 6000 platform, generating > 30 million paired-end raw sequencing reads per sample.

### Cloning of gene candidates and mutagenesis

Candidate genes for *RvDTS* were identified through BLAST analysis using CrPAS (GenBank accession number AWJ76616.1) as the query sequence. *RvDTR* candidates were similarly identified using previously characterized medium-chain dehydrogenase/reductases (MDRs) as query sequences (Supplementary Table 4). All candidate genes were commercially synthesized and cloned into 3Ω1 plasmids^45^ by GenScript Biotech Corporation (Nanjing, China).

#### Construction of bacterial expression vectors

Genes of interest were cloned into either pOPINF vectors (containing an N-terminal His6-tag) or pOPINM vectors (containing an N-terminal MBP-tag). The vectors were linearized using HindIII and KpnI restriction enzymes. In-Fusion assembly products were transformed into chemically competent *E. coli* TOP10 cells, and transformants were selected on LB agar plates containing 100 μg/mL ampicillin. Positive clones were identified by colony PCR and verified by Sanger sequencing.

#### Construction of yeast expression vectors

*RvDTS1* and codon-optimized *RvDTR*, *MsCYP72056*, and *Arabidopsis thaliana* cytochrome P450 reductase *AtATR2* were cloned into pESC yeast epitope tagging vectors (pESC-URA and pESC-LEU) using the Hieff Clone Universal II One Step Cloning Kit. Genes were inserted into multiple cloning sites (MCS1: BamHI/HindIII; MCS2: SpeI) according to the manufacturer’s protocol. Primer sequences are provided in Supplementary Table 3.

#### Site-directed mutagenesis

Site-directed point mutations were introduced into the *RvDTR* gene by PCR-based mutagenesis. The mutagenesis strategy involved amplifying the gene in two fragments, each containing complementary overlapping sequences with the desired mutations. These fragments were then ligated into linearized 3Ω1 vector using In-Fusion cloning. The 3Ω1 empty vector was linearized with the restriction enzyme BsaI. The specific mutations and primer sequences are listed in Supplementary Table 3. All constructs were verified by Sanger sequencing before using for downstream applications.

### Transformation of Agrobacterium tumefaciens

The 3Ω1 plasmids containing the full-length gene were transformed into *A. tumefaciens* (strain GV3101) cells following the manufacturer’s protocol. The transformed cells were plated onto LB agar plates containing 25 µg/mL rifampicin, 25 µg/mL gentamycin, and 50 µg/mL spectinomycin, and incubated at 28 °C for 2-3 days. Single colonies were picked and grown overnight at 28 °C in 2 mL of LB medium supplemented with 25 µg/mL rifampicin, 25 µg/mL gentamycin, and 50 µg/mL spectinomycin with a shaking of 220 rpm. The overnight cultures were mixed with an equal volume of 50% glycerol and stored at -80 °C as frozen stocks.

### Transient expression of candidate genes in *Nicotiana benthamiana*

Transient expression of candidate genes in *N. benthamiana* was performed following the Hawes *et al.* protocol with modifications^46^. In brief, *Agrobacterium* GV3101 strains harboring the constructs of interest were cultivated in 10 mL liquid LB medium supplemented with 25 µg/mL rifampicin, 25 µg/ml gentamycin, and 50 µg/ml spectinomycin overnight at 28 °C with a shaking of 220 rpm. The cells were then harvested by centrifugation at 4,000 rpm for 20 minutes and washed twice with 5 mL of infiltration buffer (50 mM MES pH 5.5, 27.8 mM glucose, 100 µM acetosyringone, 10 mM MgCl_2_). The optical density at 600 nm (OD_600_) was measured for each culture, and bacterial suspensions were mixed as needed to make a 10 mL culture with a final OD of 0.2. The mixtures were then incubated at room temperature for 2 hours in the dark with gentle shaking before being infiltrated into the abaxial (bottom) side of 4-week-old *N. benthamiana* leaves using a needleless 1 mL syringe. After 4 days, 100 μM of substrates were infiltrated into the same leaf area. Leaf tissue was harvested 24 hours post substrate infiltration by collecting six 12-mm diameter leaf disks. Each experiment included at least two biological replicates, with samples taken from two different leaves on separate plants.

### Extraction of metabolites from harvested *N. benthamiana* leaf material

Harvested leaves were immediately placed inside a 2 mL safe-lock tube (Axygen) and ground to a fine powder with liquid nitrogen by pestle. Metabolites were extracted by adding 1mL of HPLC-grade methanol to each sample, followed by sonication at room temperature for 20 minutes. The extracts were centrifuged at 13,000 *g* for 10 minutes at room temperature to remove cell debris. The supernatant was filtered through 0.22 μm nylon syringe filters and stored at -20 °C until analysis.

### Phylogenetic analysis

Protein sequences were aligned using MEGA software (version 7.0.14)^47^ with the MUSCLE algorithm using default parameters. The aligned sequences were manually inspected and trimmed to remove poorly aligned regions. Phylogenetic analysis was performed using IQ-TREE (version 2)^48^ using the Maximum Likelihood method in a bootstrap test of 1,000 replicates. The resulting tree was visualized in iToL v7.

### Yeast strain construction

*S. cerevisiae* strain BY4741 was used as the host strain for transformation. Chemically competent cells were prepared and transformed using the lithium acetate/single-stranded carrier DNA/polyethylene glycol (LiAc/SS carrier DNA/PEG) method as described previously^49^. Briefly, 1 µg of each recombinant plasmid was introduced into competent cells, and transformants were selected on synthetic defined (SD) medium lacking uracil and histidine (SD-URA/LEU). Plates were incubated at 30 °C for 2-3 days. Positive transformants were verified by colony PCR using the primers listed in Supplementary Table 3.

Single colonies of recombinant *S. cerevisiae* strains were inoculated into 1 mL of SD-URA/LEU medium containing 2% glucose and grown overnight at 30 °C with a shaking of 220 rpm. The overnight cultures were diluted in 1:100 in 10 mL of fresh SD-URA/LEU medium containing 2% glucose and grown for an additional 24 hours at 30 °C with a shaking of 220 rpm. For induction, cells were harvested by centrifugation at 4,000 rpm for 5 minutes at room temperature and resuspended in SD-URA/LEU medium containing 2% galactose. The cultures were then incubated with 50 µM tetrahydroalstonine (**5**) at 30 °C with a shaking of 220 rpm for 5 days.

### Extraction of metabolites from harvested yeast

The culture broth was harvested by centrifugation at 4,000 rpm for 10 min at 4°C. The supernatant was extracted by an equal volume of ethyl acetate (1:1, v/v) three times. The organic phases were combined and concentrated to dryness using a rotary evaporator under reduced pressure. The dried samples were resuspended in 150 μL of LCMS-grade methanol and filtered through a 0.22 µm nylon membrane prior to LC-MS analysis. Samples were stored at -20 °C if not analyzed immediately.

### LC-MS/MS

All compounds and extracts were analyzed using liquid chromatography-mass spectrometry (LC-MS). The analytical system consisted of a High-Performance Liquid Chromatography system (HPLC; Thermo Fisher) connected to an Impact II UHR-Q-ToF (Ultra-High Resolution Quadrupole-Time-of-Flight) mass spectrometer (Bruker).

Compound separation was achieved using reverse-phase liquid chromatography on a Waters XBridge BEH C18 column (100 x 3.0 mm, 2.5 µm; 100 Å) operated at 40 °C. The mobile phases consisted of (A) water with 0.1 % formic acid and (B) acetonitrile. The flow rate was 0.3 mL/min. The injection volume was 2 µL in each run. Authentic standards (1-5 µM) were prepared in methanol. The chromatography condition was as follows: 0-2 min, 5 % B; 2-5 min, 5 % B to 20 % B; 5-15 min, 20 % B to 30 % B; 15-18 min, 30 % B to 95 % B; 18-20 min, 95 % B; 20-22 min, 5 % B.

Mass spectrometry analysis was performed using positive electrospray ionization (ESI+) mode with the following source parameters: capillary voltage, 4500 V; end plate offset, 500 V; nebulizer pressure, 2 bar; drying gas (nitrogen) temperature, 220 °C; drying gas flow rate, 10 L/min. Mass spectrometry data was recorded at 12 Hz over a mass range of m/z 50 to 1300 using data-dependent MS acquisition with an active exclusion window of 0.2 min.

For tandem mass spectrometry (MS/MS), fragmentation was triggered on precursor ions exceeding an absolute threshold of 400 counts, with a total cycle time restriction of 3 s. Collision energy was deployed in a stepping option model (30-45 eV). Mass accuracy was ensured by calibrating the system at the beginning of each analytical run using a sodium formate-isopropanol calibration solution, delivered via direct source infusion using an external syringe pump (flow rate: 0.3 μL/min, 1 mL syringe).

### Molecular docking and MD simulation

Molecular dockings of 3-dehydro-tetrahydroalstonine (**15**) and 3-dehydro-ajmalicine (**16**) into RvDTR variants were performed using Autodock Vina package^41^. Protein structures were predicted with AlphaFold 3^40^, with two NADPH cofactors and four zinc metal ions. The 3D conformations of substrates were fully optimized at the B3LYP/def2-SVP level of theory with GD3BJ dispersion correction using Turbomole^50^ (https://www.turbomole.org/).

All MD simulations were performed with the Amber packages^51^. In details, The Amber ff19SB force field was employed for the protein residues, while the general AMBER force field (GAFF)^52^ was used for substrates and NADPHs with RESP2 partial charges^53^ employing particle mesh Ewald^54^ for long-range electrostatic interactions. In addition, the parameters of four zinc ions and coordinated residues were calculated with MCPB.py program^55^.

#### Chemical synthesis of 3-dehyro-tetrahydroalstonine (15) and 3-dehydro-ajmalicine (16)

Compounds **15** and **16** were synthesized according to a previously reported method^9^. The reactions were conducted under argon atmosphere at -20 ℃ as follows: For **16**: *t*-BuOCl (0.4 mmol, 45 μL) in dry CH_2_Cl_2_ (0.2 mL) was added dropwise into a stirred solution of **7** (0.15 mmol, 52.8 mg) and dry Et_3_N (0.3 mmol, 41 μL) in dry CH_2_Cl_2_ (1 mL); For **15**: *t*-BuOCl (0.15 mmol, 17 μL) in dry CH_2_Cl_2_ (0.2 mL) was added dropwise into a stirred solution of **5** (0.06 mmol, 21 mg) and dry Et_3_N (0.12 mmol, 16 μL) in dry CH_2_Cl_2_ (1 mL). The subsequent steps were identical for both compounds: the mixtures were stirred at -20 ℃ for 12 h, quenched with water, and extracted with CH_2_Cl_2_ (3 × 15 mL). The combined organic layers were washed with brine, dried over Na_2_SO_4_, filtered, and concentrated under reduced pressure. The crude product was treated with 5 M methanolic HCl (1mL) at 0 ℃ under argon atmosphere and stirred for 12 h. After concentration under reduced pressure, the products were purified by recrystallization. The structure was confirmed by NMR (Supplementary Fig. 29-30).

### NMR

NMR measurements were recorded on a Bruker Ascend^TM^ 400 MHz spectrometer. Chemical shifts were referenced to the residual solvent signals of CD_3_OD.

### Data availability

Nucleotide sequences for genes described and used in this study are provided in Supplementary Table 2. The raw transcriptome data have been deposited in the National Genomics Data Center Bioproject database with accession number PRJCA035431.

## Acknowledgements

We are grateful to the Shenzhen Synthetic Biology Infrastructure for instrument support and technical assistance. This research was supported by the National Key R&D Program of China (2023YFA0915500) and Shenzhen Science and Technology Program (KJZD20230923115906013 to Y.J. and JCYJ20220818100804010 to Y.G.).

## Author contributions

D.C. and K.L. performed the experimental work and data analysis. H.W. conducted structural modeling and molecular dynamics simulations. Z.N. and J.C. synthesized and characterized the chemical standards. M.Y. performed transcriptome sequencing and analysis. D.C., Y.G., and Y.J. wrote the manuscript. Q.Y., Y.G., and Y.J. conceived and supervised the project. All authors participated in discussions and manuscript review.

