## Supplementary for "Collective biosynthesis of plant spirooxindole alkaloids through enzyme discovery and engineering"

Supplementary Fig. 26. LC-MS and LC-MS/MS analysis of fluorinated and deuterated tetrahydroalstonine (**5-5e**). 32
Supplementary Fig. 27. LC-MS and LC-MS/MS analysis of fluorinated and deuterated 3-dehydro-tetrahydroalstonine (**15-15e**). 33
Supplementary Fig. 28. LC-MS and LC-MS/MS analysis of fluorinated and deuterated akuammigine (**6-6e**). 34
Supplementary Fig. 29. $^{1}$H-NMR spectra of 3-dehydro-Ajmalicine (**16**) in $CD_3OD$. 35
Supplementary Fig. 30. $^{13}$C-NMR spectra of 3-dehydro-Ajmalicine (**16**) in $CD_3OD$. 36
**3. Supplementary Tables 37**
Supplementary Table 1. Chemical standards used in this study. 37
Supplementary Table 2. Nucleotide sequences for genes described and used in this study. 38
Supplementary Table 3. Primers used for genes cloning in this study. 49
Supplementary Table 4. Previously characterized MDRs used to construct phylogenetic tree in Supplementary Fig. 9. 53
**4. Supplementary References 53**

### 1. Supplementary Materials and Methods

#### Chemicals

All chemicals and analytical standards used in this study were of analytical grade or higher purity (⩾98%). A list of analytical standards is provided in Supplementary Table 1.

#### Molecular biology reagents and kits

Oligonucleotide primers were synthesized by Tsingke Biotech (Beijing, China). Gene amplifications were performed using 2× Hieff Canace AdvanceFast PCR Master Mix polymerase from Yeasen (Shanghai, China). Restriction enzymes were purchased from New England BioLabs (Beverly, MA, US). DNA fragments were assembled using the Hieff Clone Universal II One Step Cloning Kit from Yeasen. PCR reactions were carried out in a ProFlex PCR system from Thermo Fisher Scientific (Waltham, MA, US). DNA fragments were purified using a Gel Extraction Kit from Omega Bio-tek (Norcross, GA, US). DNA and protein concentrations were measured using a Nanodrop One Spectrophotometer from Themo Fisher Scientific. Plasmid DNA was isolated using a Plasmid Mini Kit I from Omega Bio-tek. DNA sequencing was performed by Tsingke Biotech.

#### Plant materials

*Nicotiana benthamiana* plants were grown from seeds in an Illumination incubator (Ningbo Jiangnan Instrument Factory, China) under controlled conditions (25 °C, 16/8 h light/dark cycle, 60% relative humidity). Plants were regularly irrigated with water to maintain optimal growth conditions. Four-week-old plants were utilized for *Agrobacterium tumefaciens*-mediated transformation experiments.

Two-year-old *Uncaria rhynchophylla* seedlings were purchased from Jianhe county, Qiandongnan Miao and Dong Autonomous Prefecture, Guizhou Province, China (26°20′-26°55′ N, 108°17′-109°04′E). One-year-old *Rauvolifa verticillata* seedlings were obtained from Pingnan County, Guigang city, Guangxi Zhuang Autonomous Region, China (23°2′-24°2′ N, 110°3′-110°39′E).

#### Bacterial strains

*Escherichia coli* Top10 and *E. coli* BL21 (DE3) chemically competent cells were purchased from AlpalifeBio (Shenzhen, China). *E. coli* was grown in Luria-Bertani (LB) medium (Guangdong Huankai Microbial, Guangzhou, China). *Agrobacterium tumefaciens* GV3101 chemically competent cells were obtained from Tsingke Biotech. *Saccharomyces cerevisiae* BY4741 were obtained from Beijing Zoman Biotechnology (Beijing, China). *S. cerevisiae* was cultivated and maintained on yeast extract peptone dextrose (YPD, Thermo Fisher Scientific) or synthetic complete (SC) medium (Coolaber, Beijing, China) at 28 °C.

#### Expression and purification of proteins in *E. coli*

For protein expression, sequence-verified constructs were transformed into *E. coli* BL21 (DE3) cells using standard by heat shock transformation. Positive transformants were confirmed by colony PCR and cultured in 5 mL LB medium supplemented with 100 μg/mL ampicillin overnight at 37 °C with a shaking of 200 rpm. The overnight cultures were mixed with an equal volume of 50% glycerol and stored at -80 °C as glycerol stocks.

For production of recombinant enzymes in *E. coli*, glycerol stocks were used to inoculate 2 mL LB medium containing 100 μg/mL ampicillin. The cultures were grown overnight at 37 °C with a shaking of 200 rpm. The overnight cultures were then used to inoculate 100 mL LB medium containing 100 μg/mL ampicillin and incubated at 37 °C with shaking at 220 rpm until reaching an optical density ($OD_{600}$) of 0.6-0.8. Protein expression was induced by cooling the cultures to 16 °C and adding isopropyl β-D-1-thiogalactopyranoside (IPTG) to a final concentration of 200 μM. After 16-18 h of induction, cells were harvested by centrifugation (4,000 rpm, 4 °C, 20 min) and resuspended in 2.5 mL B-PER Complete Bacterial Protein Extraction Reagent (Thermo Fisher Scientific). The cell suspension was incubated at room temperature for 30 min with gentle shaking to facilitate lysis. The bacterial lysate was cleared by centrifugation (16,000 rpm, 4 °C, 20 min), and the supernatant was transferred to a 15 mL centrifuge tube. Subsequently, 150 μL of HisSep Ni-NTA Agarose Resin (Yeasen) was added to the cleared lysate and gently shaken for 1 hour at 4 °C. The resin was collected by centrifugation (1,000 rpm, 4 °C, 1 min) and washed three times with buffer A1 (50 mM Tris-HCl pH 8.0, 50 mM glycine, 500 mM NaCl, 5% glycerol, 20 mM imidazole). His-tagged proteins were eluted twice with 600 μL Buffer B (50 mM Tris-HCl pH 8; 50 mM glycine; 500 mM NaCl; 5% glycerol; 500 mM imidazole). The eluted proteins were buffer-exchanged into Buffer A4 (20 mM HEPES pH 7.5, 150 mM NaCl) using Amicon® Ultra Centrifugal Filter units (Merck Millipore). The purified proteins were aliquoted in 100 μL, snap-frozen in liquid nitrogen, and stored at -80 °C until further use. Protein concentrations were determined using a Nanodrop Spectrophotometer, and protein purity was assessed by SDS-PAGE analysis.

**Enzymatic preparation for 20*R*-corynantheidine**

Reaction mixtures (500 μL total volume in 50 mM HEPES, pH 7.4) were composed of 20 μg of each enzyme (CrSTR, CrSGD, CpDCS, and MsEnolMT), 400 μM NADPH, 200 μM SAM, 200 μM ascorbate, 40 μM secologanin, and 40 μM tryptamine[1]. The reactions were incubated at 30 °C for 16 h and then quenched with equal volumes of ethyl acetate. The supernatant was concentrated using a rotary evaporator, and the resulting residue was resuspended in 50 μL methanol as the 20*R*-corynantheidine substrate.

### 2. Supplementary Figures

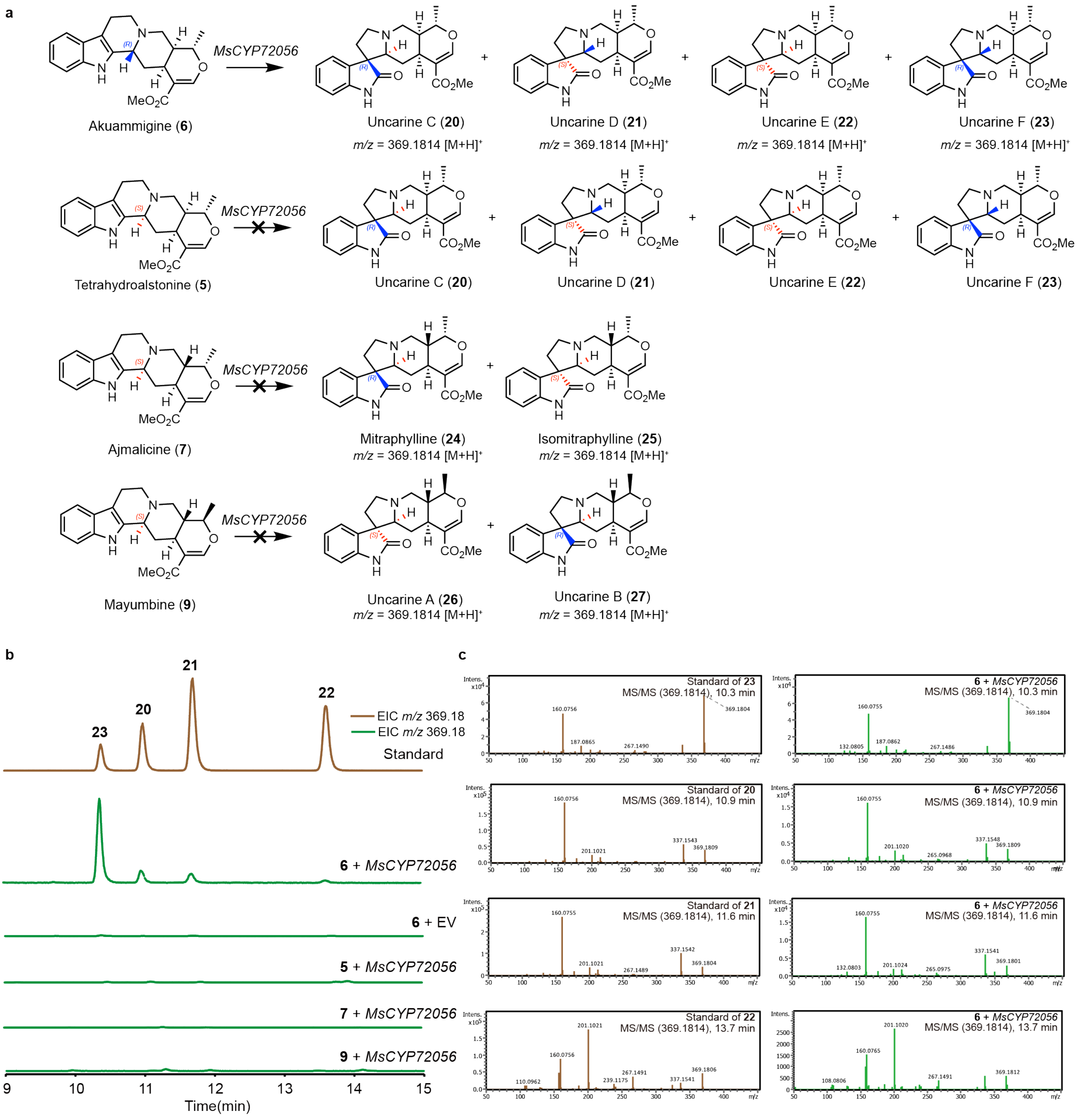

**Supplementary Fig. 1. Substrate specificity of MsCYP72056 in N. benthamiana expression system.** (**a**) Biosynthetic pathway depicting the conversion of pentacyclic alkaloids (**5, 6, 7, 9**) to pentacyclic spirooxindole alkaloids isomers (**20-27**). (**b**) Extracted ion chromatograms of *m/z* corresponding to pentacyclic spirooxindole alkaloids (**20-27**). Brown trace: authentic standards; green traces: compounds from transient expression of *MsCYP72056* with **5, 6, 7, 9** as substrate in *N. benthamiana*. (**c**) MS/MS spectra comparing authentic standards (**20-23**) with in planta-generated products.

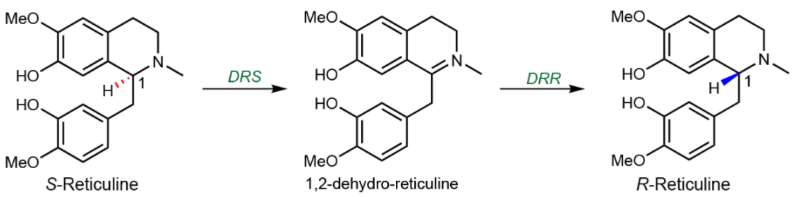

**Supplementary Fig. 2. Two-step stereochemical conversion of S-reticuline to R-reticuline catalyzed by 1,2-dehydroreticuline synthase (DRS) and 1,2-dehydroreticuline reductase (DRR).** DRS is a cytochrome P450 enzyme, while DRR is an aldo-keto reductase enzyme.

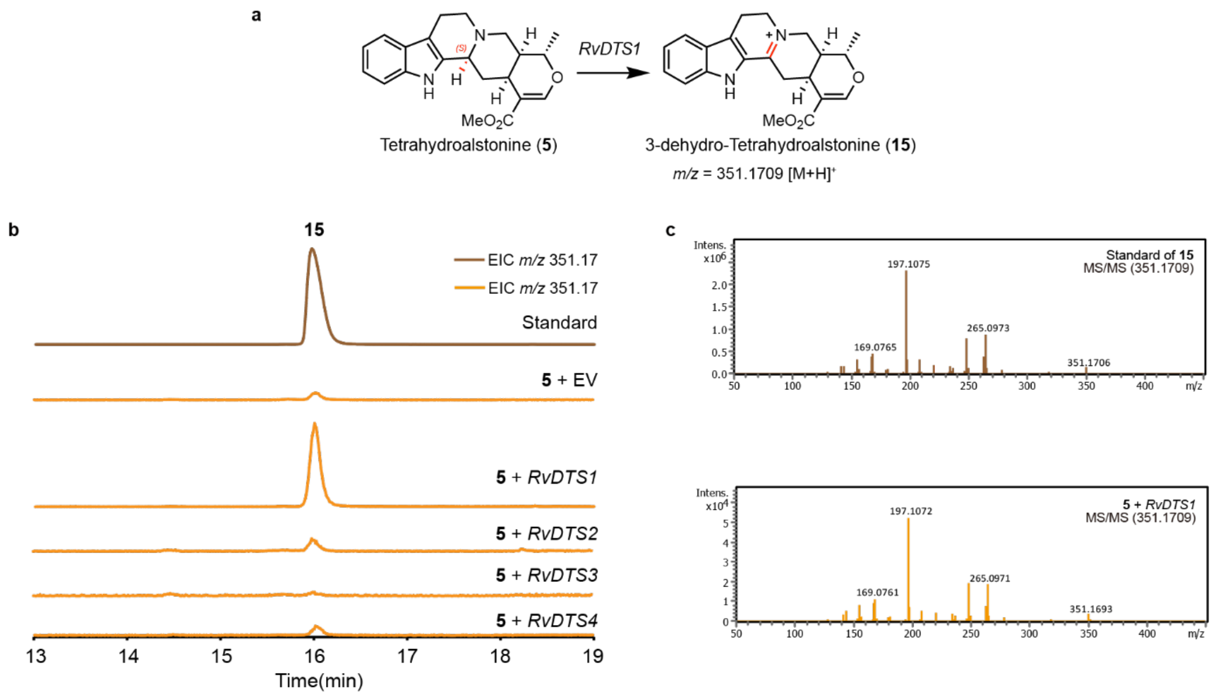

**Supplementary Fig. 3. LC-MS and LC-MS/MS analysis of 3-dehydro-tetrahydroalstonine (15).** (**a**) Biosynthetic pathway depicting the conversion of tetrahydroalstonine (**5**) to 3-dehydro-tetrahydroalstonine (**15**). (**b**) Extracted ion chromatograms of *m/z* corresponding to **15**. Brown trace: authentic standard; orange traces: compound from transient expression of four *RvDTS* candidate genes with **5** as substrate in *N. benthamiana*. (**c**) MS/MS spectra comparing authentic standard (**15**) with in planta-generated product.

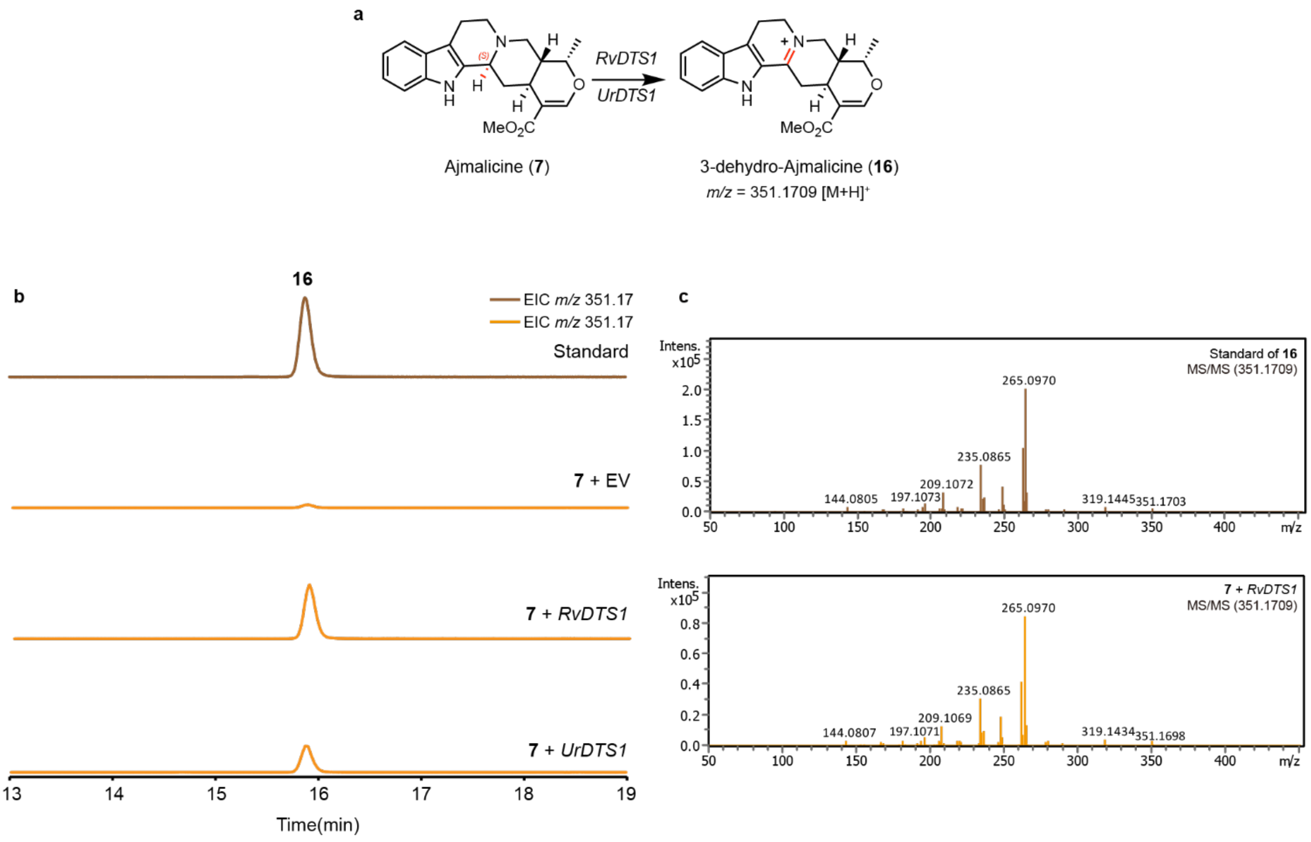

**Supplementary Fig. 4. LC-MS and LC-MS/MS analysis of 3-dehydro-ajmalicine (16).** (**a**) Biosynthetic pathway depicting the conversion of ajmalicine (**7**) to 3-dehydro-ajmalicine (**16**). (**b**) Extracted ion chromatograms of *m/z* corresponding to **16**. Brown trace: authentic standard; orange traces: compound from transient expression of *RvDTS1* or *UrDTS1* with **7** as substrate in *N. benthamiana*. (**c**) MS/MS spectra comparing authentic standard (**16**) with in planta-generated product.

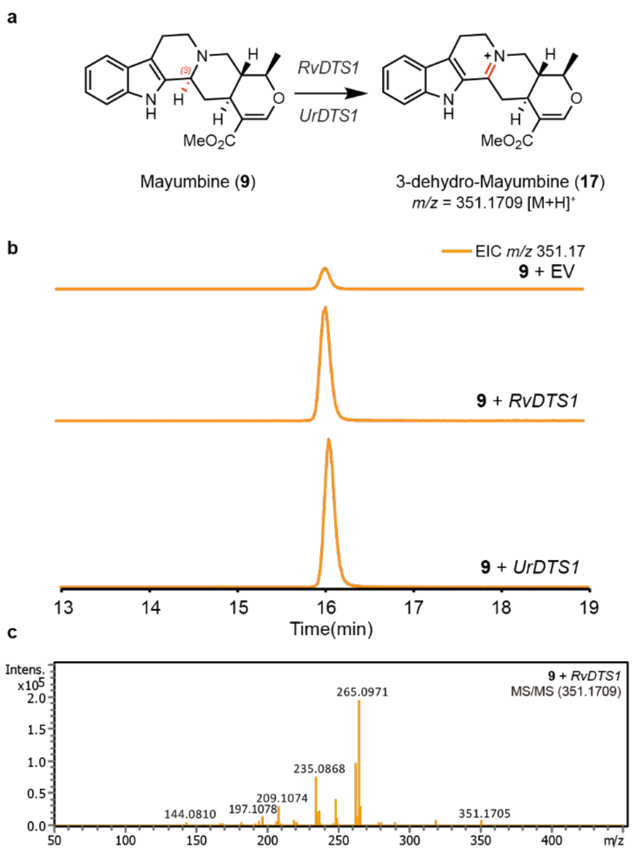

**Supplementary Fig. 5. LC-MS and LC-MS/MS analysis of 3-dehydro-mayumbine (17).** (**a**) Biosynthetic pathway depicting the conversion of mayumbine (**9**) to 3-dehydro-mayumbine (**17**). (**b**) Extracted ion chromatograms of *m/z* corresponding to **17**. Orange traces: compound from transient expression of *RvDTS1* or *UrDTS1* with **9** as substrate in *N. benthamiana*. (**c**) MS/MS spectra of in planta-generated product.

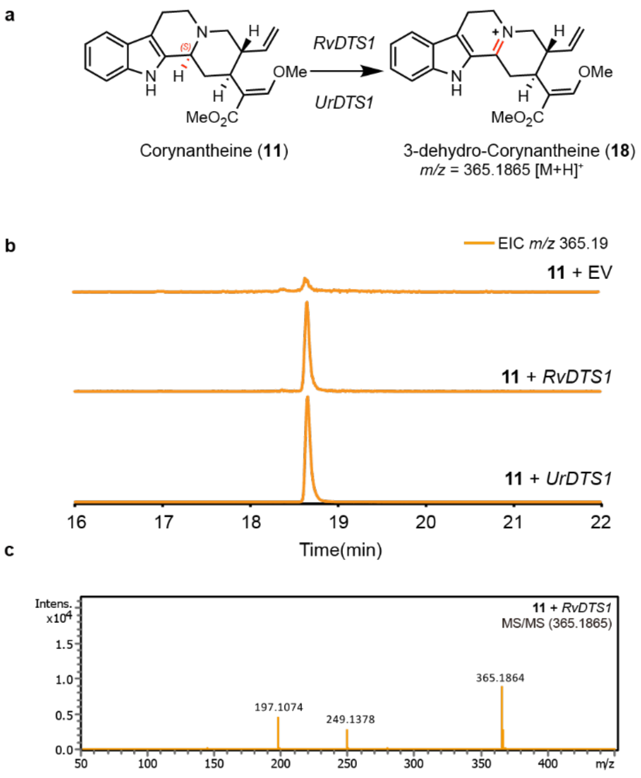

**Supplementary Fig. 6. LC-MS and LC-MS/MS analysis of 3-dehydro-corynantheine (18).** (**a**) Biosynthetic pathway depicting the conversion of corynantheine (**11**) to 3-dehydro-corynantheine (**18**). (**b**) Extracted ion chromatograms of *m/z* corresponding to **18**. Orange traces: compound from transient expression of *RvDTS1* or *UrDTS1* with **11** as substrate in *N. benthamiana*. (**c**) MS/MS spectra of in planta-generated product.

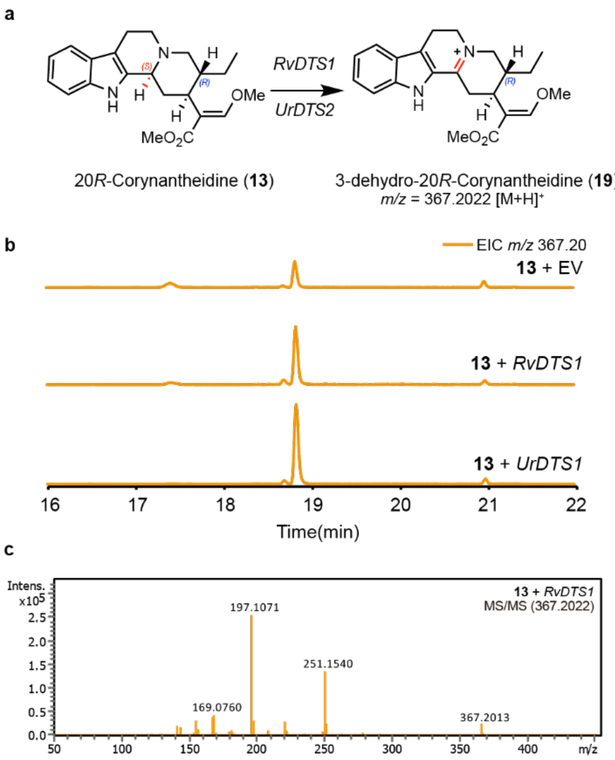

**Supplementary Fig. 7. LC-MS and LC-MS/MS analysis of 3-dehydro-20R-corynantheidine (19).** (**a**) Biosynthetic pathway depicting the conversion of 20*R*-corynantheidine (**13**) to 3-dehydro-20*R*-corynantheidine (**19**). (**b**) Extracted ion chromatograms of *m/z* corresponding to **19**. Orange traces: compound from transient expression of *RvDTS1* or *UrDTS1* with **13** as substrate in *N. benthamiana*. (**c**) MS/MS spectra of in planta-generated product.

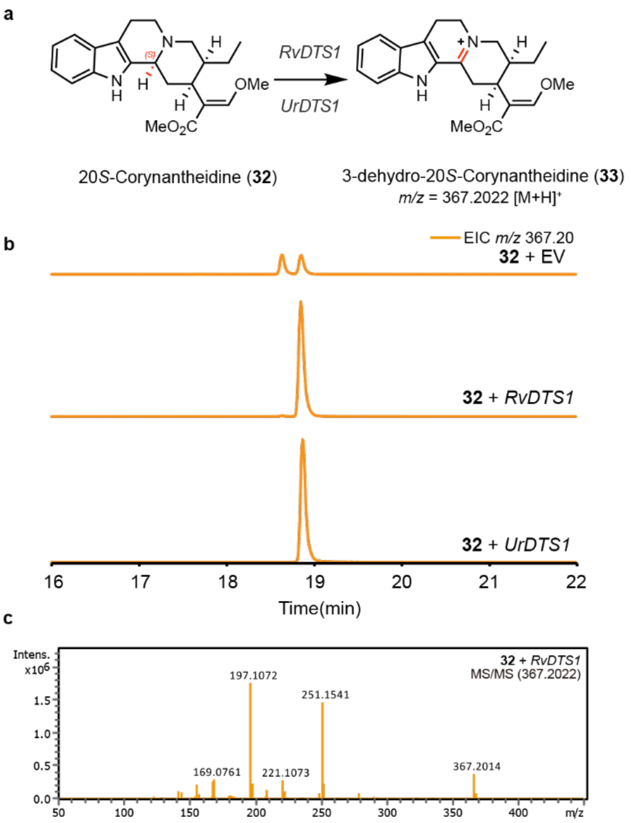

**Supplementary Fig. 8. LC-MS and LC-MS/MS analysis of 3-dehydro-20S-corynantheidine (33).** (**a**) Biosynthetic pathway depicting the conversion of 20*S*-corynantheidine (**32**) to 3-dehydro-20*S*-corynantheidine (**33**). (**b**) Extracted ion chromatograms of *m/z* corresponding to **33**. Orange traces: compound from transient expression of *RvDTS1* or *UrDTS1* with **32** as substrate in *N. benthamiana*. (**c**) MS/MS spectra of in planta-generated product.

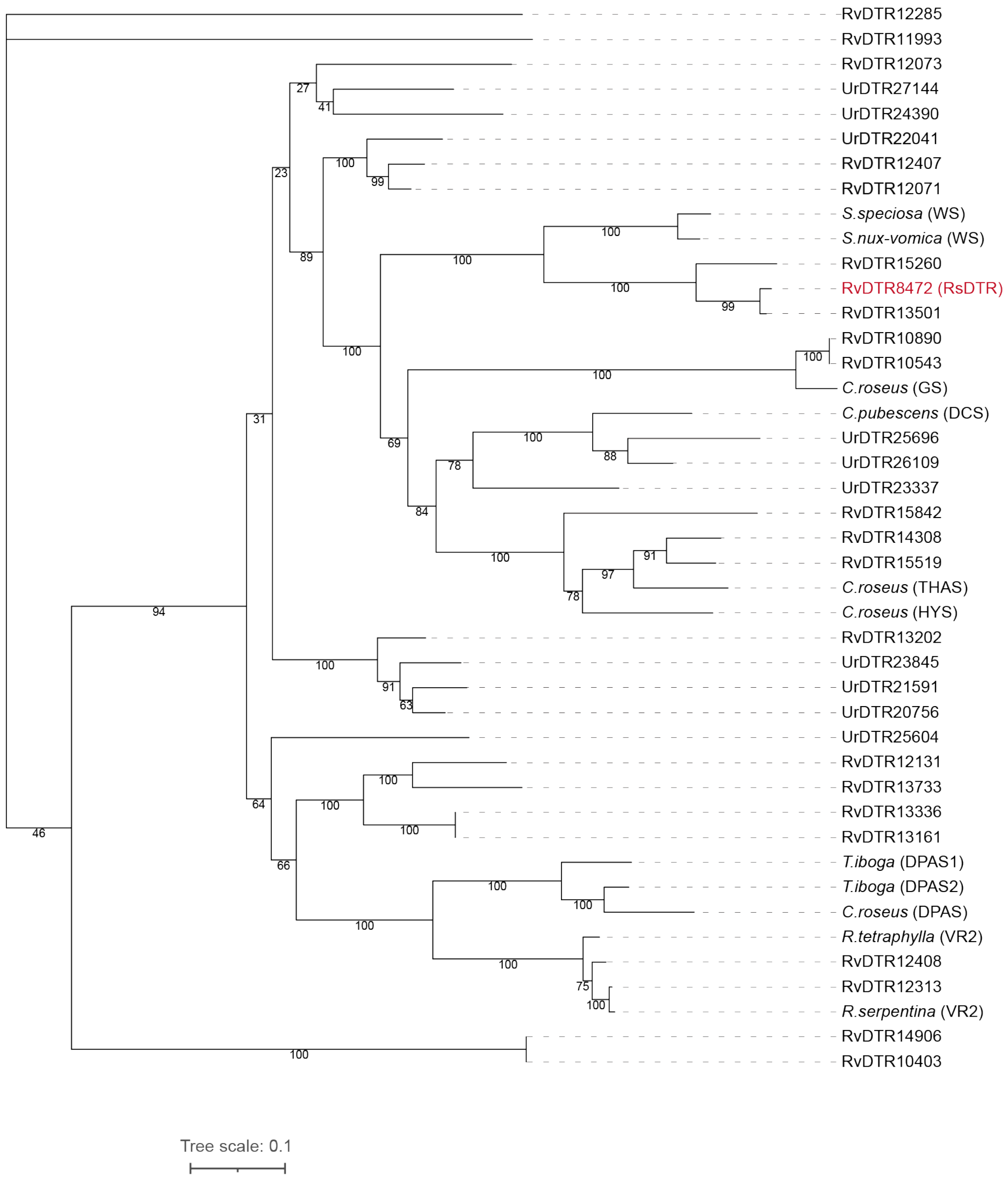

**Supplementary Fig. 9. Phylogenetic tree of MDRs in *R. verticillata* and *U. rhynchophylla* with previously characterized MDRs in MIA biosynthetic pathway.** Previously characterized MDRs used in this phylogenetic tree were shown in Supplementary Table 4.

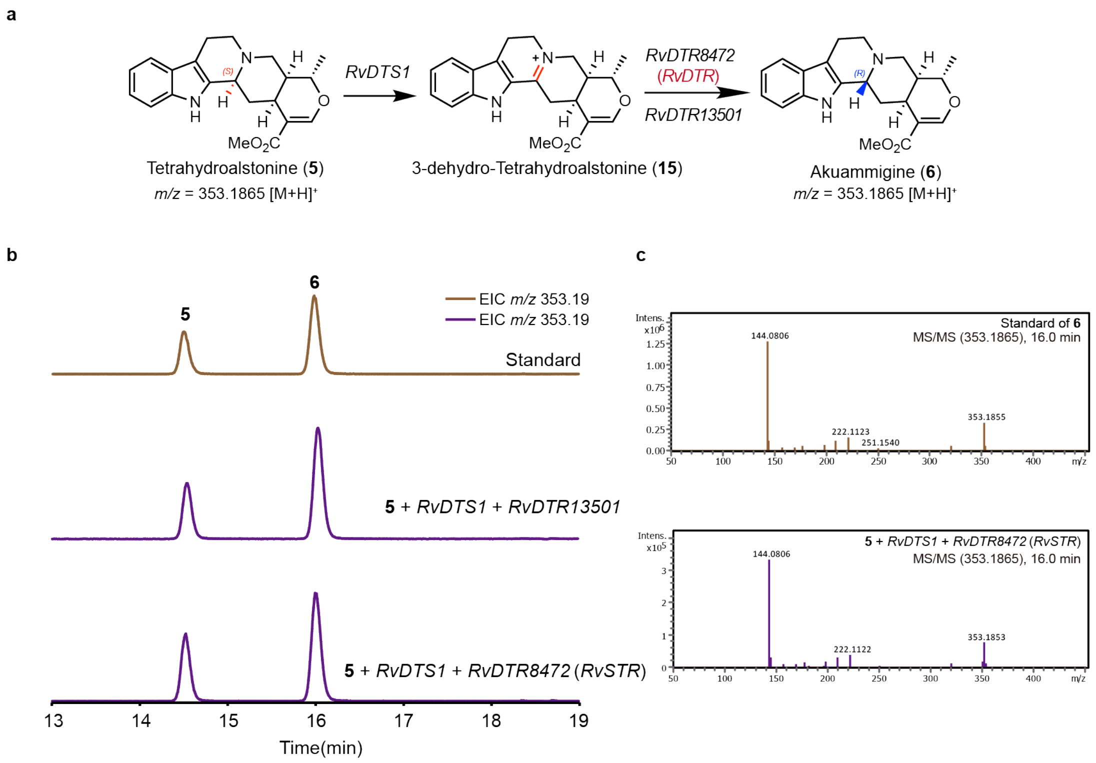

**Supplementary Fig. 10. LC-MS and LC-MS/MS analysis of *RvDTR* candidate genes.** (**a**) Biosynthetic pathway depicting the stepwise conversion of tetrahydrolastonine (**5**) to akuammigine (**6**). (**b**) Extracted ion chromatograms of *m/z* corresponding to **6**. Brown trace: authentic standards; purple traces: compound from transient expression of *RvDTS1*, *RvDTR* candidate genes with **5** as substrate in *N. benthamiana*. (**c**) MS/MS spectra comparing authentic standard (**6**) with in planta-generated product.

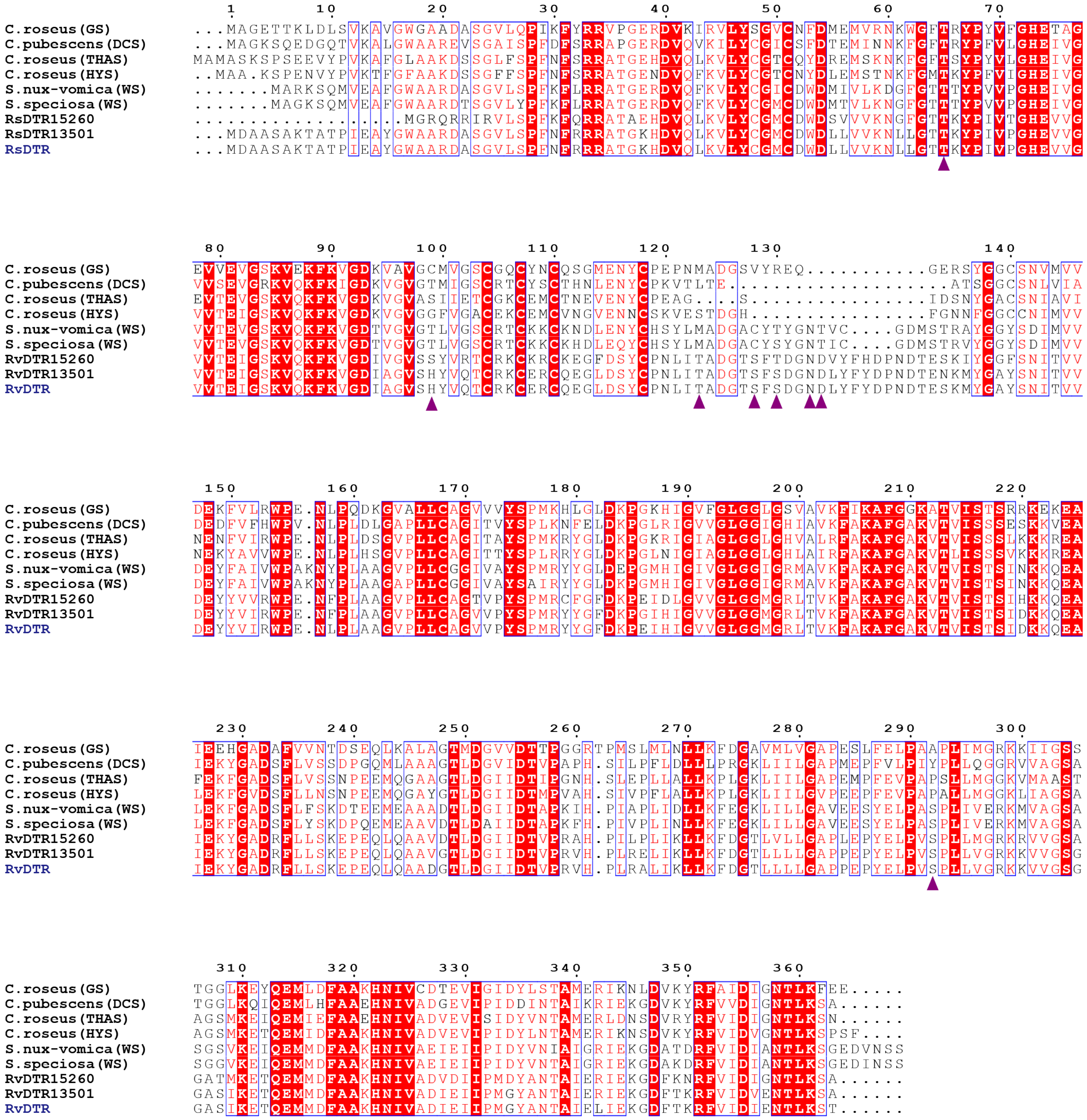

**Supplementary Fig. 11. Amino acid alignment of MDRs used in this study.** Substrate binding sites are marked with purple triangles.

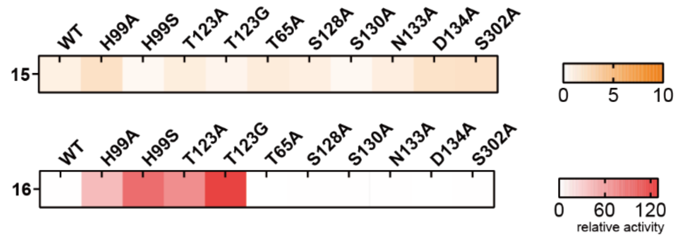

**Supplementary Fig. 12. Relative activities of wild-type RvDTR and its mutants toward different 3-dehydro substrates.** The color intensity corresponds to product formation (normalized to wild-type activity).

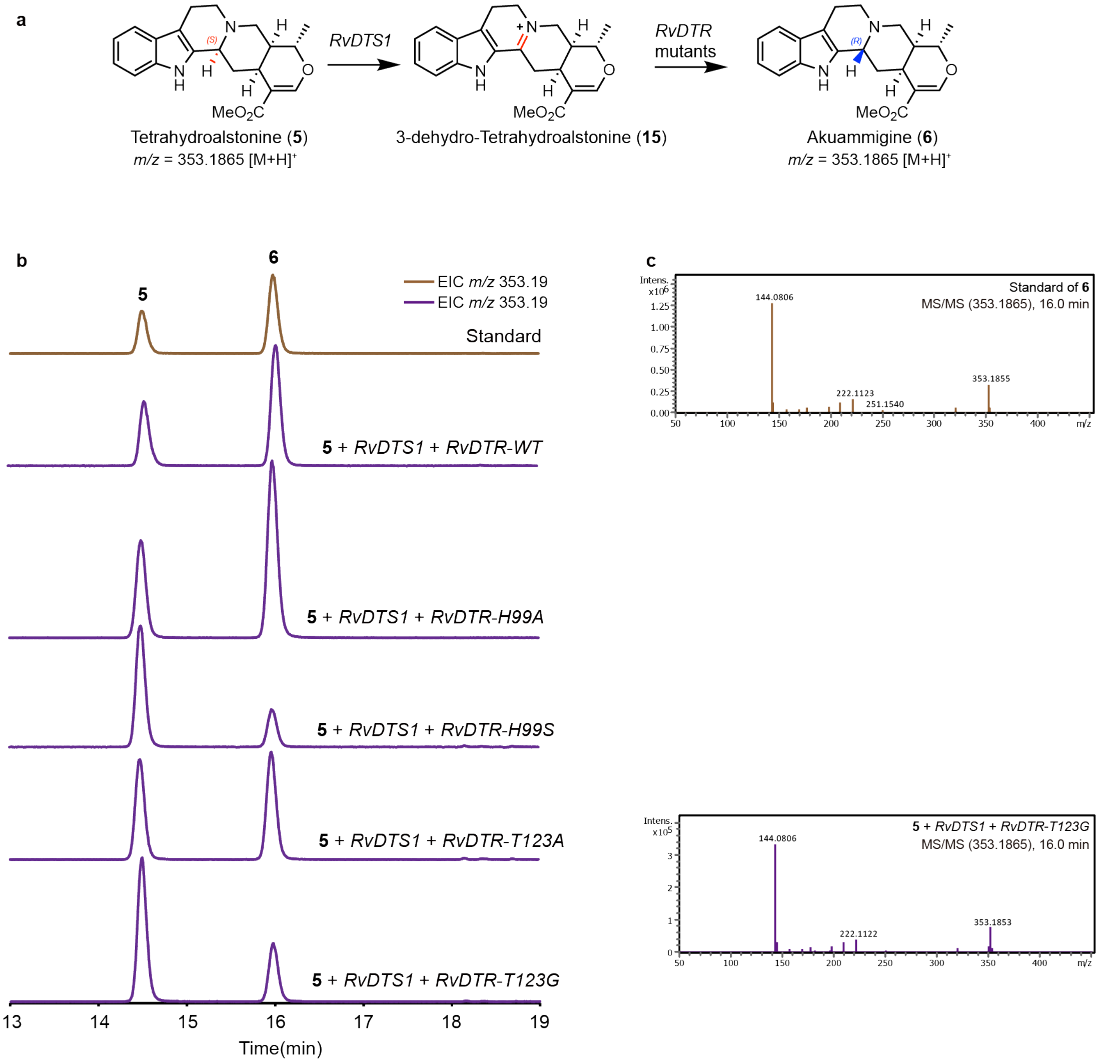

**Supplementary Fig. 13. LC-MS and LC-MS/MS analysis of akuammigine (6).** (**a**) Biosynthetic pathway depicting the stepwise conversion of tetrahydrolastonine (**5**) to akuammigine (**6**). (**b**) Extracted ion chromatograms of *m/z* corresponding to **6**. Brown trace: authentic standards; purple traces: compound from transient expression of *RvDTS1*, *RvDTR* variants with **5** as substrate in *N. benthamiana*. (**c**) MS/MS spectra comparing authentic standard (**6**) with in planta-generated products.

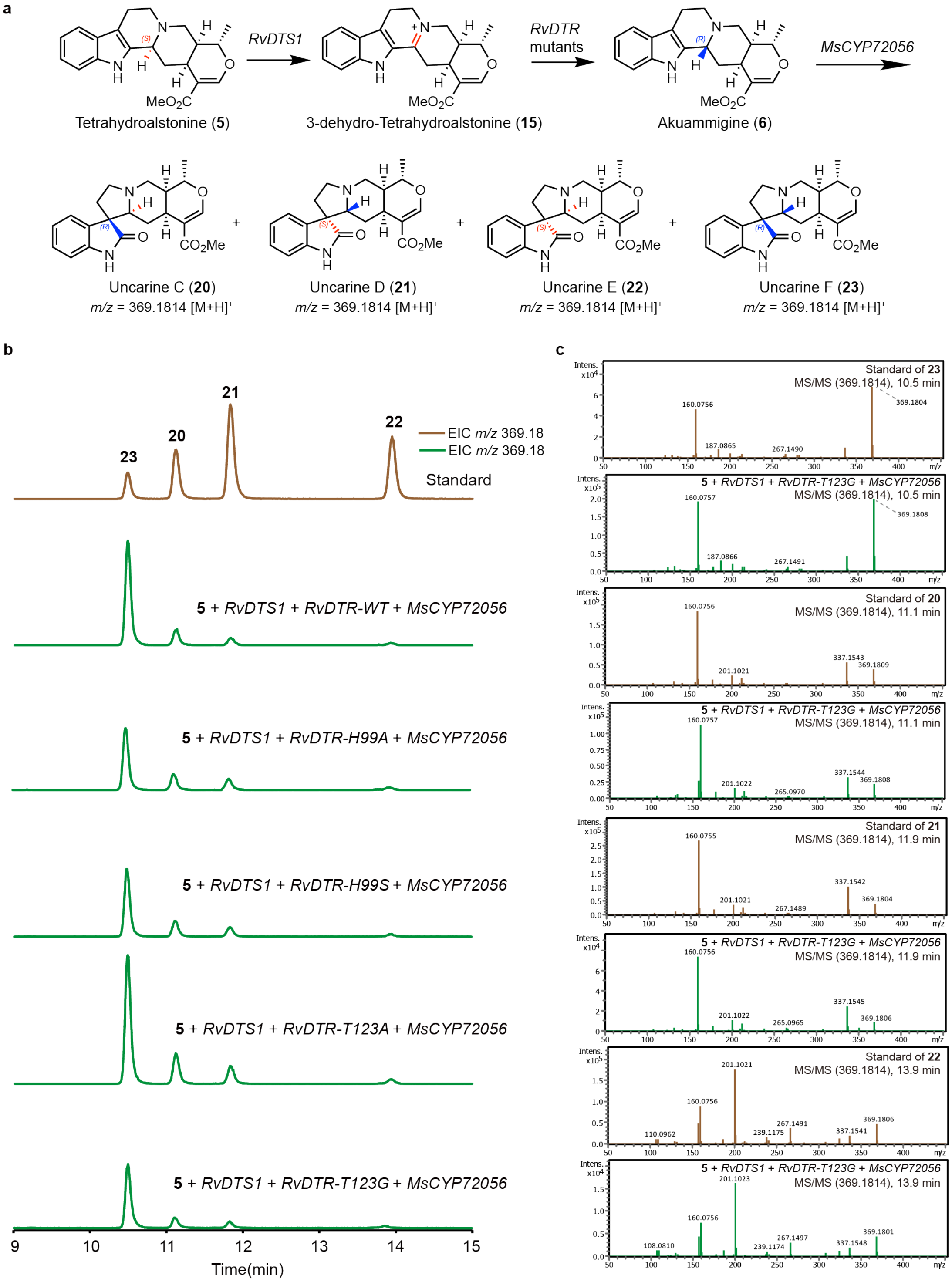

**Supplementary Fig. 14. LC-MS and LC-MS/MS analysis of uncarine isomers (20-23).** (**a**) Biosynthetic pathway depicting the stepwise conversion of tetrahydroalstonine (**5**) to uncarine isomers (**20-23**). (**b**) Extracted ion chromatograms of *m/z* corresponding to uncarine isomers (**20-23**). Brown trace: authentic standards; green traces: compounds from transient expression of *RvDTS1*, *RvDTR* variants, *MsCYP72506* with **5** as substrate in *N. benthamiana*. (**c**) MS/MS spectra comparing authentic standards (**20-23**) with in planta-generated products.

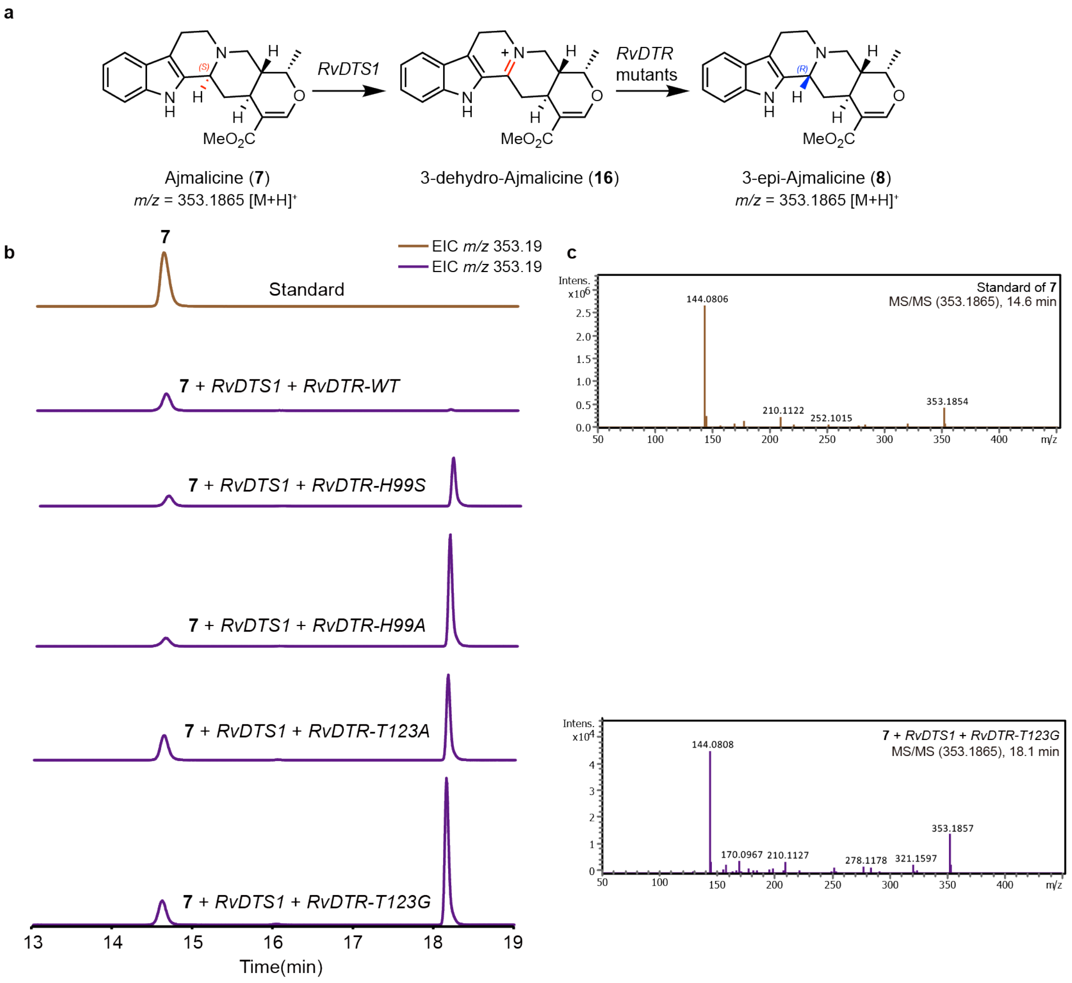

**Supplementary Fig. 15. LC-MS and LC-MS/MS analysis of 3-epi-ajmalicine (8).** (**a**) Biosynthetic pathway depicting the stepwise conversion of ajmalicine (**7**) to 3-epi-ajmalicine (**8**). (**b**) Extracted ion chromatograms of *m/z* corresponding to **8**. Brown trace: authentic standards; purple traces: compound from transient expression of *RvDTS1*, *RvDTR* variants with **7** as substrate in *N. benthamiana*. (**c**) MS/MS spectra comparing authentic standard (**7**) with in planta-generated products. Standard of **8** is unavailable.

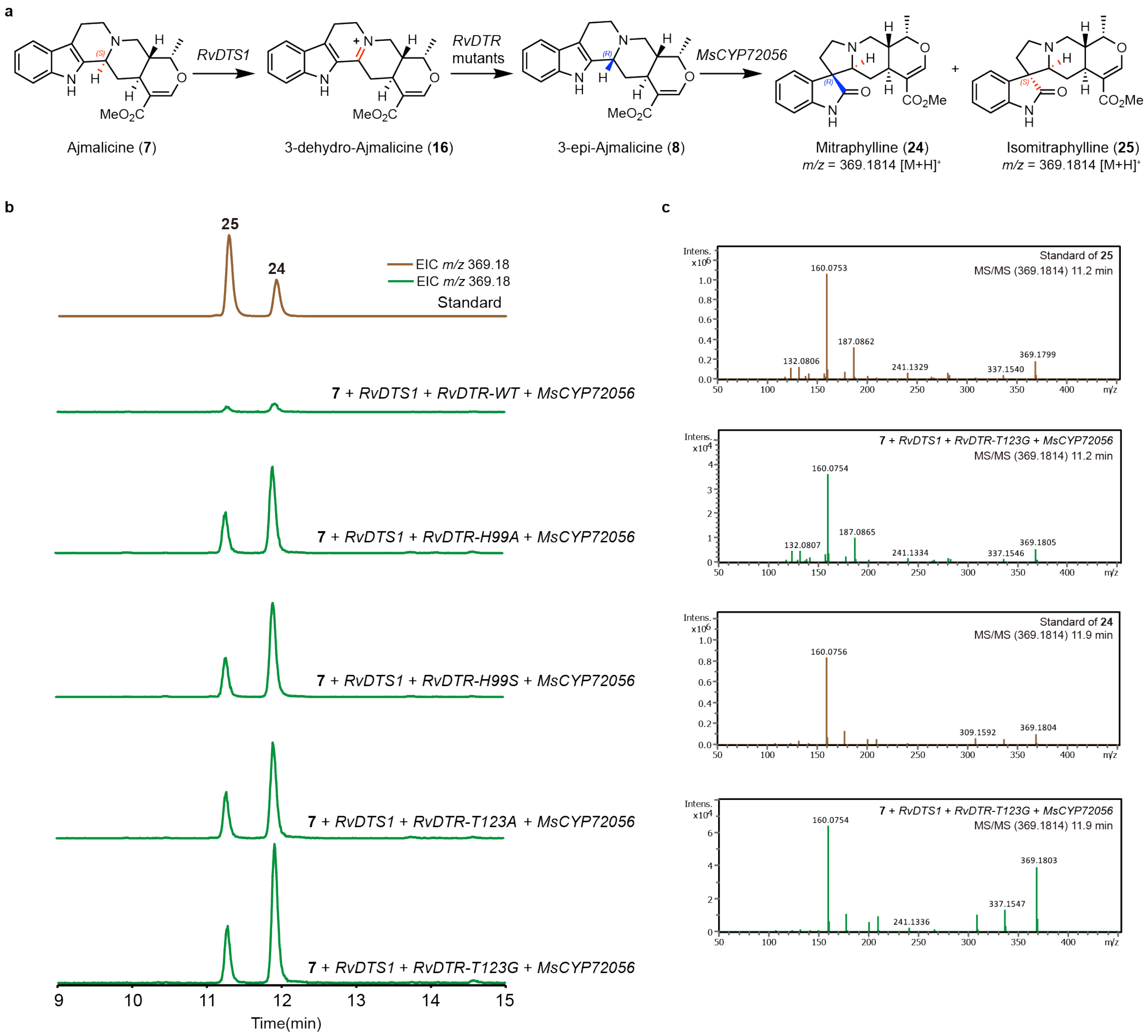

**Supplementary Fig. 16. LC-MS and LC-MS/MS analysis of mitraphylline (24) and isomitraphylline (25).** (**a**) Biosynthetic pathway depicting the stepwise conversion of ajmalicine (**7**) to mitraphylline (**24**) and isomitraphylline (**25**). (**b**) Extracted ion chromatograms of *m/z* corresponding to **24-25**. Brown trace: authentic standards; green traces: compounds from transient expression of *RvDTS1*, *RvDTR* variants, *MsCYP72506* with **7** as substrate in *N. benthamiana*. (**c**) MS/MS spectra comparing authentic standards (**24-25**) with in planta-generated products.

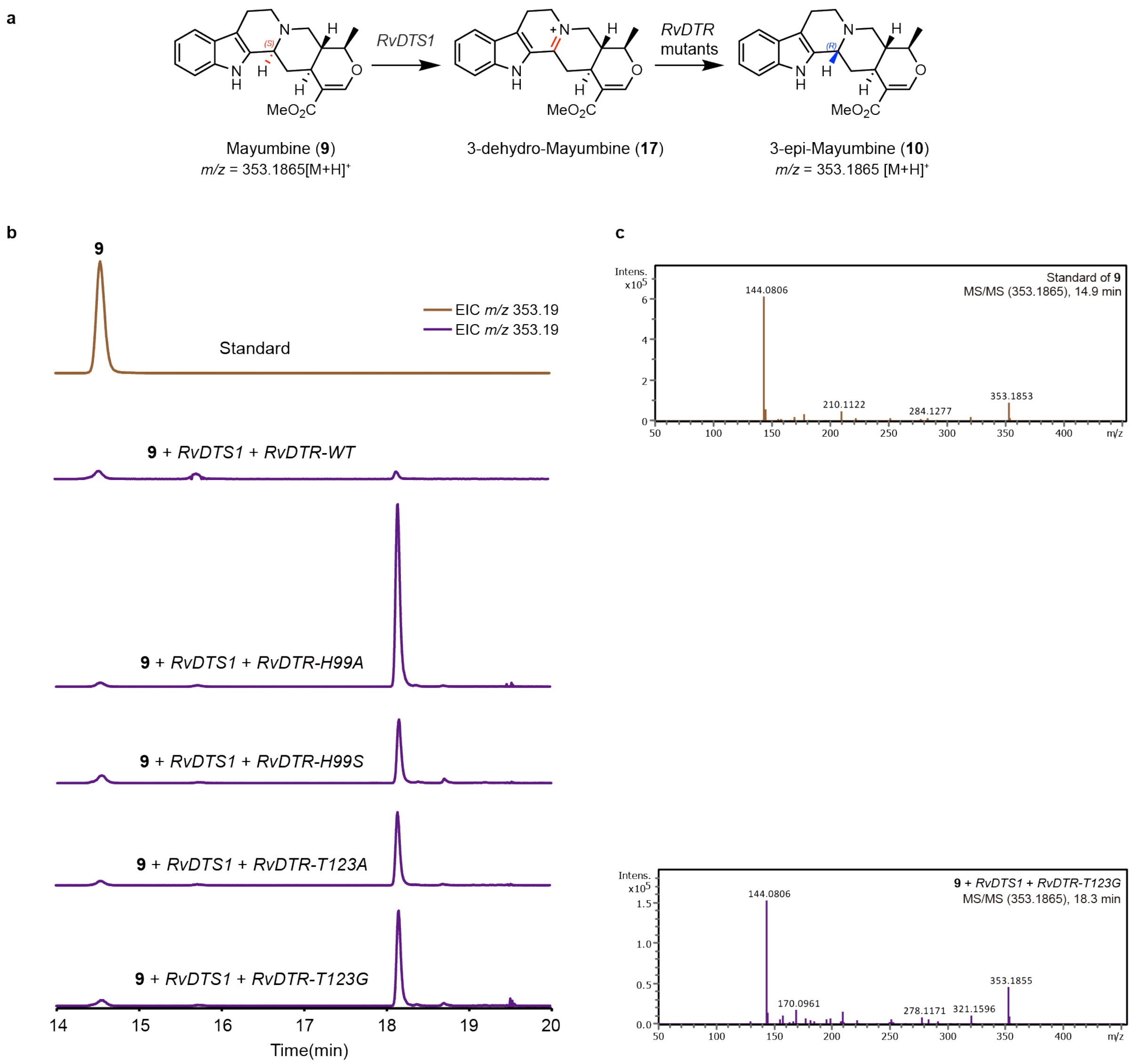

**Supplementary Fig. 17. LC-MS and LC-MS/MS analysis of 3-epi-mayumbine (10).** (**a**) Biosynthetic pathway depicting the stepwise conversion of mayumbine (**9**) to 3-epi-mayumbine (**10**). (**b**) Extracted ion chromatograms of *m/z* corresponding to **10**. Brown trace: authentic standards; purple traces: compound from transient expression of *RvDTS1*, *RvDTR* variants with **9** as substrate in *N. benthamiana*. (**c**) MS/MS spectra comparing authentic standard (**9**) with in planta-generated products. Standard of **10** is unavailable.

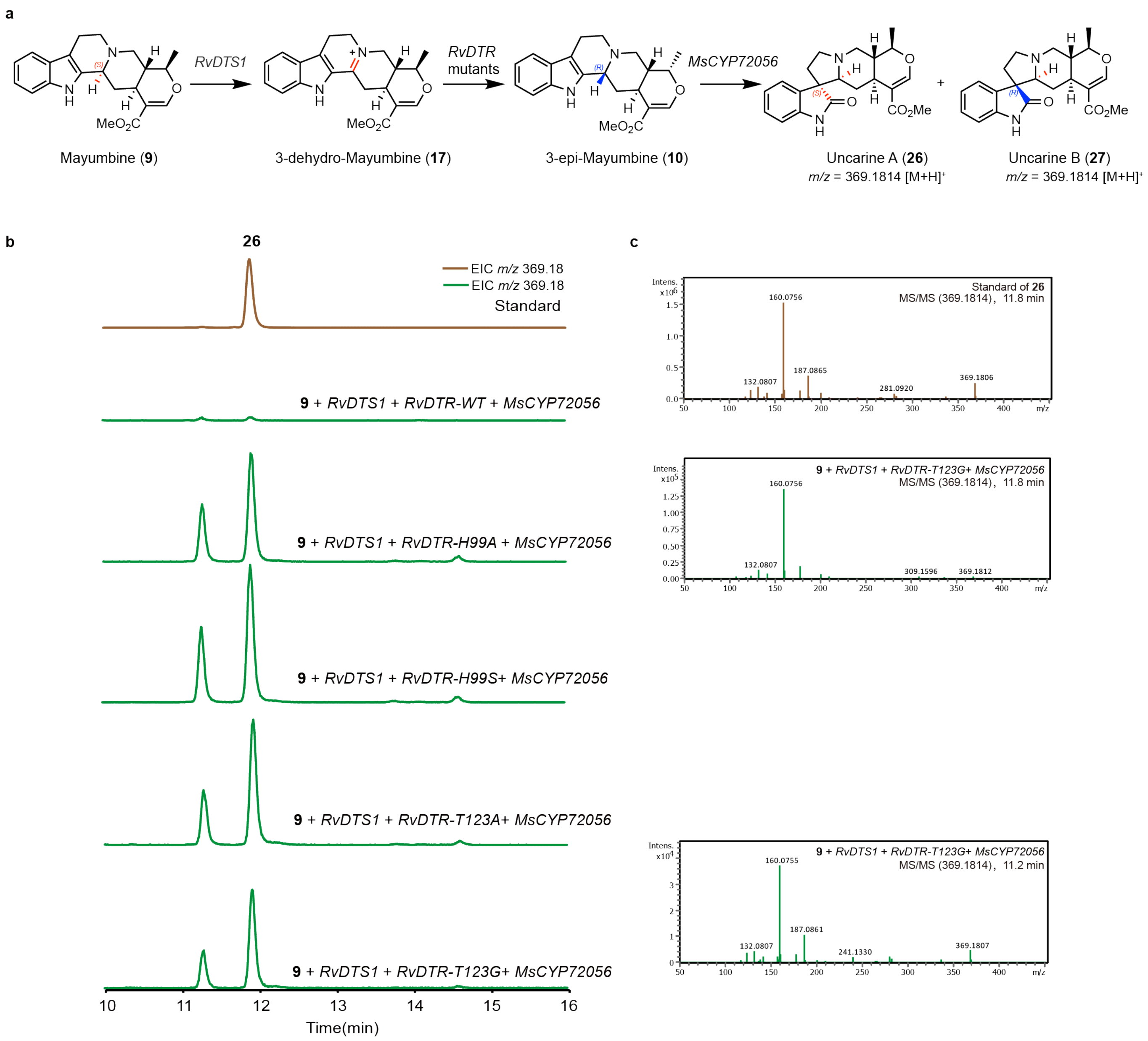

**Supplementary Fig. 18. LC-MS and LC-MS/MS analysis of uncarine isomers (26-27).** (**a**) Biosynthetic pathway depicting the stepwise conversion of mayumbine (**9**) to uncarine isomers (**26-27**). (**b**) Extracted ion chromatograms of *m/z* corresponding to **26-27**. Brown trace: authentic standards; green traces: compounds from transient expression of *RvDTS1*, *RvDTR* variants, *MsCYP72506* with **9** as substrate in *N. benthamiana*. (**c**) MS/MS spectra comparing authentic standard (**26**) with in planta-generated products. Standard of **27** is unavailable.

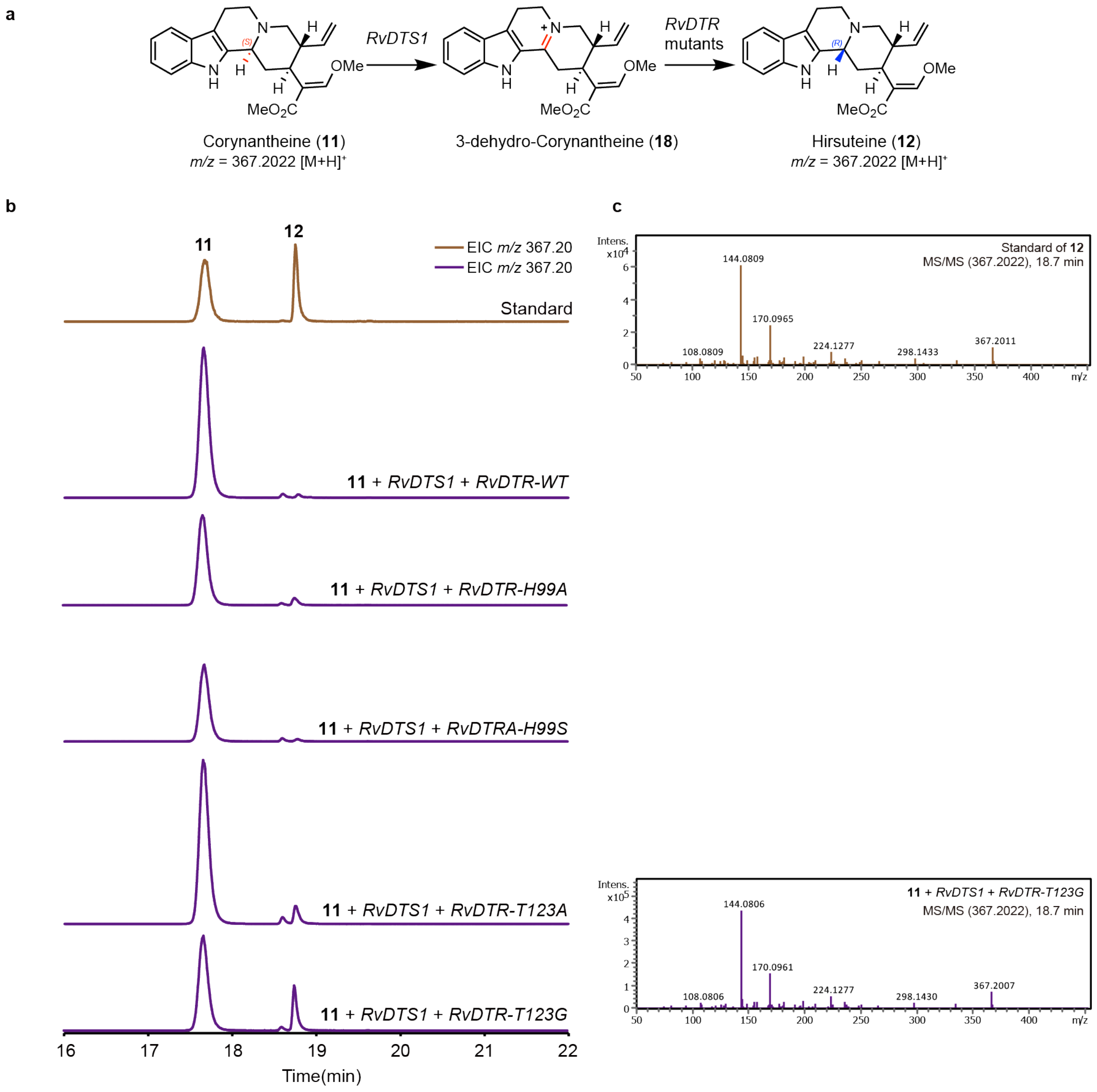

**Supplementary Fig. 19. LC-MS and LC-MS/MS analysis of hirsuteine (12).** (**a**) Biosynthetic pathway depicting the stepwise conversion of corynantheine (**11**) to hirsuteine (**12**). (**b**) Extracted ion chromatograms of *m/z* corresponding to **12**. Brown trace: authentic standards; purple traces: compound from transient expression of *RvDTS1*, *RvDTR* variants with **11** as substrate in *N. benthamiana*. (**c**) MS/MS spectra comparing authentic standard (**12**) with in planta-generated product.

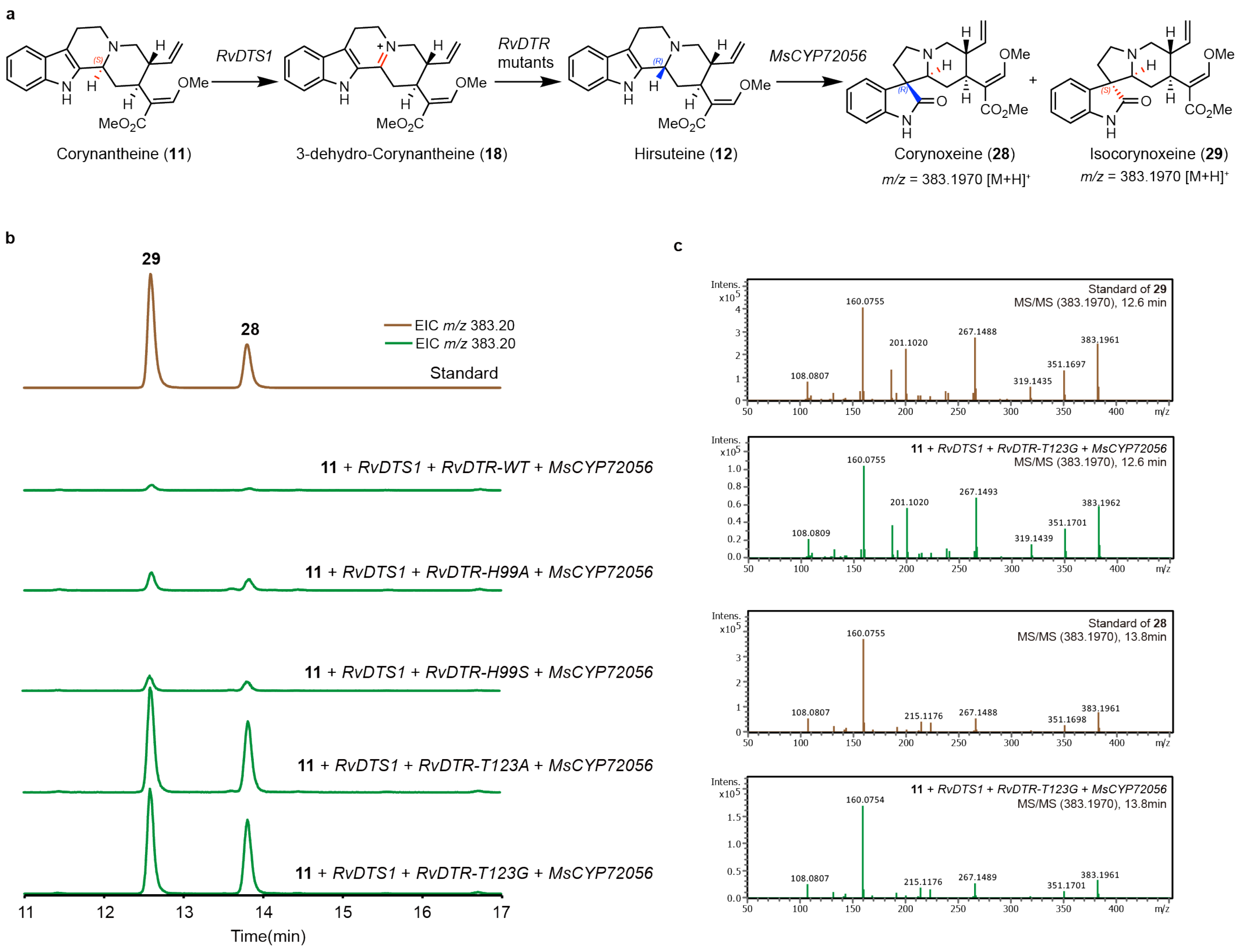

**Supplementary Fig. 20. LC-MS and LC-MS/MS analysis of corynoxeine (28) and isocorynoxeine (29).** (**a**) Biosynthetic pathway depicting the stepwise conversion of corynantheine (**11**) to corynoxeine (**28**) and isocorynoxeine (**29**). (**b**) Extracted ion chromatograms of *m/z* corresponding to **28-29**. Brown trace: authentic standards; green traces: compounds from transient expression of *RvDTS1*, *RvDTR* variants, *MsCYP72506* with **11** as substrate in *N. benthamiana*. (**c**) MS/MS spectra comparing authentic standard (**28-29**) with in planta-generated products.

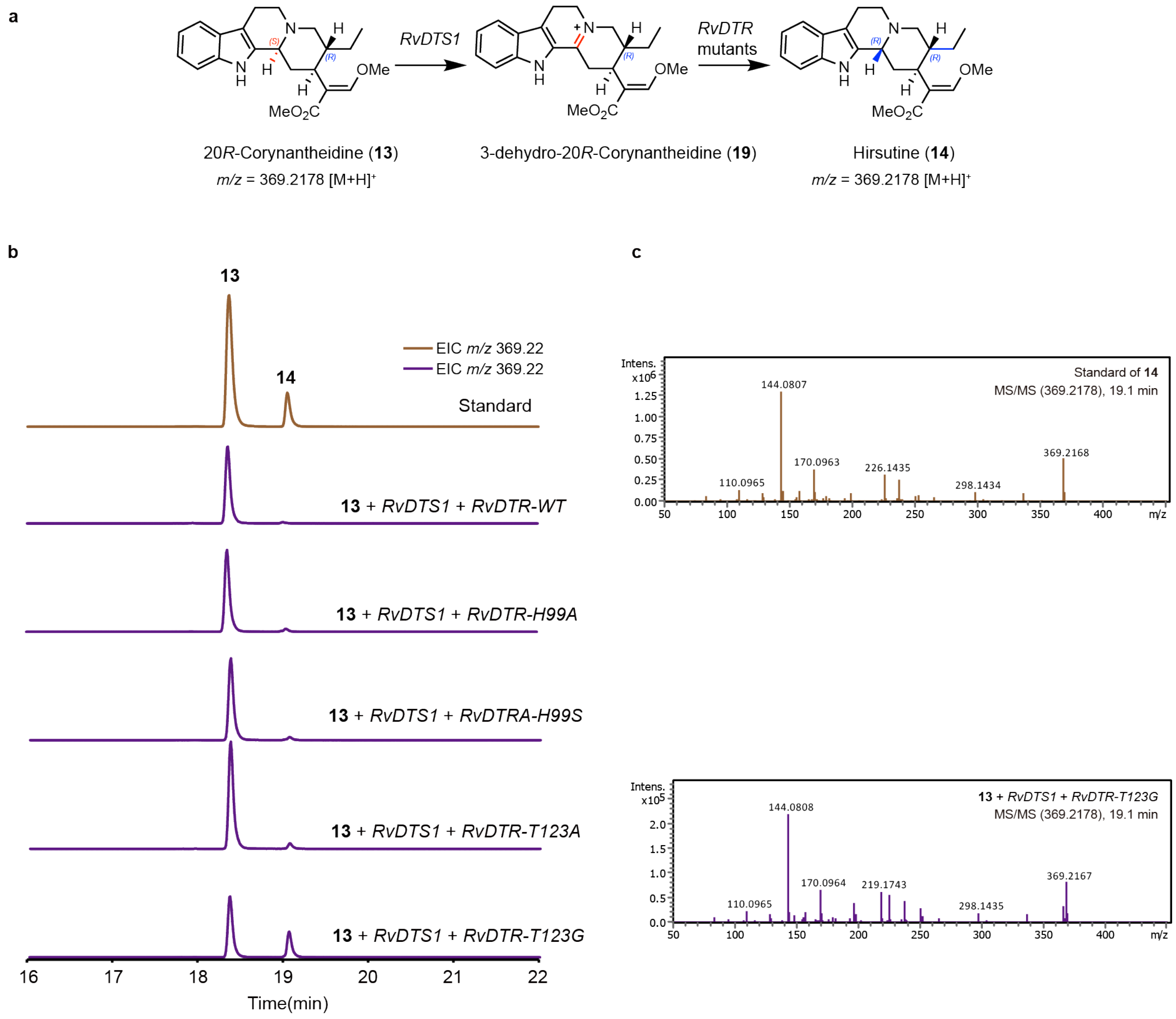

**Supplementary Fig. 21. LC-MS and LC-MS/MS analysis of hirsutine (14).** (**a**) Biosynthetic pathway depicting the stepwise conversion of 20*R*-Corynantheidine (**13**) to hirsutine (**14**). (**b**) Extracted ion chromatograms of *m/z* corresponding to **14**. Brown trace: authentic standards; purple traces: compound from transient expression of *RvDTS1*, *RvDTR* variants with **13** as substrate in *N. benthamiana*. (**c**) MS/MS spectra comparing authentic standard (**14**) with in planta-generated product.

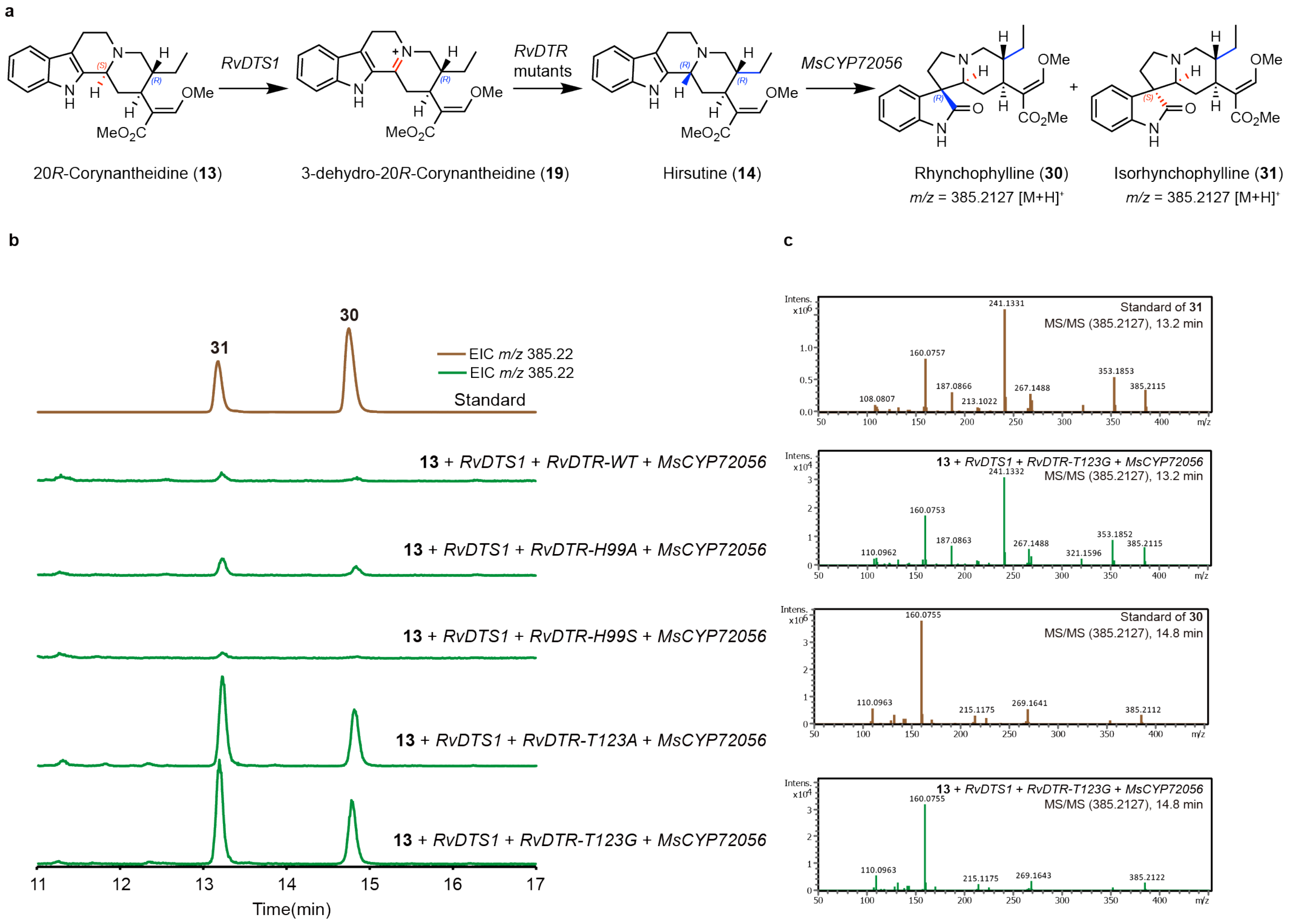

**Supplementary Fig. 22. LC-MS and LC-MS/MS analysis of rhynchophylline (30) and isorhynchophylline (31).** (**a**) Biosynthetic pathway depicting the stepwise conversion of 20*R*-corynantheidine (**13**) to rhynchophylline (**30**) and isorhynchophylline (**31**). (**b**) Extracted ion chromatograms of *m/z* corresponding to **30-31**. Brown trace: authentic standards; green traces: compounds from transient expression of *RvDTS1*, *RvDTR* variants, *MsCYP72506* with **13** as substrate in *N. benthamiana*. (**c**) MS/MS spectra comparing authentic standard (**30-31**) with in planta-generated products.

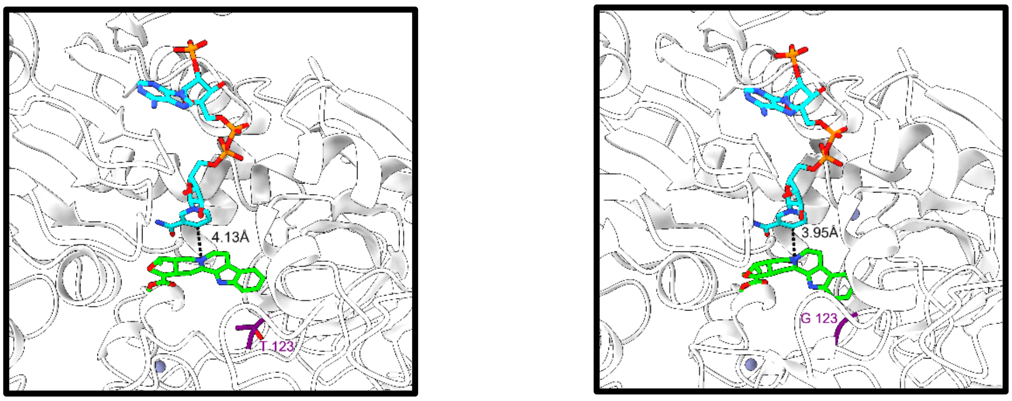

**Supplementary Fig. 23. Close-up views of active pocket of RvDTR-WT (left) and RvDTR-T123G (right).** The substrate-binding pockets are shown as a cartoon model, with NADPH, 3-dehydro-ajmalicine (**16**) and mutated site shown as a stick model. The distance between the C4N atom of NADPH and the carbon atom of activated iminium is represented in dark dot line.

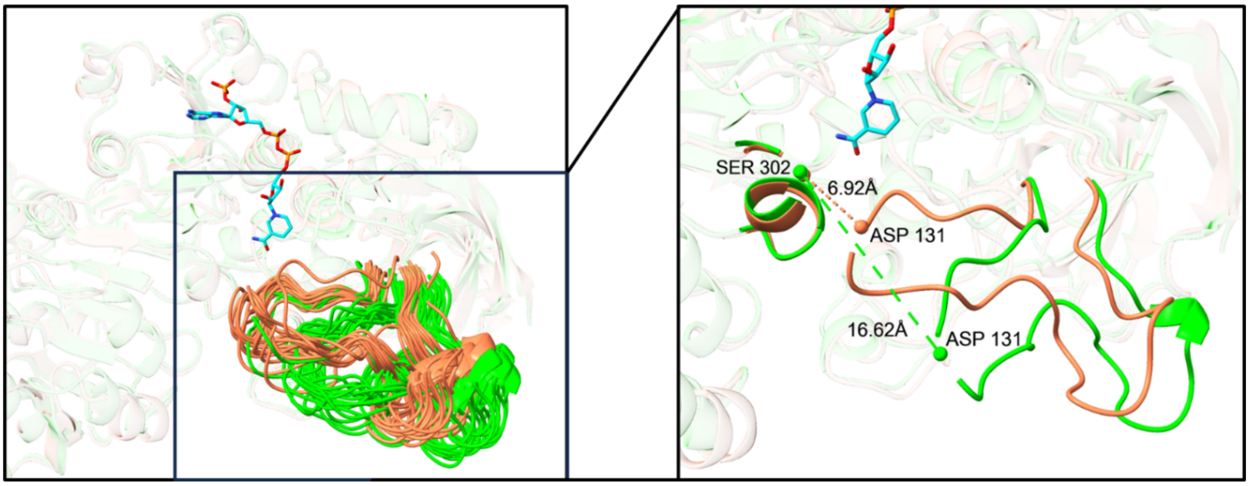

**Supplementary Fig. 24. Overlay of 20 snapshots (every 10 ns) obtained from 200 ns MD simulations for RvDTR-WT (in coral) and mutated variant RvDTR-T123G (in lime).** Insert containing two representative snapshots highlighted the flexibility and more-opened conformations acquired by the gate loop (residue 125-148) due to newly introduced mutation. Dot lines showed the distance change between the gate loop and the SER 302 (just as a reference residue for convenient measurement of distance) located at another chain.

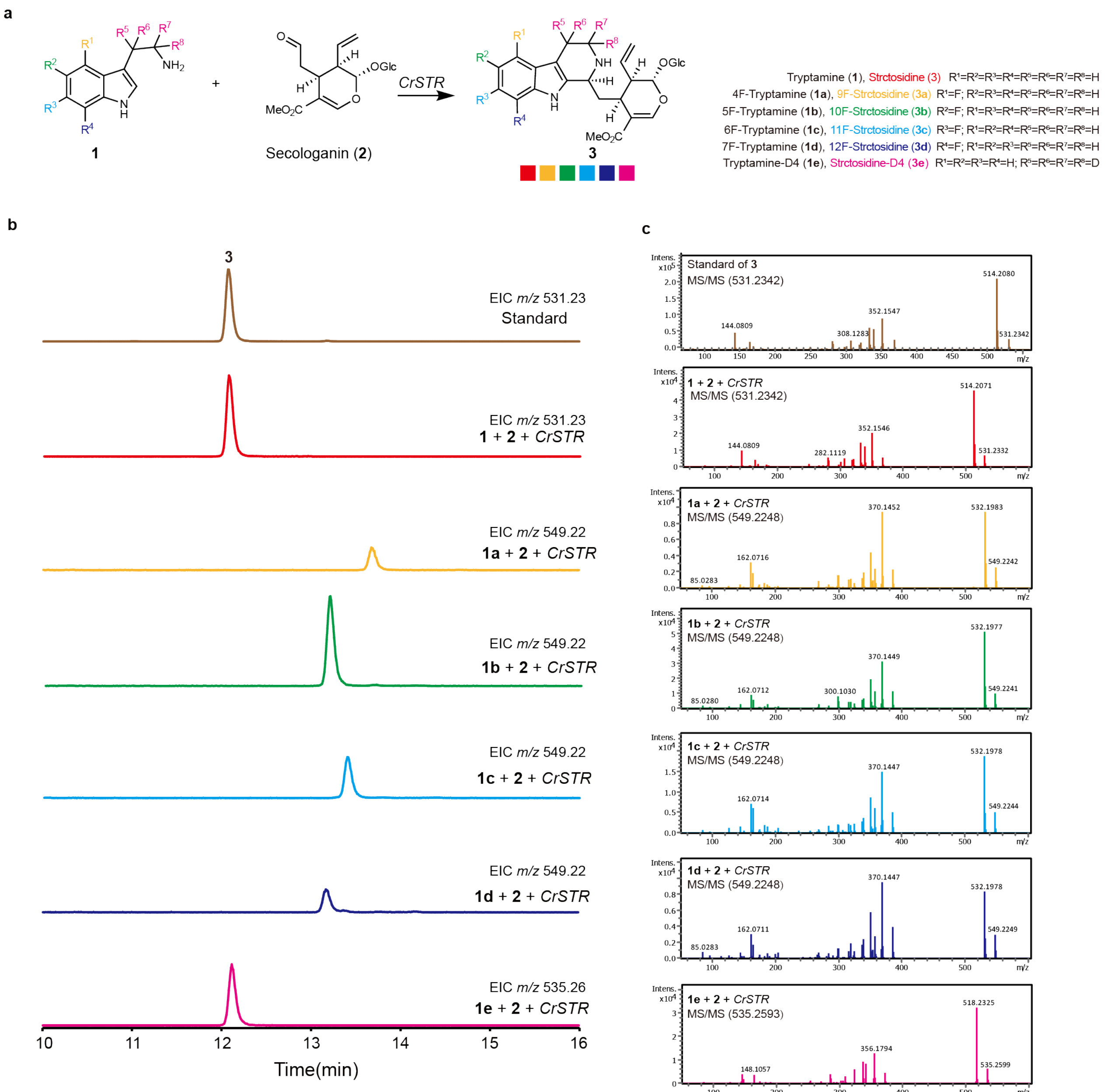

**Supplementary Fig. 25. LC-MS and LC-MS/MS analysis of fluorinated and deuterated srtictosidine (3-3e).** (**a**) Biosynthetic pathway depicting the conversion of secologanin (**2**) and fluorinated or deuterated tryptamines (**1-1e**) to fluorinated or deuterated srtictosidine (**3-3e**). Filled squares indicate the generation of product corresponding to *m/z*. (**b**) Extracted ion chromatograms of *m/z* corresponding to fluorinated or deuterated srtictosidine. Brown trace: authentic standard; Red, yellow, green, light blue, dark blue, pink traces: compound from transient expression of *CrSTR* with **2** and **1-1e** as substrate in *N. benthamiana*. (**c**) MS/MS spectra comparing authentic standard (**3**) with in planta-generated products.

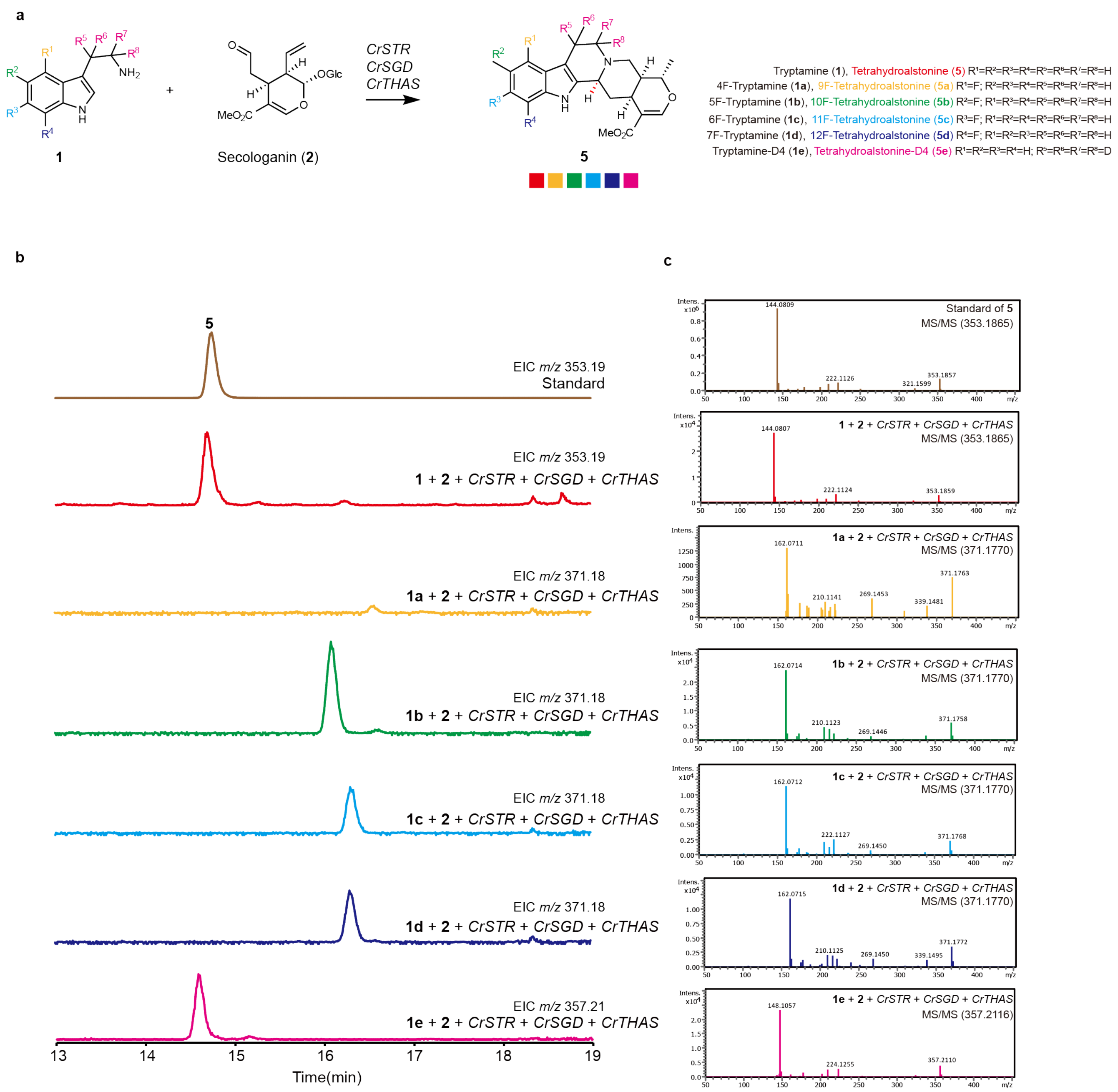

**Supplementary Fig. 26. LC-MS and LC-MS/MS analysis of fluorinated and deuterated tetrahydroalstonine (5-5e).** (**a**) Biosynthetic pathway depicting the stepwise conversion of secologanin (**2**) and fluorinated or deuterated tryptamines (**1-1e**) to fluorinated or deuterated tetrahydroalstonine (**5-5e**). Filled squares indicate the generation of product corresponding to *m/z*. (**b**) Extracted ion chromatograms of *m/z* corresponding to (**5-5e**). Brown trace: authentic standard; Red, yellow, green, light blue, dark blue, pink traces: compound from transient expression of *CrSTR, CrSGD, CrTHAS* with **2** and **1-1e** as substrate in *N. benthamiana*. (**c**) MS/MS spectra comparing authentic standard (**5**) with in planta-generated products.

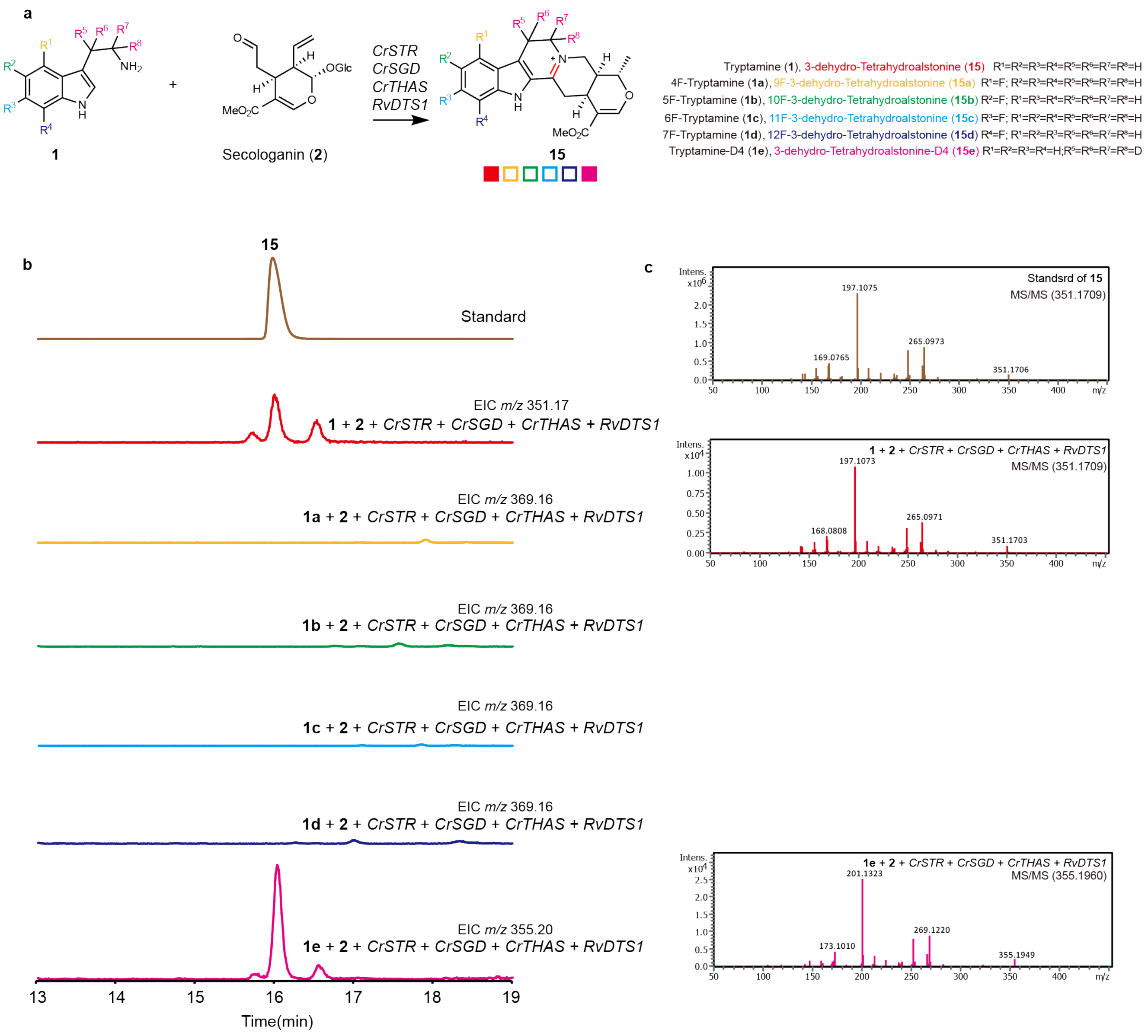

**Supplementary Fig. 27. LC-MS and LC-MS/MS analysis of fluorinated and deuterated 3-dehydro-tetrahydroalstonine (15-15e).** (**a**) Biosynthetic pathway depicting the stepwise conversion of secologanin (**2**) and fluorinated or deuterated tryptamines (**1-1e**) to fluorinated or deuterated 3-dehydro-tetrahydroalstonine (**15-15e**). Filled squares indicate the generation of product corresponding to *m/z* and empty squares indicate no generation of product corresponding to *m/z*. (**b**) Extracted ion chromatograms of *m/z* corresponding to **15-15e**. Brown trace: authentic standard; Red, yellow, green, light blue, dark blue, pink traces: compound from transient expression of *CrSTR, CrSGD, CrTHAS, RvDTS1* with **2** and **1-1e** as substrate in *N. benthamiana*. (**c**) MS/MS spectra comparing authentic standard (**15**) with in planta-generated products.

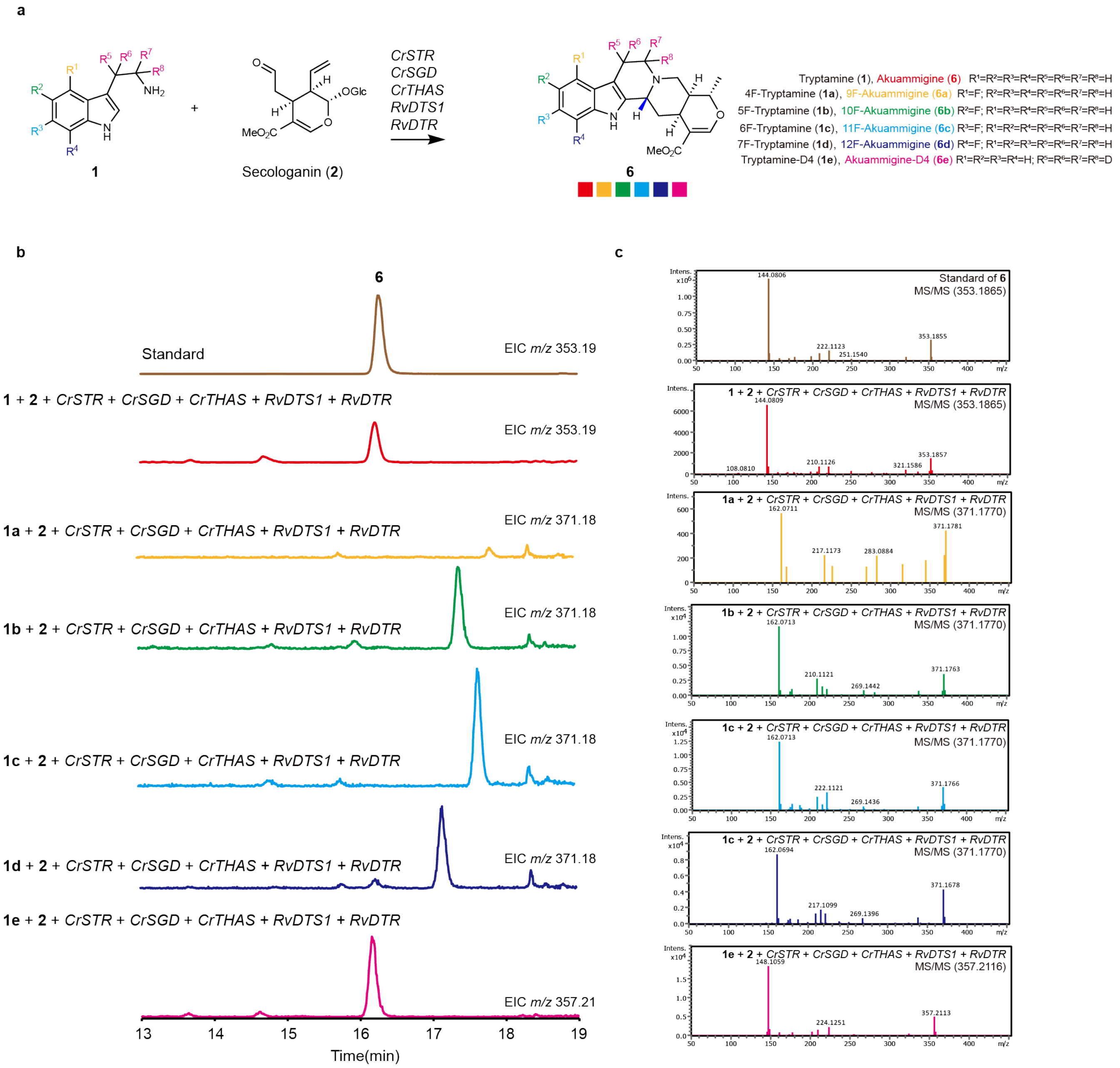

**Supplementary Fig. 28. LC-MS and LC-MS/MS analysis of fluorinated and deuterated akuammigine (6-6e).** (**a**) Biosynthetic pathway depicting the stepwise conversion of secologanin (**2**) and fluorinated or deuterated tryptamines (**1-1e**) to fluorinated or deuterated akuammigine (**6-6e**). Filled squares indicate the generation of product corresponding to *m/z*. (**b**) Extracted ion chromatograms of *m/z* corresponding to **6-6e**. Brown trace: authentic standard; Red, yellow, green, light blue, dark blue, pink traces: compound from transient expression of *CrSTR, CrSGD, CrTHAS, RvDTS1, RvDTR* with **2** and **1-1e** as substrate in *N. benthamiana*. (**c**) MS/MS spectra comparing authentic standard (**6**) with in planta-generated products.

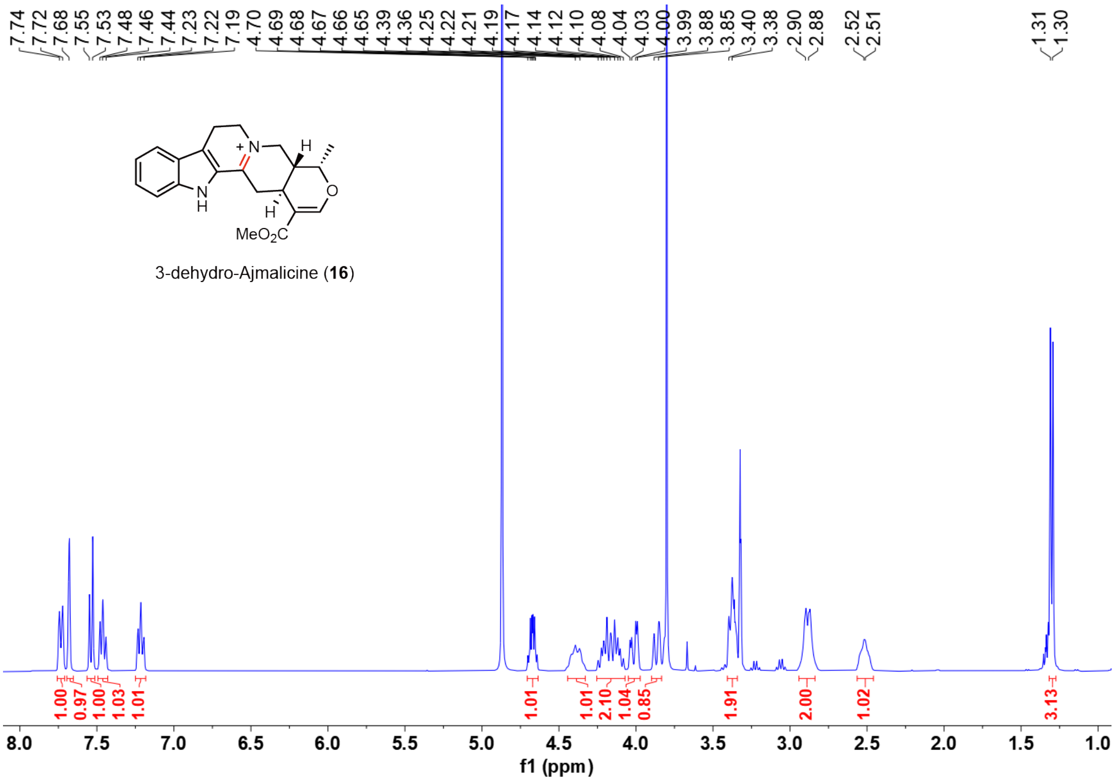

**Supplementary Fig. 29.** **$^{1}$H-NMR spectra of 3-dehydro-Ajmalicine (16) in $CD_3OD$.**

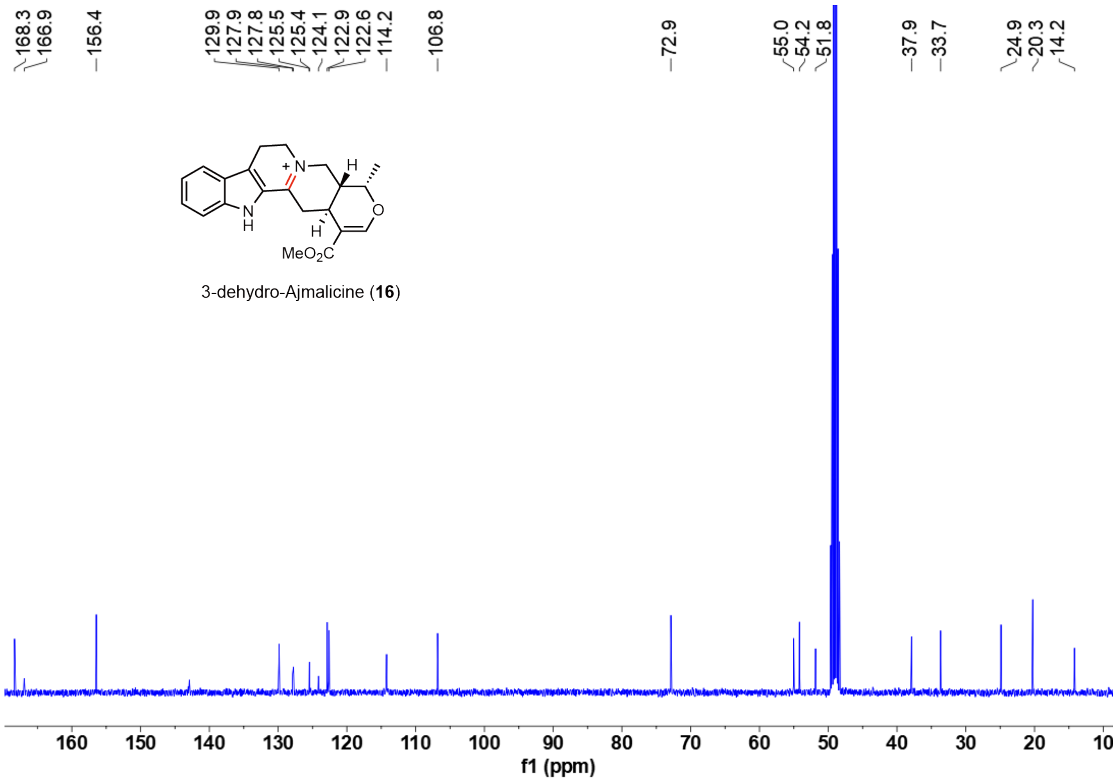

**Supplementary Fig. 30.** $^{13}$**C-NMR spectra of 3-dehydro-Ajmalicine (16) in CD₃OD.**

### 3. Supplementary Tables

**Supplementary Table 1. Chemical standards used in this study.**

| Chemical | CAS | Manufacturer |
|---|---|---|
| Corynantheine | 18904-54-6 | MedChemExpress |
| 20*S*-Corynantheidine | 23407-35-4 | GLPBIO |
| Mitraphylline | 509-80-8 | WeiKeQi |
| Isomitraphylline | 4963-01-3 | ChemFaces |
| Uncarine D | 4697-68-1 | zzstandard |
| Akuammigine | 642-17-1 | BioBioPha |
| Secologanin | 19351-63-4 | Sigma-Aldrich |
| Tryptamine | 61-54-1 | Aladdin |
| 4-Fluorotryptamine | 467452-26-2 | Macklin |
| 5-Fluorotryptamine hydrochloride | 2711-58-2 | KAIWEI CHEMICAL |
| 7-Fluorotryptamine hydrochloride | 159730-09-3 | TargetMol |
| Ethyl acetate | 141-78-6 | Thermo Fisher Scientific |
| Methanol | 67-56-1 | Thermo Fisher Scientific |
| Tryptamine-D4 Hydrochloride | 340257-60-5 | Anpel |
| 6-Fluorotryptamine hydrochloride | 55206-24-1 | Mreda |
| Tetrahydroalstonine | 6474-90-4 | BioBioPha |
| Uncarine F | 14019-66-0 | BioBioPha |
| Uncarine A | 6899-73-6 | BioBioPha |
| Hirsutine | 7729-23-9 | HerbSubstance |
| Hirsuteine | 35467-43-7 | BioBioPha |
| Corynoxeine | 630-94-4 | Must bio-technology |
| Isocorynoxeine | 51014-29-0 | WeiKeQi |
| Rhynchophylline | 76-66-4 | WeiKeQi |
| Isorhynchophylline | 6859-1-4 | WeiKeQi |
| Mayumbine | 25532-45-0 | HerbSubstance |
| Ajmalicine | 483-04-5 | HerbSubstance |
| Uncarine C | 5629-60-7 | HerbSubstance |
| Uncarine E | 5171-37-9 | HerbSubstance |
| Strictosidine | 20824-29-7 | Macklin |

**Supplementary Table 2. Nucleotide sequences for genes described and used in this study.**

| Gene Name | Nucleotide sequence |
|---|---|
| *RvDTS1* | ATGGAAACAAAAGTCCATATGGTTCTTTCATTTGTTATTTCC TTTCTCCTCCTCCTTTCATCTACCCATTCTAGTTCGATTCCT CAAGGCTTCATCGATTGCCTTTCCAATGAATTTTCATCAGA CATATATCTACTAGATGTTCTTTATGAGCCCAGCAATTCTTC TTTTGAACCTCTCCTGGAATCTACCATTCAAAATCTTAGATT CTTATCGTCTTCCACATTAAAGCCACTAGCTATAATCACCCC TATAGCTTTCTCCCATGTCCAAGCTACAGTAGTCTGCTGCA AACAAAGTGGACTGCAAATTAGAATACGAAGTGGCGGCC ATGATTATGAGGGGCTATCTTACCGATCCCGCCTTCCTTTTG TAATTCTTGACCTCCGAAATCTTACATCAGTAAGCGTGGAC GTCAAAGATAACAGCGCTTGGGTCGAGTCTGGAGCGACAC TTGGTGATTTGTATTATGGGATAGCTGAGAAAAGTCCTGTG CTTGCCTTCCCTGCTGGGCTCTGCCCAACTGTTGGTGTTGG CGGGCACTTGAGTGGTGGTGGAACTGGAAACTTGGTCAG AAAATACGGACTAGCTGCTGATAATGTCATCGATGCCCTCA TAGTTGTTGCCGACGGTCGAATTCTTGACCGGAAAACTAT GGGAACAGATCTGTTTTGGGCCATCAGGGGAGGTGGAGCT GCAAGTTTCGGGGTCGTGGTAGCATGGAAGATCAGGCTCG TACGTGTTCCTCCTACGGTCACTGTTTTCAAATTGAATAAG ACTTTAGAGCAAGGAGCTGTAAATCTTCTTCACAAATGGC AGTATATAGCGCACAAACTCCACGAAGACCTACTCATTACT GCAGAGATTTTCCGGAATGTTGGGAATAGAACACTTCAAG CGCAGTTCAGCTCGTTCTTCCTCGGTAGAGCTCACCAACT ACTGAAAACAATGGAGGAAATCTTCCCTGAACTTGGCCTG AGGAAAGAAGACTGCTCAGAAATGAGTTGGATCGAGTCC GTCGACTATTTTGCAGATTTTCCTAGCAGAGAAACTACAGA TAGCCTCAAGAAAAGCATACATTCCCCAGAGTTAACCAAA AACTACTTCAAGTCTAAGTCAGATTTCATTGTGGAACCACT TCCCCTTAGTTCATTACAAAAATTATGGAACCTGTGCCTCG AGGAGGAAAATCTTTCATTGCTAATGCATCCTTCTGGCGGG AAAATGGAGATGATAGCGGAATCAGAAACTCCATTTCCGT TCAGGCAAGGTTTGCTGTATGACATCCAATATGAAGTGGAC TGGTATAGCGAAAATCAATCATCAGAAAAACATATTGACTG GGTAAGAAAGATGTATGATTACATGACTCCTTACGCATCGA AACGACCAAGGGGTGCTTATCTTAATGCCAGAGATCTTGAT TTGGGTACAAATGACAATCCTTTCACAACGTATTCAGAGGC TCAAAGATGGGGGTTCAAATATTTCAAGAAAAACTTTAAG AGATTGGCTACTGTTAAGGGTGTCGTTGATCCAGAAAACT |

| | |
|---|---|
| | TCTTTTTCTTCGAGCAAAGCATTCCGCCTCTATGTAACAAG AATAACTTTAAGAGATTGGCTACTGCATAG |
| *UrDTS1* | ATGATCACAAAAATCTGTAAGATGCTATTGCTGATTTCAATT TTCTACTTAGTGATCCCATCATCACATTCAAGCTTGATTCTT CATAGTTTCATCAATTGCATTTCACGTAAGTTTCCATCAAAC ACTTCTATACTTGATGTCCTGTATCTCCCCAACAATTCTTCT TATCCACTTATCTTGCAGTCCACCATTCACAATCTTAGATTC TTGACACCTACCGCCCCCACCCCGCTTGCAATAATCACTCC TTTAGATTACTCTCATGTCCAAGCCACCGTTAAATGTAGCA AGCTGAACGGATTAAATATCAGAATCCGAAGTGGTGGCCA TGACTATGAAGGCATGTCATATACATCTGAAGTTCCGTTTG TCATACTTGACCTCAAAAACCTGAGGTCTATTAGCATTGAC ATTAAGAATAATAGTGCATGGGTTGAGGCTGGTGCAACAAT AGGCGAATTGTATTATTCCATTGCCGAGAGAAGTCCCATTC ATGGCTTTCCAGCAGGCCTTTGTCCAACTGTTGGTGTTGGT GGACACTTCAGTGGCGGCGGCGTAGGTAATTTGATCAGAA AGTATGGACTAGCTGCTGATAATGTCATCGATGCACTCATA GTTGATGTTAATGGTCGAATTCTAGATAGAAAATCAATGGG AGCTGATCTTTTTTGGGCAATAAGAGGAGGTGGAGGAGCA AGTTTCGGAGTTATAGTTGCCTGGAAAATCAAGCTTGTGC GTGTTCCACCTGTAGTTACTGTTTTCAATTTAACCAAGAGT TCAAGTCAAGAAGCCATAGGTCTTTTTAACAAGTGGCAAT ATGTTGCGCACAAGCTGAGTGAAGATTTGCTGGTTTACATT ACAATATCATCGATAGATGCCAAGGGAGGAGGAATTGCCG CAACATTTAATTCATTGTTCCTCGGTAAAACTGGTCAGCTC TTAAAAATGATGGAGGAGAGCTTCCCCGAAATTCACCTGA GAAAAGAAGATTGTATTGAGATGAGCTGGATTGATTCAGT CCTCCATTTTGCAGCATATCAAAATGGGGAAACTACAAAG GCCCTAAAGAGTAGAATTAACTCAATACCAAATAGTTACTT CAAGGGTAAGTCAGACGTGGTTCGCAAGCCTATACCATAT GAAGCATTAGAAGAGTTCTGGAAATGGTGTTCAGATATAA ATGCTCCTAACTTTTATGCAGAATTTCGTCCTTATGGTGGA AGAATGAATGAGATACCGGAATCAGAAACTCCATATCCAC ACAGGAAAAAAGTGCTCTATGAAATCCTCTACATGGTGTTT TGGACGAAGGATAAACATGGTGGATCTTCAAAAAATAACA TCAATTGGTTAAGAGGGTTATACAAGTTCATGACTCCTTAT GTGTCGAAAGGGCCAAGAGGCGCTGTTTGGAATTGTAGAG ATCTTGATTTAGGTGCAAATGGTGCTTCAGAAACTACTTAT TCTGAAGCCAAGGAATGGGGATCAAAGTATTTCAAGAACA ATTTTAAGAGGTTGGCAGTTATTAAAGGTGAAGTTGATCCA AATAATTTTTTCAATTATGAGCAAAGCATTCCACCTCTGGTT |

| | |
|---|---|
| | TTCCATGGATAA |
| *RvDTR* | ATGGATGCAGCATCTGCAAAAACAGCAACGCCAATTGAGG CCTACGGATGGGCAGCCAGAGACGCATCTGGAGTTCTCTC TCCATTCAACTTCCGAAGAAGGGCTACAGGGAAGCACGAT GTGCAGCTCAAAGTGTTGTATTGTGGTATGTGCGATTGGGA TCTACTTGTAGTCAAGAATTTGCTTGGCACTACTAAATATC CCATTGTACCTGGGCATGAGGTGGTGGGTGTGGTGACTGA GATCGGTAGCAAGGTGCAAAAGTTCAAGGTTGGGGACATA GCAGGTGTTAGCCACTACGTTCAGACATGTCGTAAATGTG AGAGATGCCAAGAAGGTCTTGACAGTTATTGTCCAAACTT GATAACAGCAGATGGAACTTCTTTTAGTGACGGAAACGAC CTATATTTCTACGATCCAAATGACACAGAGAGCAAGATGTA CGGTGCCTATTCCAACATCACGGTTGTCGATGAGTACTACG TAATCCGTTGGCCGGAAAACCTTCCTTTGGCTGCCGGCGT ACCTCTTTTATGTGCTGGTGTAGTTCCCTACAGCCCCATGA GATACTACGGATTTGATAAACCCGAAATTCATATTGGTGTC GTTGGACTTGGTGGGATGGGCAGATTAACCGTGAAATTTG CCAAGGCTTTCGGAGCAAAAGTTACAGTAATCAGTACATC CATTGACAAGAAGCAAGAAGCTATTGAGAAATACGGTGCA GATAGATTTTTACTCAGCAAAGAACCTGAGCAGCTGCAGG CCGCGGATGGGACGCTCGATGGCATCATTGACACAGTCCC TAGAGTTCACCCCCTTCGCGCATTGATCAAATTGTTGAAAT TCGACGGCACTCTTCTTTTGCTTGGAGCACCCCCGGAGCC ATATGAGTTGCCAGTCTCTCCACTGCTCGTAGGTAGGAAG AAGGTGGTTGGAAGTGGTGGTGCGAGTATAAAAGAAACA CAAGAGATGATGGATTTTGCAGCAAAGCACAATATAGTCG CAGATATAGAGATCATTCCAATGGGTTATGCAAACACTGCA ATCGAGCTTATAGAGAAGGGTGATTTCACAAAACGTTTCG TGATTGATATAGAGAATACATTGAAATCTACTTAG |
| *RvDTR13501* | ATGGACGCAGCATCTGCAAAAACAGCAACGCCAATTGAG GCCTACGGATGGGCAGCCAGAGACGCATCTGGAGTTCTCT CTCCATTCAACTTCCGAAGAAGGGCTACAGGAAAGCACG ATGTGCAGCTCAAAGTGTTGTATTGTGGGATGTGCGACTG GGATCTACTTGTAGTCAAGAATTTGCTTGGCACTACTAAAT ATCCCATTGTACCTGGGCATGAGGTGGTGGGTGTGGTGAC TGAGATCGGTAGCAAGGTGCAAAAGTTCAAGGTTGGGGA CATAGCAGGTGTTAGCCACTACGTTCAGACATGTCGTAAAT GTGAGAGATGCCAAGAAGGTCTTGACAGTTATTGTCCAAA CTTGATAACAGCGGATGGAACTTCTTTTAGTGATGGAAAC GATCTATATTTCTACGATCCAAATGACACAGAGAACAAGAT GTACGGTGCCTATTCCAACATCACGGTTGTCGATGAGTACT |

| | |
|---|---|
| | ACGTGATCCGTTGGCCGGAAAACTTTCCTTTGGCTGCCGG<br>CGTACCTCTTTTATGTGCTGGTGTAGTTCCCTACAGCCCCAT<br>GAGATACTACGGATTTGATAAACCCGGAATTCATATTGGTG<br>TCGTTGGACTTGGTGGGATGGGCAGATTAACCGTGAAATT<br>TGCTAAGGCTTTCGGAGCAAAAGTTACAGTAATCAGTACA<br>TCCATTGACAAGAAGCAAGAAGCTATTGAGAAATACGGTG<br>CAGATAGATTTTTACTCAGCAAAGAACCTGAGCAGCTGCA<br>GGCGGCGGTTGGGACGCTCGATGGCATCATTGACACAGTC<br>CCTAGAGTTCACCCCCTTCGCGAATTGATCAAATTGTTGAA<br>ATTCGACGGCACTCTTCTTTTGCTTGGAGCACCCCCGGAG<br>CCATATGAGTTGCCAGTCTCTCCACTGCTCGTAGGTAGGAA<br>GAAGGTGGTTGGAAGTGGTGGTGCGAGTATAAAAGAAAC<br>ACAAGAGATGATGGATTTTGCAGCAAAGCACAATATAGTC<br>GCAGATATAGAGATCATTCCAATGGGTTATGCAAACACTGC<br>AATCGAGCGGATAGAGAAGGGTGATTTCACAAAACGATTC<br>GTGATTGATGTAGAGAATACATTGAAATCTGCTTAG |
| *MsCYP72056* | ATGGCAAGCTTCTCAGCATTTCAGGAAGAAGCTAAGAGAT<br>GGCTCTCAAAGCATTTAGTGGTAGTATCTTTGCTTCTTCCTT<br>CGTTCATGATCTTCATTCTATTTCTTTCCAAATGGTGGTTCC<br>TCACTACCAGTAAGAAAAAGAACCCTCCACCCTCACCTCC<br>TAAGCTGCCAATAATTGGAAACCTTCATCAAATTGGTTCTG<br>TCGCCCACCGCTCTTTCAAATCCTTGGCCGAAAAGCATGG<br>TCCGCTAATGCTGCTTCATGTAGGCTGGCAAAAACTGCTAG<br>TTGTCTCTTCCGCCGAAACAGCTCGTGAGGTCCTCAAAAC<br>ACATGATGTTGCCTTTTGCGGTAGACCTGATTCAGAGGTTA<br>CCCGAAGGATTTTCTATAACTTGAAGAACATATCTCTCTCC<br>CCCTATGGTGATTATTGGAAGATAGTGAGAAGCATCGCTGT<br>GAATCAAATTCTAAATCAGAAGAGGGTTCAGTCGTTCCGA<br>AGTGTAAGAGAAGAAGAGATTTCATTACTGGTGGAAAGAA<br>TCAAAGAATCTTGCGCTTCTTCCTCCGTAATATGCATGAAC<br>ACATTGTTGACAACGCTTGTAAATGACATAGTGTCAAGGAT<br>AACCATTGGGAAGAGGTCCTCAGGAATAGGTGGCAGCAG<br>ATTTGCAGAGTTTTTGTTTGCATCCGTCGAGTTATTAGGTTC<br>TTTTCTTGCAGGGGACTTCTTCCCGGGGCTTGGATGGCTCG<br>GTCACATAACTGGATTCGAAGCAAAAATTAAGAAAGTTTC<br>CAAAGATTTGGATCAATGTTTGGAGAACTTAATTGAAGAG<br>GAAATAAACAGGAACAAAAGAGGAGATGACCAAGGTAAA<br>GACAATCAGAATTTCCTCCAGGTTTTGCTTGAAATCCAGA<br>GAACCGACTCATCTGGCCATGCTTTGGATCGAGAATCCATT<br>AAGGCTGTCATAATGGACATGGTTGCCGGTGCATTTGACTC<br>GTATACACTTTTGGAGTGGGCGACGTCATTGCTAGTAAAAC |

| | |
|---|---|
| | ATCCAGATGTCATGAAAAAATTGCAAAATGAGGTGAAAGA<br>AGTTGCTGGATCCAAATCATTCATATCGGAGGATGATTTAA<br>GTAAACTGCAATACTTGAAAGCAGTAATAAAAGAAACTTT<br>CCGACTCTGTCCCGGTGCATTTATTGTAAGGGTATCAACCA<br>AGGATGTCAAAATAATGGGCTATGATATTGCAGCAGGCACT<br>CAAGTCCTCGTCAATGCATTGGCAATTGGAAGGGATCCAA<br>CGTTGTGGGAAAAACCTGAGGAGTTTCGGCCAGAAAGGT<br>TCTTGAATAGTTCAATAGATCTGAAAGGACATCACTTTGAA<br>CTGCTTCCATTTGGTTCAGGAAGAAGGTCTTGCCCCGGTT<br>CTACATTTGCCTTGGTCATAGATGAGCTCGTACTGGCAAAT<br>TTGGTGTGCAACTTTAATTACGCATTGCCTGGTGGAGCAAA<br>AGCCGAGGACTTGGACATGACTGAAGCTCCTGGTGCCATG<br>CCTCGTAGGAGAACCCCTCTGCTGCTCGTTCCATCTCTTTG<br>TTAA |
| Codon Optimized *RvDTR* | ATGGACGCTGCTTCTGCTAAGACCGCTACTCCAATCGAAG<br>CTTACGGTTGGGCTGCACGTGATGCTTCTGGTGTCTTGTCT<br>CCATTCAACTTCAGAAGAAGAGCTACCGGTAAGCACGATG<br>TTCAATTGAAGGTTTTGTACTGTGGTATGTGTGACTGGGAC<br>TTGTTGGTTGTCAAGAACTTGTTGGGTACCACTAAATACCC<br>AATTGTTCCTGGCCACGAAGTTGTTGGTGTTGTCACCGAA<br>ATCGGTTCTAAGGTCCAAAAGTTCAAGGTTGGCGATATTGC<br>TGGTGTTTCTCACTACGTCCAAACTTGTAGAAAGTGTGAA<br>AGATGCCAAGAAGGTTTGGACTCTTACTGTCCAAACTTGA<br>TTACCGCCGATGGTACCTCTTTCTCAGACGGTAACGACTTA<br>TATTTCTACGACCCAAACGATACTGAATCCAAGATGTACGG<br>TGCCTACTCTAATATCACTGTCGTTGACGAATACTACGTTAT<br>CAGATGGCCAGAAAACTTACCATTGGCTGCCGGTGTTCCA<br>TTATTGTGTGCTGGTGTTGTTCCTTACAGCCCAATGAGATA<br>CTATGGTTTCGACAAGCCAGAAATTCACATTGGTGTCGTCG<br>GTTTAGGTGGTATGGGTAGATTGACTGTTAAGTTTGCCAAG<br>GCTTTCGGTGCCAAGGTCACAGTTATCTCCACTTCTATTGA<br>CAAGAAGCAAGAAGCTATTGAGAAGTACGGTGCTGACAG<br>ATTCTTGTTGTCCAAGGAACCAGAACAATTGCAAGCTGCT<br>GACGGTACTTTGGATGGTATCATTGACACCGTCCCAAGAGT<br>TCATCCTTTGAGAGCTTTGATCAAGTTGTTGAAATTCGATG<br>GTACTCTACTTCTATTGGGTGCTCCACCAGAACCATACGAA<br>TTGCCAGTTTCCCCATTGTTGGTCGGACGTAAGAAAGTGG<br>TCGGTTCCGGTGGTGCCTCCATCAAGGAAACTCAAGAAAT<br>GATGGATTTCGCTGCCAAACACAACATCGTTGCTGATATTG<br>AAATCATCCCAATGGGTTACGCTAACACCGCTATTGAATTA<br>ATCGAAAAGGGTGACTTCACCAAGAGATTTGTCATTGATAT |

| | |
|---|---|
| | CGAAAACACTTTGAAGTCTACCTGA |
| Codon Optimized *AtATR2* | ATGTCCTCTTCTTCTTCTTCGTCAACCTCCATGATCGATCTC<br>ATGGCAGCAATCATCAAAGGAGAGCCTGTAATTGTCTCCG<br>ACCCAGCTAATGCCTCCGCTTACGAGTCCGTAGCTGCTGA<br>ATTATCCTCTATGCTTATAGAGAATCGTCAATTCGCCATGAT<br>TGTTACCACTTCCATTGCTGTTCTTATTGGTTGCATCGTTAT<br>GCTCGTTTGGAGGAGATCCGGTTCTGGGAATTCAAAACGT<br>GTCGAGCCTCTTAAGCCTTTGGTTATTAAGCCTCGTGAGGA<br>AGAGATTGATGATGGGCGTAAGAAAGTTACCATCTTTTTCG<br>GTACACAAACTGGTACTGCTGAAGGTTTTGCAAAGGCTTT<br>AGGAGAAGAAGCTAAAGCAAGATATGAAAAGACCAGATT<br>CAAAATCGTTGATTTGGATGATTACGCGGCTGATGATGATG<br>AGTATGAGGAGAAATTGAAGAAAGAGGATGTGGCTTTCTT<br>CTTCTTAGCCACATATGGAGATGGTGAGCCTACCGACAATG<br>CAGCGAGATTCTACAAATGGTTCACCGAGGGGAATGACAG<br>AGGAGAATGGCTTAAGAACTTGAAGTATGGAGTGTTTGGA<br>TTAGGAAACAGACAATATGAGCATTTTAATAAGGTTGCCAA<br>AGTTGTAGATGACATTCTTGTCGAACAAGGTGCACAGCGT<br>CTTGTACAAGTTGGTCTTGGAGATGATGACCAGTGTATTGA<br>AGATGACTTTACCGCTTGGCGAGAAGCATTGTGGCCCGAG<br>CTTGATACAATACTGAGGGAAGAAGGGGATACAGCTGTTG<br>CCACACCATACACTGCAGCTGTGTTAGAATACAGAGTTTCT<br>ATTCACGACTCTGAAGATGCCAAATTCAATGATATAAACAT<br>GGCAAATGGGAATGGTTACACTGTGTTTGATGCTCAACATC<br>CTTACAAAGCAAATGTCGCTGTTAAAAGGGAGCTTCATAC<br>TCCCGAGTCTGATCGTTCTTGTATCCATTTGGAATTTGACAT<br>TGCTGGAAGTGGACTTACGTATGAAACTGGAGATCATGTT<br>GGTGTACTTTGTGATAACTTAAGTGAAACTGTAGATGAAGC<br>TCTTAGATTGCTGGATATGTCACCTGATACTTATTTCTCACT<br>TCACGCTGAAAAAGAAGACGGCACACCAATCAGCAGCTC<br>ACTGCCTCCTCCCTTCCCACCTTGCAACTTGAGAACAGCG<br>CTTACACGATATGCATGTCTTTTGAGTTCTCCAAAGAAGTC<br>TGCTTTAGTTGCGTTGGCTGCTCATGCATCTGATCCTACCG<br>AAGCAGAACGATTAAAACACCTTGCTTCACCTGCTGGAAA<br>GGATGAATATTCAAAGTGGGTAGTAGAGAGTCAAAGAAGT<br>CTACTTGAGGTGATGGCCGAGTTTCCTTCAGCCAAGCCAC<br>CACTTGGTGTCTTCTTCGCTGGAGTTGCTCCAAGGTTGCA<br>GCCTAGGTTCTATTCGATATCATCATCGCCCAAGATTGCTGA<br>AACTAGAATTCACGTCACATGTGCACTGGTTTATGAGAAA<br>ATGCCAACTGGCAGGATTCATAAGGGAGTGTGTTCCACTT<br>GGATGAAGAATGCTGTGCCTTACGAGAAGAGTGAAAACTG |

| | |
|---|---|
| | TTCCTCGGCGCCGATATTTGTTAGGCAATCCAACTTCAAGC<br>TTCCTTCTGATTCTAAGGTACCGATCATCATGATCGGTCCA<br>GGGACTGGATTAGCTCCATTCAGAGGATTCCTTCAGGAAA<br>GACTAGCGTTGGTAGAATCTGGTGTTGAACTTGGGCCATC<br>AGTTTTGTTCTTTGGATGCAGAAACCGTAGAATGGATTTCA<br>TCTACGAGGAAGAGCTCCAGCGATTTGTTGAGAGTGGTGC<br>TCTCGCAGAGCTAAGTGTCGCCTTCTCTCGTGAAGGACCC<br>ACCAAAGAATACGTACAGCACAAGATGATGGACAAGGCTT<br>CTGATATCTGGAATATGATCTCTCAAGGAGCTTATTTATATG<br>TTTGTGGTGACGCCAAAGGCATGGCAAGAGATGTTCACAG<br>ATCTCTCCACACAATAGCTCAAGAACAGGGGTCAATGGAT<br>TCAACTAAAGCAGAGGGCTTCGTGAAGAATCTGCAAACG<br>AGTGGAAGATATCTTAGAGATGTATGGGGATCCTAA |
| Codon Optimized *MsCYP72056* | ATGGCTTCTTTCTCTGCTTTCCAAGAAGAAGCTAAGAGATG<br>GTTATCTAAGCATTTGGTTGTTGTTTCCCTTCTACTACCATC<br>CTTCATGATTTTCATTTTGTTCTTGTCAAAATGGTGGTTCTT<br>GACTACCTCCAAGAAAAAGAACCCACCTCCATCTCCACCA<br>AAGTTGCCAATCATTGGTAACTTACACCAAATTGGTTCTGT<br>TGCTCACAGATCCTTCAAGTCTCTAGCAGAAAAGCACGGT<br>CCATTGATGTTGTTGCACGTTGGTTGGCAAAAGCTGCTGG<br>TCGTCAGTTCTGCTGAAACTGCTAGGGAAGTCCTGAAGAC<br>CCACGATGTCGCGTTCTGTGGTAGACCAGATTCTGAAGTTA<br>CCAGACGTATTTTCTACAACTTGAAGAACATCTCTTTGTCC<br>CCATACGGTGACTACTGGAAGATTGTCAGATCCATTGCCGT<br>TAACCAAATCTTGAACCAAAAGAGAGTCCAATCTTTCCGT<br>TCTGTCAGAGAAGAAGAAATCTCTTTGTTAGTCGAACGTA<br>TAAAGGAATCTTGTGCTTCTTCGTCCGTGATCTGTATGAAC<br>ACCCTCTTGACCACTTTGGTCAACGACATTGTCTCCAGAAT<br>CACTATTGGTAAGAGATCCTCTGGTATCGGTGGTTCCAGAT<br>TCGCCGAATTTTTGTTTGCTTCCGTTGAATTATTAGGTTCCT<br>TCTTGGCCGGTGACTTCTTCCCTGGTTTAGGATGGTTGGGT<br>CACATTACTGGTTTCGAAGCTAAGATCAAAAAGGTTTCTA<br>AGGACCTCGATCAATGTTTGGAAAACTTGATTGAAGAAGA<br>AATCAACAGAAACAAGAGAGGTGATGACCAAGGTAAAGA<br>CAACCAAAATTTTTTGCAAGTTTTGTTGGAAATCCAAAGA<br>ACCGATTCCTCTGGTCACGCTTTGGACAGAGAATCCATCA<br>AGGCTGTCATCATGGACATGGTTGCCGGTGCTTTCGACTCT<br>TACACCTTGTTGGAATGGGCTACTTCTTTGTTGGTTAAACA<br>TCCAGACGTTATGAAGAAGCTGCAAAACGAGGTCAAGGA<br>AGTTGCCGGTTCCAAGTCTTTCATCTCCGAAGACGATTTGT<br>CTAAATTGCAATACTTGAAGGCCGTTATCAAGGAAACCTTC |

| | |
|---|---|
| | AGATTGTGTCCAGGTGCCTTTATCGTTCGTGTTTCTACCAA GGATGTTAAGATCATGGGTTACGACATTGCTGCTGGTACTC AAGTCTTGGTTAACGCTTTGGCTATTGGTAGAGACCCAACT TTGTGGGAAAAGCCAGAAGAATTTAGACCAGAAAGATTCT TGAACTCTTCCATTGATTTGAAGGGTCATCACTTCGAATTG TTGCCATTCGGTTCCGGTCGTCGTTCCTGCCCCGGCTCAAC TTTCGCTTTGGTCATCGATGAATTGGTTCTTGCTAACTTGGT CTGTAACTTCAACTATGCCTTGCCAGGTGGTGCTAAGGCTG AAGATTTGGACATGACTGAAGCTCCAGGTGCTATGCCAAG AAGAAGAACTCCATTGTTATTGGTTCCATCCTTATGTTGA |
| *CrSTR* | ATGGCAAACTTTTCTGAATCTAAATCCATGATGGCAGTTTT CTTCATGTTTTTCCTTCTTCTTCTTTCTTCTTCTTCTTCTTCT TCTTCTTCTTCACCAATTTTGAAAAAGATTTTTATTGAAAG CCCTTCCTATGCTCCGAATGCCTTCACCTTCGATTCAACTG ATAAAGGGTTCTACACTTCCGTCCAAGATGGCCGAGTTATC AAATATGAAGGGCCAAATTCAGGCTTCACTGACTTCGCCT ACGCATCTCCCTTCTGGAACAAAGCTTTTTGTGAGAACAG CACCGATCCAGAGAAAAGACCATTGTGTGGGAGGACATAT GATATTTCCTATGACTATAAGAACAGCCAAATGTACATTGTT GATGGCCATTACCATCTTTGTGTGGTTGGAAAAGAAGGTG GGTATGCCACACAACTAGCCACAAGTGTGCAAGGAGTGCC ATTCAAATGGCTCTATGCAGTAACTGTTGATCAGAGAACAG GGATTGTTTATTTCACTGATGTTAGCTCCATACATGATGACA GTCCCGAAGGTGTGGAAGAAATCATGAATACAAGTGATAG AACAGGGAGATTAATGAAGTATGATCCTTCAACAAAAGAA ACCACCTTATTATTGAAAGAGCTACATGTTCCCGGCGGTGC AGAAATCAGCGCAGATGGTTCCTTTGTTGTAGTAGCAGAA TTTTTAAGCAATCGGATAGTGAAGTATTGGCTAGAAGGGCC AAAGAAAGGCAGTGCAGAGTTCTTAGTTACAATCCCAAAT CCAGGAAATATAAAGAGGAATTCTGATGGCCATTTTTGGGT GTCTTCAAGTGAAGAATTAGATGGAGGTCAACATGGAAGA GTTGTTTCAAGAGGAATTAAGTTTGATGGATTTGGGAATAT TCTTCAAGTTATACCACTTCCACCACCATATGAAGGTGAAC ATTTTGAACAGATTCAAGAGCACGATGGTTTGTTATACATT GGAAGTCTCTTCCATAGCTCTGTGGGTATATTAGTGTATGAT GATCATGATAACAAGGGAAATTCTTATGTTTCTAGCTAG |
| *CrSGD* | ATGGGATCTAAAGATGATCAGTCCCTTGTTGTTGCCATTTC TCCAGCTGCTGAACCAAATGGAAATCATTCTGTCCCCATCC CATTCGCCTACCCCAGTATCCCCATTCAACCTAGAAAGCAC AACAAGCCCATCGTTCATCGTCGAGATTTCCCCTCAGATTT CATCTTGGGTGCCGGAGGATCTGCTTATCAGTGTGAGGGT |

| | |
|---|---|
| | GCATATAATGAAGGCAACCGCGGTCCCAGTATATGGGATAC TTTCACAAACCGATATCCAGCCAAAATAGCTGATGGATCTA ATGGCAATCAAGCCATCAATTCTTACAATTTGTACAAGGAA GATATCAAGATTATGAAGCAAACAGGCTTGGAATCATATAG GTTTTCAATTTCATGGTCAAGAGTATTGCCAGGTGGGAATC TATCCGGTGGAGTGAATAAAGATGGTGTCAAGTTCTATCAT GACTTTATAGATGAGCTTCTAGCCAATGGCATCAAACCCTT TGCAACTCTCTTCCACTGGGATCTTCCCCAAGCTCTTGAAG ACGAGTATGGAGGCTTCTTGAGTGATCGAATTGTGGAAGA TTTTACGGAGTATGCAGAATTTTGCTTTTGGGAATTCGGTG ACAAAGTAAAATTTTGGACGACTTTCAATGAACCACATAC TTATGTTGCAAGTGGATATGCCACTGGTGAATTTGCACCAG GAAGAGGTGGTGCAGATGGCAAGGGGGAACCTGGCAAAG AACCCTATATAGCGACACATAATTTACTTCTTTCTCACAAA GCTGCTGTGGAAGTATATAGGAAAAATTTTCAGAAATGTCA AGGAGGTGAAATTGGAATTGTACTTAATTCAATGTGGATGG AGCCTCTCAATGAAACCAAAGAAGATATTGATGCTCGGGA AAGGGGTCTTGATTTCATGCTCGGATGGTTCATAGAGCCAT TAACAACGGGTGAATACCCAAAATCCATGAGAGCTCTTGT AGGAAGCCGTCTTCCAGAATTTTCAACAGAAGTTTCCGAA AAATTAACAGGATGCTATGATTTTATCGGAATGAATTATTAT ACAACTACTTATGTTTCTAATGCAGACAAAATTCCCGATAC TCCGGGTTACGAAACAGATGCTCGAATTAATAAGAATATTT TTGTCAAAAAAGTTGATGGGAAGGAAGTGCGCATTGGTGA ACCGTGCTATGGGGGATGGCAGCATGTTGTTCCATCTGGAC TCTACAATCTCTTGGTTTACACTAAGGAGAAATACCATGTT CCAGTGATTTATGTCTCAGAATGTGGTGTGGTTGAGGAAA ATAGAACCAACATATTACTTACAGAAGGTAAAACCAACATA TTACTTACAGAAGCTCGTCACGATAAACTCAGGGTTGATTT TCTACAAAGTCATCTCGCTAGCGTGCGAGATGCTATTGATG ATGGTGTGAATGTAAAAGGATTCTTTGTTTGGTCATTCTTC GACAACTTCGAATGGAATTTGGGATATATATGCCGTTATGG AATTATCCATGTTGATTATAAAACTTTTCAAAGATATCCAAA GGATTCTGCCATATGGTACAAGAATTTCATTAGTGAAGGAT TTGTTACGAATACAGCTAAAAAGAGATTCCGAGAAGAAGA TAAACTAGTTGAGTTAGTCAAGAAGCAAAAATACTAA |
| *CpDCS* | ATGGCCGGAAAATCTCAAGAAGATGGGCAGACGGTAAAG GCTCTAGGATGGGCCGCTAGGGAAGTTTCTGGGGCGATCT CTCCTTTCGATTTCTCAAGAAGGGCCCCAGGAGAGCGCGA TGTGCAGGTTAAAATACTATATTGTGGAATCTGTAGTTTTG ACACAGAAATGATCAATAACAAGTTTGGCTTTACCAGATAT |

| | |
|---|---|
| | CCCTTTGTACTCGGGCATGAGATTGTGGGAGTGGTATCTGAAGTTGGTAGAAAGGTGCAAAAATTCAAGATTGGGGATAAAGTTGGTGTAGGAACCATGATTGGATCTTGTCGCACTTGTTATAGCTGCACTCACAATCTCGAAAATTACTGCCCAAAAGTTACATTAACAGAAGCAACTTCTGGTGGTTGTTCTAATCTTGTGATAGCAGATGAAGACTTTGTGTTCCATTGGCCGGTGAATTTGCCTCTTGATCTTGGAGCTCCTCTCCTTTGTGCTGGGATTACTGTTTATAGCCCTTTGAAAAATTTTGAACTTGATAAGCCTGGATTGCGTATTGGTGTGGTTGGTCTTGGTGGTATTGGCCATATAGCTGTAAAATTTGCCAAGGCTTTTGGGGCTAAGGTGACAGTGATTAGTTCATCAGAAAGTAAAAAGGTTGAAGCCATTGAAAAATATGGTGCAGATTCCTTTTTGGTTAGCAGTGATCCAGGGCAGATGCTGGCAGCTGCCGGAACCTTGGATGGTGTCATTGATACCGTCCCAGCACCTCACTCTATTTTGCCATTCCTTGATTTACTCTTGCCTCGTGGAAAGCTAATTATATTAGGTGCACCAATGGAGCCATTTGTACTGCCAATCTATCCCCTGCTTCAAGGTGGGAGAGTAGTTGCTGGGAGTGCCACTGGAGGATTGAAACAAATCCAAGAAATGCTTCATTTTGCAGCAGAGCACAACATAGTAGCAGATGGCGAGGTTATCCCAATCGACGACATTAACACTGCGATAAAGCGCATTGAGAAAGGCGATGTCAAATATCGATTTGTGGTTGACATTGGCAATACCTTAAAATCTGCTTGA |
| *MsEnolMT* | ATGCAACCACAGAGAGGGAGAAAGAGAGAGAGAGAGAGAGATAGAGAAGAGATGGAATCCGTGCAGAGCAACAGTAGTTCTTCTGATCAATTCGCAATGAAAGGTGGAGATGACGACTTCAGTTACACAAAGAATTCCACCTGGCAGAGAGATGCAATTCAAGCAACCAAATTTTTCATTCAAGAATCTATTGCTGAGAAGCTTGACGTCAATAAATTTTGTGGAAAGGCATTTTGCGTTGCTGATTTGGGATGCTCAGTTGGACCTAACACTTTGATAGCAATGCAGAACATTGTTGAAGCTGTGGAGCTTAAATTCAAAAATAGAAAAGGATTCCATTCTCCCACTATCCCTGAATTTCAAGTCTTCTTTAACGATCATACGGTGAATGATTTCAATACCCTCTTTAGATCTCTCCCAACTGGTCACGACAAGCGCTATTACGGCGTTGGGGTTCCGGGTTCCTTTTACGGTCGATTATTTCCTTGTGACTCTATTCACATAATGCACACTTCATTTTCTACACCGTTTCTTTCTCAAGTACCAAAAGAGGTGATTGACAAAAATTCAGCTGCGTGGAATAAAGGAAGGATTCATCACAATTATGCTAAAGCAGATGTTTTGAAGGCTTATGAAGCACAACATGCTGAGGATATCGACTGCTTTTTGACGGCTAGAGCTAAAGAACTGGTCCATGGAGGATTATTGATGGATGTGACTTCATTCCGCCCAGATGGGGTCCCTCATACCCATGTCTTGACTAACATAG |

| | |
|---|---|
| | GGATGGAGGTATTGGGTTATTGCCTCATGGACTTGGCTGGA CTTATCGATGAAGAAAACGTGGATTCCTACAACGTTCCAGT TTATCTTCAATCTCCTGAAGAGTTGAAACAAGCTGTTCAAC GGAACAAATACTTCAGTATAGAAAAAATGGAGAGCGTGCC TATGATGATAGATTCAGATGTTTCTGCCAAAGCTCAACAAT ATTCATTGGGAATGAGGGCCGTAATGGGGGACGTGATTAG AGAGCAATTTGGAGCGGAGATAGTGGATAAACTCTTTGAT TTGTTCAAGAAGAAACTTGAAGAGCATCCTAACTTTGCAA AAGGAGTTGTCCTTGACATGTTTGTTCTCCTTAAACGCAAT GCAGAGGATTGA |
| *CrTHAS* | ATGGCAATGGCTTCAAAGTCACCTTCTGAAGAAGTATATCC AGTGAAGGCATTTGGTTTGGCTGCTAAGGATTCTTCTGGGC TTTTCTCTCCATTCAACTTCTCAAGAAGGGCCACAGGGGA ACACGATGTGCAGCTCAAAGTATTATACTGTGGGACTTGCC AATATGACAGGGAAATGAGCAAAAACAAATTTGGATTTAC AAGCTATCCTTATGTTTTAGGGCATGAAATTGTGGGTGAGG TAACTGAAGTTGGCAGCAAGGTGCAGAAATTCAAAGTCG GGGACAAAGTGGGCGTAGCAAGCATAATTGAAACTTGTGG CAAATGTGAAATGTGTACAAATGAAGTTGAAAATTACTGT CCAGAAGCAGGATCAATAGACAGCAATTACGGGGCATGTT CAAATATAGCAGTGATAAACGAGAATTTTGTCATCCGTTGG CCTGAAAATCTTCCTTTGGATTCTGGTGTTCCTCTTCTATGT GCAGGAATCACGGCTTATAGTCCCATGAAACGTTATGGACT TGATAAACCTGGAAAACGTATCGGCATAGCCGGTCTAGGA GGACTTGGACATGTAGCTCTTAGATTTGCCAAAGCTTTTGG GGCTAAGGTGACAGTGATTAGTTCTTCACTTAAGAAAAAA CGTGAAGCCTTTGAGAAATTCGGAGCAGATTCTTTCTTGG TCAGCAGTAATCCAGAAGAAATGCAGGGTGCAGCAGGAA CATTGGATGGGATCATAGACACTATACCAGGGAATCACTCT CTTGAGCCACTCCTTGCTTTATTGAAGCCTCTTGGGAAGCT TATCATTTTAGGTGCACCAGAAATGCCCTTTGAGGTTCCCG CTCCTTCCCTGCTTATGGGTGGAAAAGTAATGGCTGCCAGT ACTGCTGGGAGTATGAAGGAAATACAAGAGATGATTGAAT TTGCAGCAGAACACAACATAGTAGCAGATGTGGAGGTTAT CTCTATTGACTATGTGAACACTGCAATGGAGCGCCTTGATA ACTCTGATGTGAGATATCGTTTCGTGATTGATATAGGGAAC ACTCTGAAATCAAATTAA |

**Supplementary Table 3. Primers used for genes cloning in this study.**

Primers for yeast vector construction (pESC vector). Cloning overhangs are underlined.

| Primer | Plasmid | Primer direction | Sequence |
|---|---|---|---|
| *RvDTS1* | pESC-Ura vector | Forward | CAAGGAGAAAAAACCCCGGATCCATGGAAACAAAAGTCCATATGGTTCT |
| | | Reverse | CAACTTCTGTTCCATGTCGACTGCAGTAGCCAATCTCTTAAAGTTATTC |
| *RvDTR* | pESC-Ura::RvDTS1 vector | Forward | TCACTAAAGGGCGGCCGCACTAGTATGGACGCTGCTTCTGCTAA |
| | | Reverse | CCTTGTAATCCATCGATACTAGTCGGGTAGACTTCAAAGTGTTTTCGATATCA |
| *MsCYP72056* | pESC-Leu vector | Forward | TCACTAAAGGGCGGCCGCACTAGTATGGCTTCTTTCTCTGCTTTCCA |
| | | Reverse | CCTTGTAATCCATCGATACTAGTCGACATAAGGATGGAACCAATAAC |
| *AtATR2* | pESC-Leu::MsCYP72056 vector | Forward | CAAGGAGAAAAAACCCCGGATCCATGTCCTCTTCTTCTTCTTCGTCAAC |
| | | Reverse | CAACTTCTGTTCCATGTCGACGGATCCCCATACATCTCTAAGATATCTT |

Primers for *E. coli* vector construction (pOPIN vector). Cloning overhangs are underlined.

| Primer | Plasmid | Primer direction | Sequence |
|---|---|---|---|
| *CrSTR* | pOPINM | Forward | AAGTTCTGTTTCAGGGCCCGATGGCAAACTTTTCTGAATCTAAATCC |
| | | Reverse | ATGGTCTAGAAAGCTTTAGCTAGAAACATAAGAATTTCCCTTGTTAT |
| *CrSGD* | pOPINF | Forward | AAGTTCTGTTTCAGGGCCCGATGGGATCTAAAGATGATCAGTCCCT |
| | | Reverse | ATGGTCTAGAAAGCTTTAGTATTTTTGCTTCTTGACTAACTCAACTAGTTTAT |
| *CpDCS* | pOPINF | Forward | AAGTTCTGTTTCAGGGCCCGATGGCCGGAAAATCTCAAGAAGATGGG |
| | | Reverse | ATGGTCTAGAAAGCTTTAGCCTAGGTCTCTCGAACCAGCAGATTTTAAG |
| *MsEnolMT* | pOPINF | Forward | AAGTTCTGTTTCAGGGCCCGATGCAACCACAGAGAGGGAGAAAGAGAGA |
| | | Reverse | ATGGTCTAGAAAGCTTTAATCCTCTGCATTGCGTTTAAGG |

Primers for mutagenesis (3Ω1 vector). Mutated codons are in red.

| Primer | Plasmid | Primer direction | Sequence |
|---|---|---|---|
| RvDTRT65A | 3Ω1 | Forward | CTTGGCACTGCCAAATATCC |
| | | Reverse | GGATATTTGGCAGTGCCAAG |
| RvDTRS128A | 3Ω1 | Forward | CAGATGGAACTGCCTTTAGTGACG |
| | | Reverse | CGTCACTAAAGGCAGTTCCATCTG |
| RvDTRS130A | 3Ω1 | Forward | TCTTTTGCCGACGGAAACG |
| | | Reverse | CGTTTCCGTCGGCAAAAGA |
| RvDTRN133A | 3Ω1 | Forward | AGTGACGGAGCCGACCTATATTTC |
| | | Reverse | GAAATATAGGTCGGCTCCGTCACT |
| RvDTRD134A | 3Ω1 | Forward | AGTGACGGAAACGCCCTATATTTCTA |
| | | Reverse | TAGAAATATAGGGCGTTTCCGTCACT |
| RvDTRS302A | 3Ω1 | Forward | TTGCCAGTCGCCCCACT |
| | | Reverse | AGTGGGGCGACTGGCAA |
| RvDTRH99S | 3Ω1 | Forward | GGTGTTAGCAGCTACGTTCAGACA |
| | | Reverse | TGTCTGAACGTAGCTGCTAACACC |
| RvDTRH99A | 3Ω1 | Forward | GGTGTTAGCGCCTACGTTCAGACA |
| | | Reverse | TGTCTGAACGTAGGCGCTAACACC |
| RvDTRT123A | 3Ω1 | Forward | AACTTGATAGCCGCAGATGGA |
| | | Reverse | TCCATCTGCGGCTATCAAGTT |
| RvDTRT123G | 3Ω1 | Forward | AACTTGATAGGAGCAGATGGA |
| | | Reverse | TCCATCTGCTCCTATCAAGTT |
| RvDTR | 3Ω1 | Forward | TTTATGAATTTTGCAGCTCGATGGATGCAGCATCTGCAAA |
| | | Reverse | GACAACCACAACAAGCACCGCTAAGTAGATTTCAATGTATTCTCTATATCAATCAC |

Primers for sequencing.

| Plasmid | Primer direction | Sequence |
| --- | --- | --- |
| 3Ω1 | Forward | GATGAAAAAGCCCTAAAATTGGAG |
| | Reverse | ATTATTCACAAATGAGAAACAGAATG |
| GaL1 | Forward | CAACATTTTCGGTTTGTATTACTTC |
| | Reverse | CTTCGAGCGTCCCAAAAC |
| GaL10 | Forward | GAGCGACCTCATGCTATAC |
| | Reverse | GGATATGTATATGGATATGTATATGGTG |
| pOPINF | Forward | TAATACGACTCACTATAGGG |
| | Reverse | TAGCCAGAAGTCAGATGCT |
| pOPINM | Forward | GAAATCATGCCGAACATC |
| | Reverse | TAGCCAGAAGTCAGATGCT |

**Supplementary Table 4. Previously characterized MDRs used to construct phylogenetic tree in Supplementary Fig. 9.**

| Gene Name | Genbank accession |
|---|---|
| *Rauwolfia serpentina* vomilenine reductase 2 (VR2) | KT369740.1 |
| *Tabernanthe iboga* dihydroprecondylocarpine acetate synthase 1 (DPAS1) | MK840855.1 |
| *Tabernanthe iboga* dihydroprecondylocarpine acetate synthase 2 (DPAS2) | MK840856.1 |
| *Catharanthus roseus* dihydroprecondylocarpine acetate synthase 1 (DPAS) | KU865331.1 |
| *Strychnos speciosa* Wieland-Gumlich aldehyde synthase (WS) | OM304303.1 |
| *Strychnos nux-vomica* Wieland-Gumlich aldehyde synthase (WS) | OM304294.1 |
| *Catharanthus roseus* Geissoschizine synthase (GS) | MF770507.1 |
| *Cinchona pubescens* dihydrocorinantheine aldehyde synthase (DCS) | MW456554 |
| *Catharanthus roseus* tetrahydroalstonine synthase (THAS) | KM524258.1 |
| *Catharanthus roseus* heteroyohimbine synthase (HYS) | KU865325.1 |
| *Catharanthus roseus* tabersonine 3- reductase (T3R) | KP122966.1 |
| *Rauwolfia tetraphylla* vomilenine reductase 2 (VR2) | KT369741.1 |
